# Human *ribomes* reveal DNA-embedded ribonucleotides as a new type of epigenetic mark

**DOI:** 10.1101/2025.06.27.661996

**Authors:** Deepali L. Kundnani, Taehwan Yang, Tejasvi Channagiri, Penghao Xu, Yeunsoo Lee, Mo Sun, Francisco Martinez-Figueroa, Supreet Randhawa, Ashlesha Gogate, Youngkyu Jeon, Stefania Marsili, Gary Newnam, Yilin Lu, Vivian S. Park, Sijia Tao, Justin A. Ling, Raymond F. Schinazi, Zachary F. Pursell, Abdulmelik Mohammed, Patricia L. Opresko, Bret D. Freudenthal, Baek Kim, Soojin V. Yi, Nataša Jonoska, Francesca Storici

## Abstract

Ribonucleoside monophosphates (rNMPs) are abundant in DNA, but their distribution and function in human nuclear genomes remain unknown. Here, we mapped nearly one million rNMPs per genome across diverse human cell types, defining a nuclear “*ribome*” with non-random distribution patterns. rNMPs are enriched in C/G-rich sequences, epigenetically marked regions, and telomeres. Conserved ribonucleotide-enriched zones (REZs) overlap with CpG islands and R-loops. rNMP concentration near transcription start sites (TSSs) correlates positively with gene expression. Wild-type cells display a broader gene-expression range than ribonuclease H2A (RNH2A) knockouts, in which loss of rNMP cleavage causes pronounced retention of embedded rG and strand-biased rC near TSSs, both increasing with gene expression. These findings establish DNA-embedded rNMPs as a novel epigenetic mark that modulates human gene expression.

## Main text

Ribonucleoside monophosphates (rNMPs) are the most abundant non-standard nucleotides embedded in genomic DNA (*1–3*). Although DNA polymerases are selective against ribonucleoside triphosphates (rNTPs), they nonetheless incorporate them as rNMPs during replication and repair, with incorporation frequency varying across polymerases and species (*1–3*). For instance, in vitro yeast replicative polymerases Pols α, *δ*, and ε incorporate rNTPs at rates of ~1 per 650 to 5,000 dNTPs (*4, 5*), while human Pol *δ* incorporates ~1 per 2,000 deoxyribonucleoside triphosphates (dNTPs) (*6*). These rates make rNMPs more prevalent than other common forms of DNA damage such as depurination or oxidation (*7*). The embedded rNMPs, due to their 2′-hydroxyl group, can alter DNA structure and increase susceptibility to breaks, mutations, and disruption of replication and transcription (*3, 7–12*). Many rNMPs are removed by ribonuclease (RNase) H2-mediated ribonucleotide-excision repair (RER) (*5*), and when RER fails, topoisomerase 1 (Top1) can cleave at rNMPs, triggering error-prone repair (*1, 13–15*). Defects in RNase H2 cause rNMP accumulation and genome instability and are linked to disorders including Aicardi-Goutières syndrome and cancer (*16–24*). While rNMPs have been studied substantially in yeast (*25–34*) and human mitochondria (*21, 35–37*), their distribution and functional impact in human nuclear DNA remains unknown. Prior work using rNMP-hyper-incorporating polymerase mutants mapped DNA polymerase usage in human cancer cell lines (*38*), but did not provide a genome-wide map of endogenously embedded rNMPs in human chromosomes. Studies in model systems suggest that rNMP incorporation is non-random and in part influenced by nucleotide pools, polymerase identity, and sequence context (*31, 32, 34, 37, 39, 40*). Whether embedded rNMPs in human chromosomal DNA have regulatory roles beyond genome destabilization remains an open question. To fill this gap, we systematically mapped rNMPs in nuclear DNA across diverse human cell types, revealing a structured nuclear “*ribome*” and uncovering DNA-embedded rNMPs as a new type of epigenetic mark.

## Results

### rNMPs are abundant and non-randomly distributed in human chromosomal DNA

To study rNMPs in human genomic DNA across diverse biological contexts, we selected multiple human cell types representing different developmental states, tissue lineages, and DNA repair capacities. These included CD4+T lymphocytes (differentiated immune cells), human embryonic stem cells (hESC-H9, pluripotent), and human embryonic kidney (HEK293T) cells in both wild-type and RNH2A-knockout (KO) backgrounds to assess the role of RNase H2-mediated rNMP repair. To confirm the presence of rNMPs across all cell types, we treated genomic DNA with *Escherichia coli* RNase HII, which nicks at embedded rNMPs, and analyzed the samples by denaturing gel electrophoresis (see Methods). Agarose gel electrophoresis revealed shorter fragment lengths in RNase HII-treated samples, particularly in RNH2A-KO cells, indicating more embedded rNMPs (**fig. S1A-D**). Fragment length shortening was also observed in wild-type cells, confirming rNMP presence across all cell types. Simulations estimated ~1 million rNMPs per diploid genome (see Methods, **table S1**).

Using ribose-seq (*27*) and Ribose-Map (*41*), we generated maps of embedded rNMPs across all cell types, defining this genomic landscape as the “*ribome*.” Ribose-seq libraries were prepared from enzymatically fragmented DNA using multiple restriction enzyme sets (RE1–RE3) and dsDNA Fragmentase (F) (see **table S2**) to ensure broad genome coverage. In total, we generated 26 libraries across all cell types (**table S2** and **S3**). Sequencing reads were aligned to GRCh38, and single-nucleotide rNMP coordinates were extracted using Ribose-Map, which excludes low-confidence regions such as centromeres and telomeres (see Methods). To assess genomic distribution, we compared biological rNMP libraries to random-control libraries matched for number of rNMP sites and frequencies of rNMPs per site (**Fig. 1A**). Observed rNMPs showed significant non-random distributions across autosomes (Mann-Whitney *U* and Kolmogorov-Smirnov tests, *p* < 0.05; **Fig. 1A, table S4**), especially in RNH2A-KO cells. Libraries with higher rNMP counts yielded greater divergence from random datasets and stronger statistical significance.

**Figure. 1.**
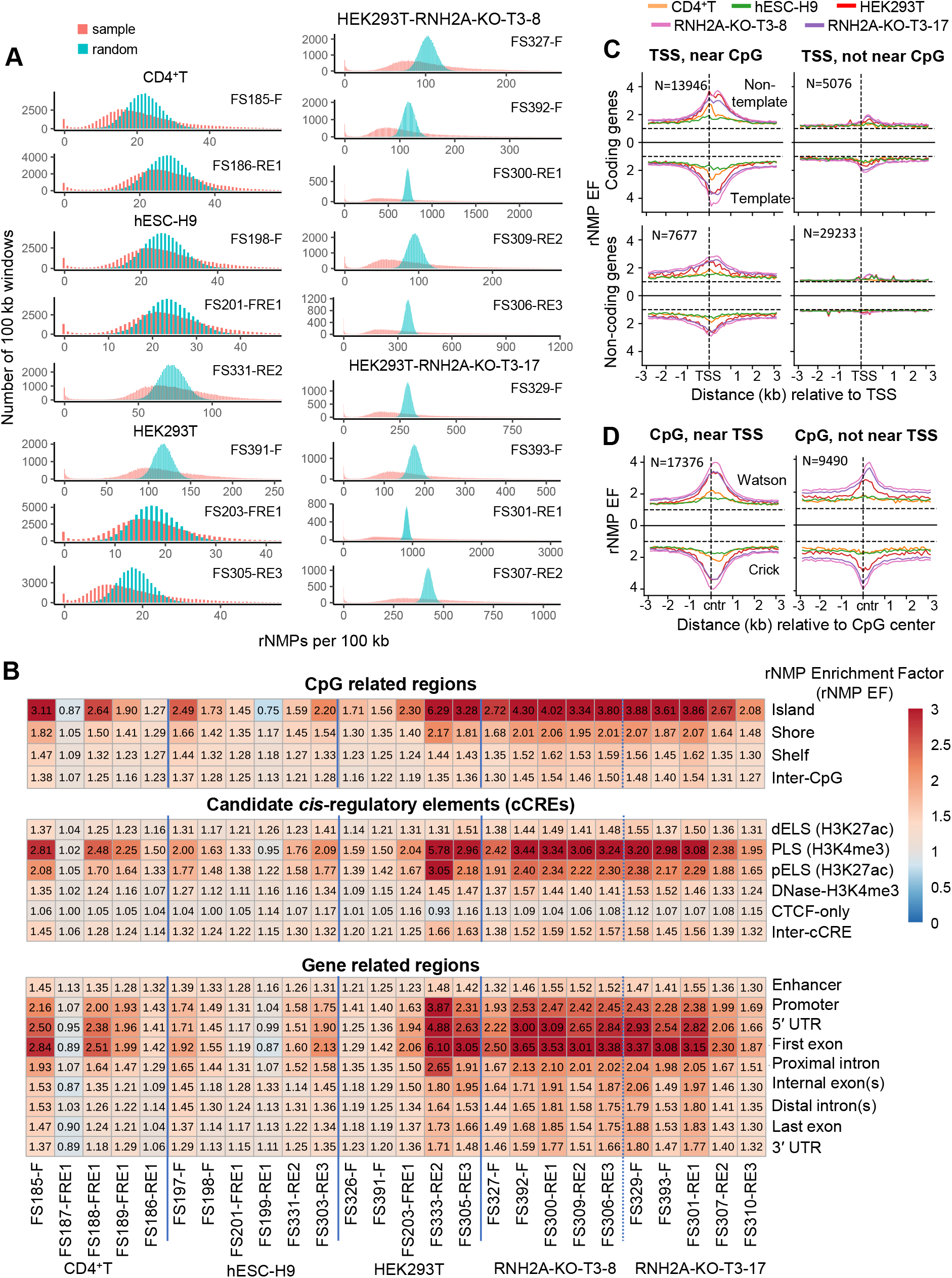
rNMP incorporation in human nuclear DNA is not random and is associated with CpG islands and promoters. (**A**) Distribution of biological (sample, orange) rNMP counts and artificially generated rNMPs (random, teal) for each library, in 100 kb discrete bins across autosomal chromosomes (chr 1 – chr 22). *X*-axis represents the count of rNMPs in 100 kb bins and *y*-axis represents the number of 100 kb bins with respective rNMP counts in 100 kb bins. Libraries with greater than 1 million rNMPs are presented in this figure. Mann-Whitney *U* and Kolmogorov-Smirnov tests between biological and random rNMP sets showed significant difference (*p*-values < 0.05, **table S4**). (**B**) Heatmaps showing rNMP enrichment mean in CpG related regions, cCREs, and gene related regions for protein coding genes. Enrichment (rNMP EF > 1) is shown in red, depletion (0 < rNMP EF < 1) in blue, and neutral enrichment (rNMP EF = 1) in white, based on rNMP frequency per base in autosomes (chr1 to chr22). Significant enrichment in wild-type and/or RNH2A-KO cells was checked by Mann-Whitney *U*-test (*p* < 0.05; **table S5**). (**C**) Line plots showing mean rNMP Enrichment Factor (EF, *y*-axis) in 100-bp bins, ± 3 kb (*x*-axis) around transcription sites (TSSs) of coding and non-coding genes, either near or not near CpG islands, on non-template (top) and template (bottom) strands. Cell types are color-coded: CD4^+^T (orange), hESC-H9 (green), HEK293T (red), HEK293T-RNH2A-KO-T3-8 (pink), and HEK293T-RNH2A-KO-T3-17 (purple), with strands separated by a solid black line. Dotted lines indicate EF = 1. (**D**) Line plots showing mean rNMP EF (*y*-axis) in 100 bp bins ±3 kb (*x*-axis) around CpG islands that are either near or not near TSSs, on the Watson (top) and Crick (bottom) strands. Cell types are shown in corresponding colors, with strands separated by a solid black line. Dotted lines indicate EF = 1. The number of genes (N) is indicated for each group. For each gene, rNMP EF was calculated and averaged across gene groups, and then across libraries of the same cell type (see Methods).

### Embedded rNMPs are enriched at CpG islands, promoters, and transcription start sites (TSSs)

To explore the non-random distribution of embedded rNMPs, we analyzed their mono- and dinucleotide context and genomic localization. Pearson correlation between rNMP and deoxyribonucleoside monophosphate (dNMP) counts in varying genomic bin sizes showed strong positive correlation with G and C mononucleotides, and with GC-rich dinucleotides (CC, CG, GC, GG), especially in RNH2A-KO cells and larger bins (**fig. S3A–D**). rNMP enrichment also correlated with regions exhibiting positive A–T skew (more A than T), but not with G–C skew, suggesting a preference for sequence composition (**fig. S3E**).

Sequence analysis ±50 bp around rNMP sites confirmed the overrepresentation of C and G bases and GC-rich dinucleotides, particularly CG, in both wild-type and RNH2A-KO libraries (**fig. S2**), implicating CpG islands as rNMP hotspots. Using three annotation sets: i) CpG-related regions with island, shore, shelf, and inter-CpG) (*42*), ii) candidate *cis*-regulatory elements (cCREs) with distal enhancer-like signatures (dELS) (H3K27ac), promoter-like signatures (PLS) (H3K4me3), proximal enhancer-like signatures (pELS) (H3K27ac), DNase-H3K4me3, CCCTC-binding factor sites (CTCF-only), and inter-cCRE (*43*), and iii) gene structure with enhancer, promoter, 5′ untranslated region (UTR), first exon, proximal intron, internal exon(s), distal intron(s), last exon, and 3′ UTR (*42*), we computed the relative rNMP enrichment factor (rNMP EF) in these regions (see Methods). CpG islands consistently showed the highest rNMP EF across all libraries (**Fig. 1B**). Promoter-like signatures (PLS), 5′ UTRs, and first exons also displayed elevated enrichment (**Fig. 1B**). These trends held across cell types, with RNH2A-KO cells showing higher rNMP EF. Enrichment differences between annotated and reference regions were statistically significant particularly in RNH2A-KO cells (Mann-Whitney *U* test *p* < 0.05; **table S5**).

We further examined rNMP localization near TSSs and CpG islands. Strong rNMP enrichment occurred within ~1 kb of TSSs that are within 3 kb of CpG island centers (**Fig. 1C**, left), ~65% of which are within 3 kb of TSSs (*44*). Enrichment was reduced when TSSs were >3 kb apart from CpG islands (**Fig. 1C**, right). Interestingly, distal CpG islands >3 kb from TSSs still showed rNMP enrichment, particularly at edges (>1kb around the CpG-island centers) (**Fig. 1D**). These distal CpG islands are often overlapping with introns, potential regulatory elements influencing gene expression (*45*), and dELS regions marked by H3K27ac, which associate with active promoters and TSSs via chromatin looping (*46*) (**fig. S4**). Together, these findings suggest that CpG islands promote rNMP incorporation, particularly near TSSs or within physically linked regulatory domains.

### Ribonucleotide-enriched zones (REZs) are strand-symmetric and colocalize with CpG islands and R-loops

Genome-wide analysis revealed discrete ribonucleotide-enriched zones (REZs), defined as five consecutive 0.5 Mb bins with rNMP EF >1.2 in ≥80% of libraries, across all cell types. REZs displayed strong strand symmetry and were enriched in CpG island-dense regions, as shown in karyotype plots (**Fig. 2**), with higher rNMP EF in RNH2A-KO cells, particularly on chromosome 19 (**Fig. 2**, zoom in).

**Figure. 2.**
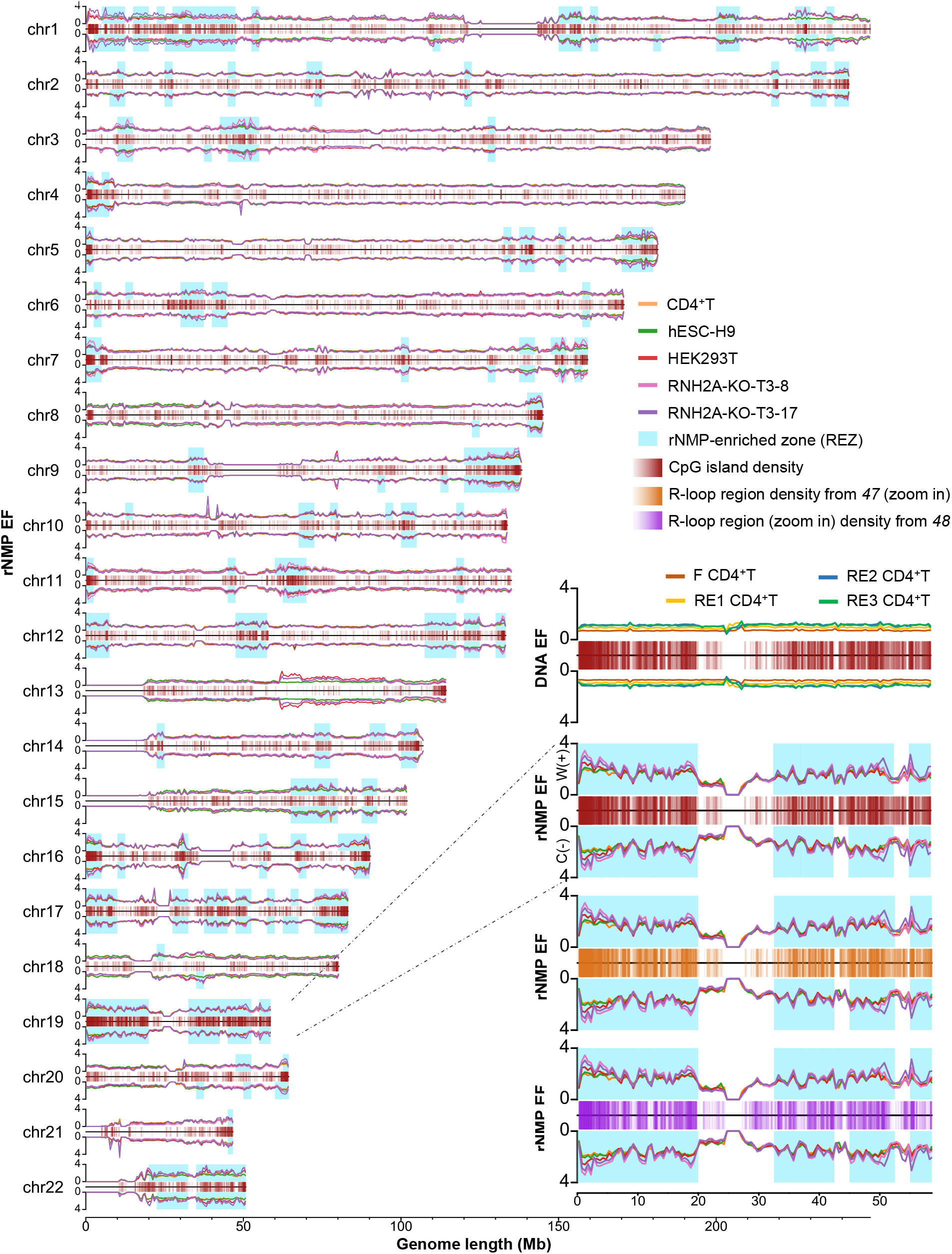
rNMP enriched zones (REZs) colocalize with CpG islands and R-loop regions across the human genome. (**A**) Karyotype-style line plots showing rNMP enrichment factor (EF, *y* axis) on the Watson (W, +) and Crick (C, −) strands across genomic coordinates (*x-*axis, in megabases, Mb) for chr1 – chr22. Cell types are color-coded: CD4^+^T (orange), hESC-H9 (green), HEK293T (red), HEK293T-RNH2A-KO-T3-8 (pink), and HEK293T-RNH2A-KO-T3-17 (purple). CpG islands are shown in the center (transparent red bars: regions with high CpG island density appear as darker red, while regions with lower density appear lighter) and REZs identified across all cell types for both strands are highlighted in blue. (**B**) Zoom-in on chr19: top plot, EF of DNA-seq read coverage in CD4+T cells fragmented using the same enzyme combinations as ribose-seq libraries, dsDNA Fragmentase (F) (red), RE1 (yellow), RE2 (blue), and RE3 (green); CpG islands in the center (transparent red bars); second plot: rNMP enrichment with CpG islands centered; third plot, rNMP enrichment with R-loop regions from (*47*) centered (transparent orange bars); bottom plot, rNMP enrichment centered on R-loop peaks from RIAN-seq data (*48*) (transparent purple bars). Full-chromosome views are shown in **fig. S5**.

To rule out artifacts from DNA fragmentation, we computed DNA enrichment factors (DNA EF) in the DNA extracted from CD4^+^T cells that were fragmented by the four enzyme sets used in ribose-seq preparations. DNA EF showed uniform coverage and no colocalization with CpG islands, confirming that REZs are not due to technical bias (**Fig. 2, fig. S5A**).

Importantly, REZs also colocalized with R-loop regions mapped by two independent studies (*47, 48*) (**fig. S5B**,**C** and **Fig. 2**, zoom in), further supporting their association with transcriptionally active, GC-rich regions. R-loops, which form when nascent RNA hybridizes to the template DNA strand, frequently occur at CpG islands and are implicated in transcriptional regulation (*47*).

### rNMP enrichment near gene TSSs correlates with gene expression levels

Analysis of rNMP EF in 100-bp bins around transcription start sites (TSSs) revealed strong enrichment within 1 kb of TSSs for coding genes across all tested cell types, with the highest levels in HEK293T and lowest in hESC-H9 cells (**Fig. 3A**). No enrichment was observed near transcription termination sites (TTSs). rNMP EF near coding TSSs reached up to a factor of 3, and up to a factor of 1.75 for non-coding genes (**fig. S6A**).

**Figure. 3.**
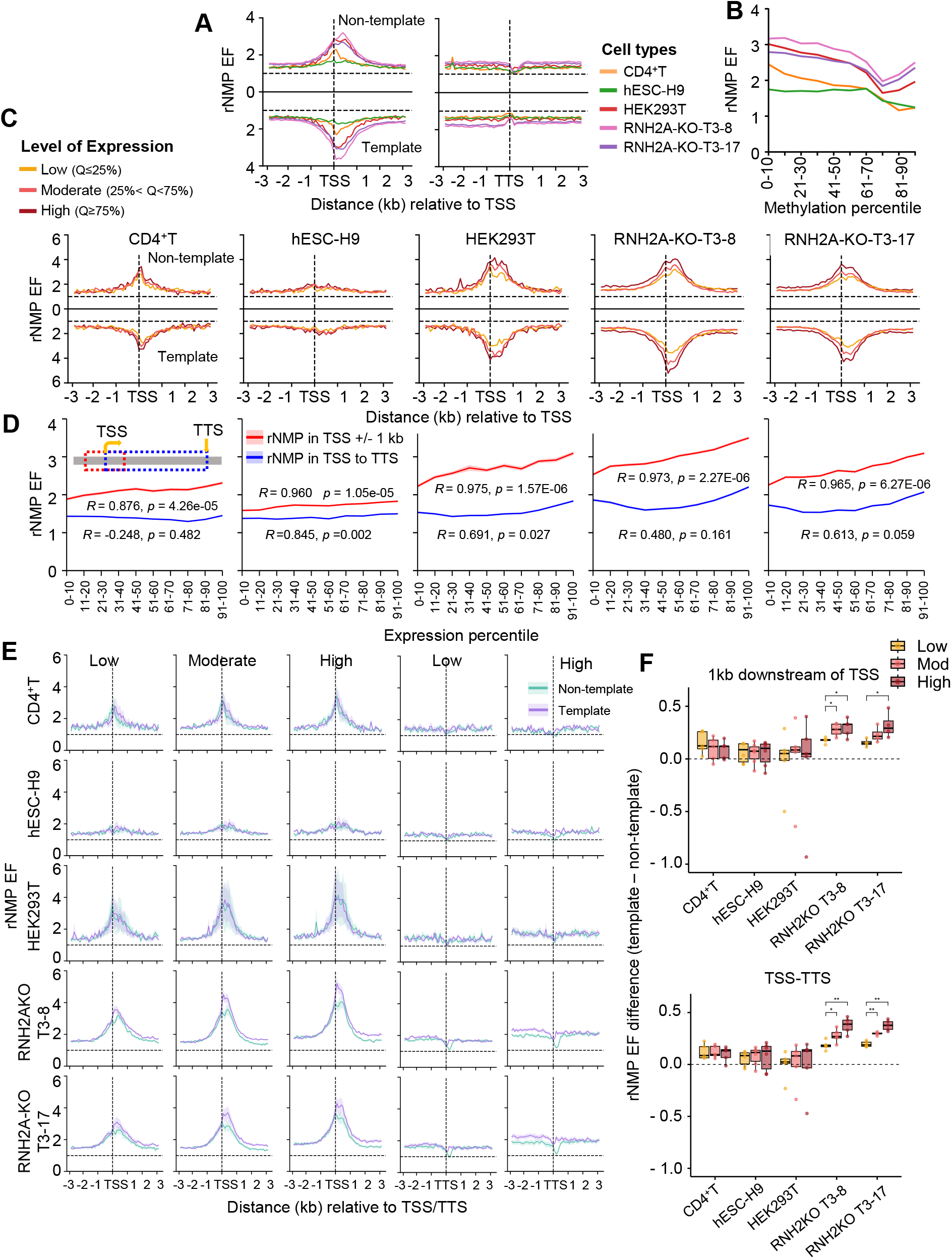
rNMP enrichment around TSSs correlates with elevated gene expression. (**A**) Line plots showing mean rNMP EF (*y*-axis) in 100-bp bins, ±3 kb (*x*-axis) around transcription TSSs, on the non-template (top) and template (bottom) strands. Cell types are color-coded: CD4^+^T (orange), hESC-H9 (green), HEK293T (red), HEK293T-RNH2A-KO-T3-8 (pink), and HEK293T-RNH2A-KO-T3-17 (purple). (**B**) Mean rNMP EF (*y*-axis) within ±1 kb of TSSs plotted against increasing DNA methylation levels, grouped by deciles (*x*-axis, left to right) for each cell type. Each group contains an equal number of TSSs. Pearson’s *R* and *p*-value between rNMP EF and methylation percentage are provided in **table S6**. (**C**) Line plots showing mean rNMP EF (*y*-axis) for 100 bp bins, ±3 kb (*x*-axis) around TSSs, grouped by gene expression levels: low (≤25th percentile, yellow), moderate (25th–75th percentile, orange), and high (≥75th percentile, red). Non-template (top) and template (bottom) strand signals are separated by a solid black line. Horizontal and vertical dotted lines indicate EF = 1 and TSS position, respectively. (**D**) Mean rNMP EF (*y*-axis) for regions ±1 kb around TSS (red) and across gene bodies (TSS to TTS, blue) plotted against increasing gene expression levels, grouped into deciles (*x*-axis, left to right) for each cell type. Each group contains an equal number of TSSs. Pearson’s *R* and *p*-value are shown in the figure and listed in **table S7**, based on mean rNMP EF and mean expression log_2_(TPM) values for rNMPs ± 1 kb around TSSs and across gene bodies. (**E**) Line plots showing mean and shaded regions indicating standard deviation of rNMP EF (*y*-axis) in 100 bp bins ± 3 kb (*x*-axis) around TSSs of previously defined low-, moderate-, and high-expressed genes, on the non-template (green) and template (purple) strands. Horizontal and vertical dotted lines indicate EF = 1 and TSS position, respectively. Similar distributions for low and highly expressed TTSs are also shared. (**F**) Box plots showing the difference in rNMP EF between the template and non-template strands for low (yellow), moderate (orange), and high (red) expressed genes across cell types. Each dot represents a library for the corresponding cell type. The top plot shows strand-specific differences within 1 kb downstream of TSSs; the bottom plot shows differences across the full gene body (TSS to TTS). Comparisons using two-tailed Mann–Whitney *U*-test are marked with asterisks (*). Significance levels: * (0.05 ≥ *p* > 0.01), ** (0.01 ≥ *p* > 0.001), *** (*p <* 0.001), see **table S8**.

To examine epigenetic context, we assessed DNA methylation via whole-genome bisulfite sequencing after mechanical shearing (CD4^+^T) (see Methods) and public datasets (hESC-H9, HEK293T, see Methods). Across all cell types, rNMP EF within 1 kb of TSSs negatively correlated with DNA methylation levels in these regions (**Fig. 3B, fig. S7A**,**B, table S6**), suggesting that rNMP enrichment may mark transcriptionally active, hypomethylated regions. Interestingly, an rNMP EF rebound is observed in highly methylated regions, particularly for HEK293T-RNH2A-KO cells (**Fig. 3B, fig. S7B**).

To evaluate whether rNMP enrichment correlates with gene expression, we then analyzed RNA-seq data (HEK293T wild-type and RNH2A-KO libraries constructed for this study, and from public datasets for CD4^+^T and hESC-H9, see Methods). Genes were grouped into low, moderate, and high expression categories (see Methods). In all cell types, higher expression levels were associated with greater rNMP EF near TSSs (**Fig. 3C**), with the strongest correlation observed within 1 kb of the TSS (**Fig. 3D, fig. S6B, table S7**). By contrast, a high and significant correlation between gene body rNMP levels and expression was observed in hESC-H9 cells, although a lower but still significant correlation was also seen in wild-type HEK293T cells. Notably, rNMP enrichment peaked immediately downstream of the TSS on both strands, in both coding and non-coding genes (**Fig. 3A,C, fig. S6A, table S7**).

### Strand-biased rNMP enrichment emerges in RNH2A-KO cells and increases with gene expression

To assess strand bias in rNMP enrichment, we compared rNMP EF on template vs. non-template strands within 3 kb of coding gene TSSs across expression levels. Unlike RNase H2 wild-type cells, RNH2A-KO cells displayed a clear bias, higher rNMP EF on the template strand, especially in highly expressed genes (**Fig. 3E**, left, and **fig. S6C**,**D**). This bias was most pronounced just downstream of the TSS and extended into the gene body (**Fig. 3F**, top, **table S8**). No significant bias was observed at and near the TSS in wild-type cells and upstream of the TSS in RNH2A-KO cells. Instead, similar strand-biased enrichment also appeared at and upstream of TTSs, again restricted to RNH2A-KO cells (**Fig. 3E**, right). Indeed, in RNH2A-KO cells, we observed an increasing difference in rNMP EF between the template and non-template strands within gene bodies (TSS to TTS) as gene expression levels rose (**Fig. 3F**, bottom, **table S8**). These results indicate that RNase H2 inactivation leads to strong, expression-dependent template-strand enrichment of rNMPs, beginning at the TSS and persisting through the transcription unit.

### rG enrichment downstream of TSS correlates with elevated gene expression and reveals strand-specific patterns in RNH2A-KO cells

To examine rNMP composition near TSSs, we analyzed rA, rC, rG, and rU levels from TSS to 1 kb downstream, where rNMP enrichment peaks, in RNase H2 wild-type and RNH2A-KO cells, which primarily contain rNMPs cleaved by RNase H2 and removed via RER in wild-type cells (*1*). We examined how differences in rNMP composition and strand-specific distribution relate to gene expression levels of wild-type cells. We computed total counts, average EF, and rNMP percentages from the TSS to 1 kb downstream on both strands, as well as separately on template and non-template strands, for each gene in all ribose-seq libraries. In wild-type libraries, all rNMP types showed positive correlations with gene expression, especially rC and rG, while their relative proportions (percentages) remained mostly stable (**Fig. 4A, fig. S8A, table S9**). Minimal correlation was seen upstream (2-3 kb) or further downstream (4-5 kb) of TSS (**fig. S8B**,**C, table S9**). In RNH2A-KO libraries, rG showed significantly increased counts, EF, and percentage near the TSS, and rG levels in RNH2A-KO libraries correlated more strongly with wild-type gene expression than rG levels in wild-type libraries. rG exhibited a larger count and EF, and a steeper slope than rC (**Fig. 4A, fig. S8A, table S9**). This pattern was less evident in regions further from the TSS (**fig. S8B**,**C, table S9**).

**Figure. 4.**
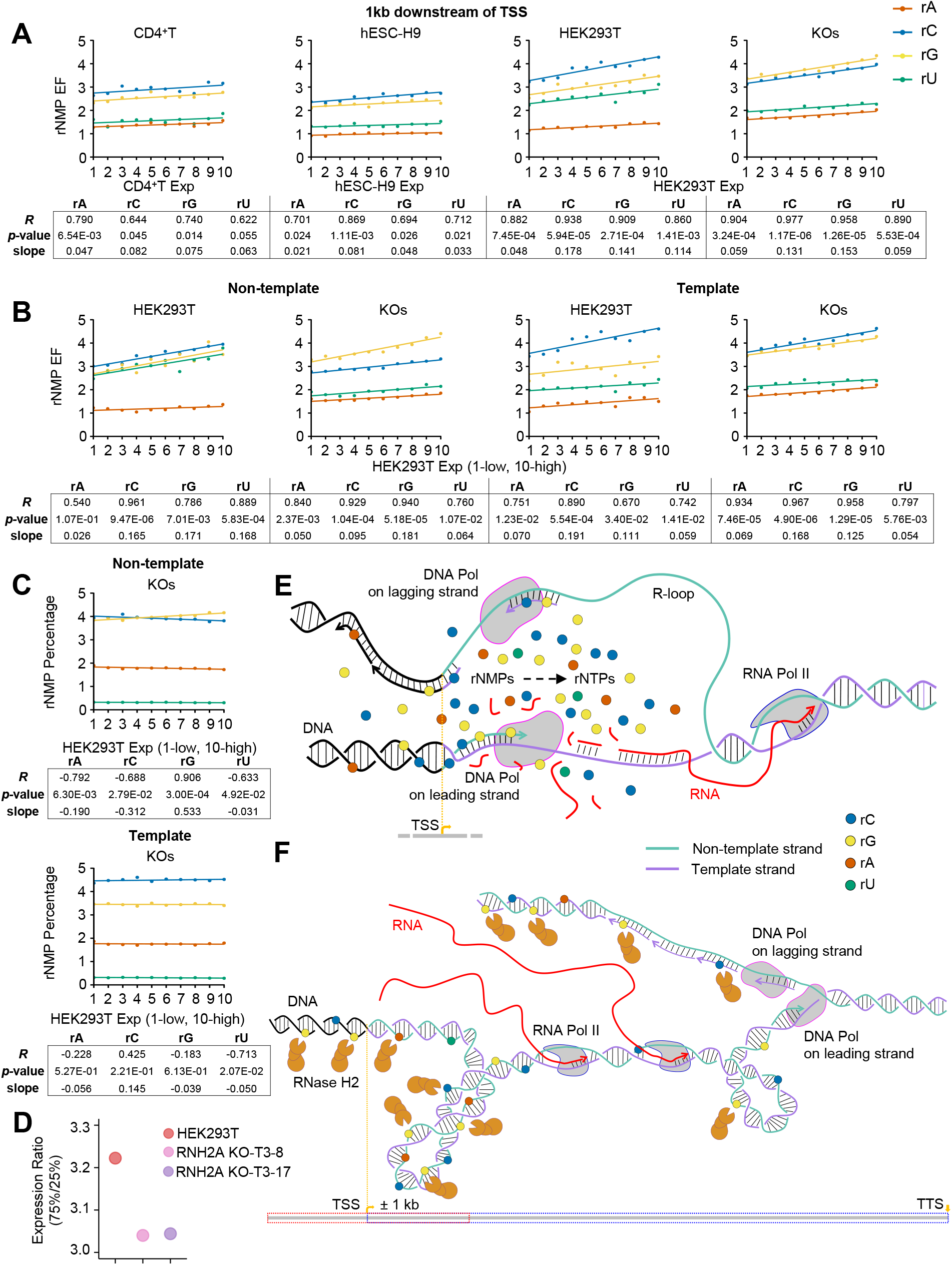
Enrichment of individual rNMPs with gene expression. (**A**) Mean rNMP EF (*y*-axis) within 1 kb downstream of TSS for each rNMP: rA (red), rC (blue), rG (yellow), and rU (green) plotted across increasing expression levels (*x*-axis, in 10 percentile groups, left, low to right, high) for each cell type. Each group contains an equal number of TSSs. Linear regression lines for each rNMP are shown, with Pearson’s *R, p*-value, and linear regression slope summarized in the table below for each cell type. (**B**) As in (A) but showing trends for each rNMP within 1 kb downstream of TSSs for HEK293T wild-type and RNH2A-KO cells, based on HEK293T expression, shown separately for the non-template and template strands. (**C**) Percentage of each rNMP base (*y*-axis) out of total rNMPs within 1 kb downstream of TSSs on the non-template and template strands, across expression percentile groups (*x*-axis) for RNH2A-KO cells, relative to wild-type HEK293T expression. Linear regression lines for each rNMP are shown, with Pearson’s *R, p*-value, and linear regression slope summarized in the table below. See also **figs. S8**,**9** and **table S9**. (**D**) Dot plots showing expression range ratios, defined as the 75th percentile (Q3) divided by the 25th percentile (Q1), used to group genes into low, moderate, and high expression categories for each cell type. RNA-seq data are shown for HEK293T (red), HEK293T-RNH2A-KO-T3-8 (pink), and HEK293T-RNH2A-KO-T3-17 (purple) cells. (**E**) Model for rNMP incorporation near TSSs. Enrichment of rC and rG may reflect local use of recycled rNMPs from R-loop degradation at C/G-rich regions, with R-loop formation potentially driven by negative supercoiling upstream of RNA polymerases in highly transcribed genes. (**F**) Model for the epigenetic role of rNMPs near TSSs as cleavage sites for RNase H2 (rGs) in wild-type cells, relieving torsional stress during replication and transcription, particularly on the leading strand, to facilitate rapid and efficient transcription restart.

Strand-specific analysis revealed that in wild-type cells, rC, rG, and rU were similarly enriched on the non-template strand, while rC dominated on the template strand. In RNH2A-KO cells, rG enrichment rose markedly on both strands, nearly matching rC levels on the template strand, while rC and rU enrichment markedly decreased on the non-template stand (**Fig. 4B, fig. S9A, table S9**). Notably, rG percentage increased with gene expression only in RNH2A-KO libraries and only on the non-template strand (**Fig. 4C, fig. S9A, table S9**). This strand bias was not observed upstream or further downstream of TSS (**fig. S9B**,**C, table S9**).

Finally, transcript interquartile (IQ) ratios used to define highly and low expressed genes were higher in RNase H2 wild-type than in KO cells (**Fig. 4D**), suggesting that RNase H2 activity may broaden the dynamic range of gene expression.

### Sequence preferences shape rNMP incorporation in human nuclear DNA

By normalizing rNMP base composition to corresponding dNMPs, we observed distinct preferences between wild-type and RNH2A-KO cells (**Fig. 5A–C, fig. S10A**). rC incorporation was consistently high across all cell types, while rA was enriched in CD4^+^T cells and rG showed specific enrichment in RNH2A-KO cells. rU was least incorporated, especially in RNH2A-KO cells. These patterns suggest that rNMP incorporation is influenced by factors beyond nucleotide availability, as they do not mirror the rNTP/dNTP ratios measured in whole-cell extracts (*37*) (**fig. S10B**).

**Figure. 5.**
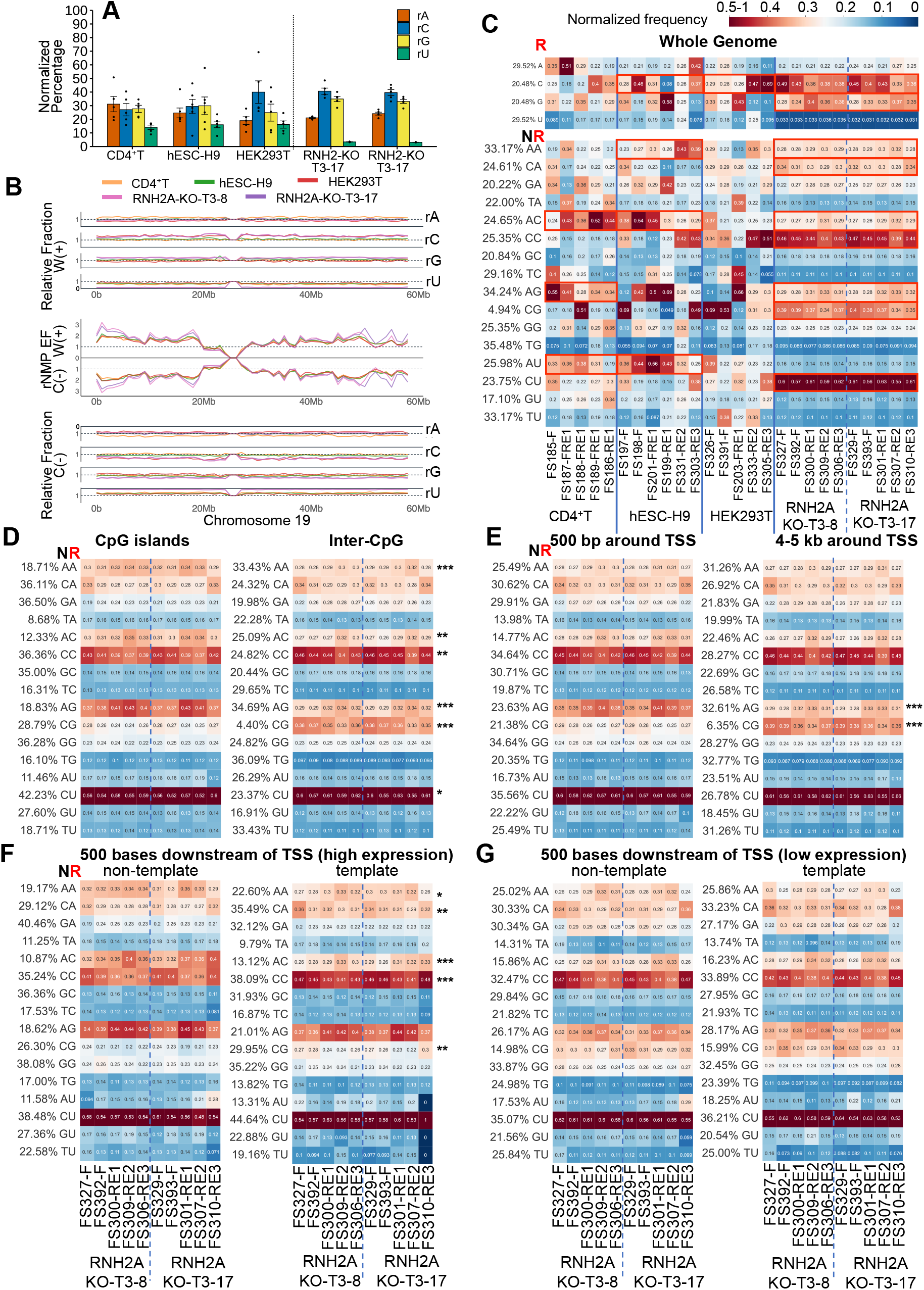
rNMP preference for upstream dA in CpG islands and near TSSs of highly expressed genes. (**A**) Bar graphs showing the mean and standard error of normalized percentages of rNMPs rA (orange), rC (blue), rG (yellow), and rU (green), normalized to the background genomic frequency of dNMPs, for libraries from the indicated cell type. (**B**) Line plots showing the distribution of rNMP EF for the Watson and Crick strands (center panel), with the relative fraction for each rNMP on the Watson strand (top panel) and Crick strand (bottom panel) plotted across 1 Mb of chromosome 19 (*x*-axis, fraction on *y*-axis), relative to the dNMP background. (**C**) Heatmaps displaying the normalized frequency for rNMPs (rA, rC, rG, and rU) and NR dinucleotides (each rNMP preceded by dA, dC, dG, or dT) across all ribose-seq libraries (columns). Libraries of different genotypes are separated by thick vertical blue lines; the two RNH2A-KO mutants are separated by a dashed vertical blue line. Genomic frequencies for each mono- and dinucleotide are shown on the left. The legend (top right) indicates frequency representation: white for 0.25; light red to red for 0.25 to 0.5–1; dark blue to light blue for 0–0.25. The dinucleotide heatmaps highlight dNMP–rNMP preferences for RNH2A-KO cells in: (**D**) CpG islands and inter-CpG regions, (**E**) 0–500 bp and 4–5 kb around TSS, (**F**) non-template and template strands, 500 bases downstream of TSSs of highly expressed genes (>75th percentile), and (**G**) non-template and template strands, 500 bases downstream of TSSs of low expressed genes (<25th percentile). Significant genome-wide nucleotide preferences (one-tailed Mann–Whitney *U*-test) are outlined with red rectangles; see **tables S10** and **S14** for data in (C-E) and (F,G), respectively. Comparisons between RNH2A-KO libraries (two-tailed Mann–Whitney *U*-test) are marked with asterisks (*). Significance levels: * (0.05 ≥ *p* > 0.01), ** (0.01 ≥ *p* > 0.001), *** (*p <* 0.001), see **table S13** for data in (D-G).

Sequence analysis around rNMP sites revealed stronger upstream dinucleotide preferences in RNH2A-KO cells, particularly for dA and dC preceding rA, dA and especially dC preceding both rC and rG, and for dC upstream of rU (**Fig. 5C, table S10**). Downstream, rU in wild-type cells was followed by dG, while in RNH2A-KO cells, rA and rG had preferences for dG, and rU for dC (**fig. S10C, table S11**). Trinucleotide analyses showed preferences for rUGG in wild-type and for ACrC, TCrC, and other pyrimidine-rich motifs in RNH2A-KO cells (**fig. S11** and **table S12**).

In RNH2A-KO cells, rNMP preferences varied by genomic context. In CpG islands, ArA, ArC, ArG motifs were preferred, with significant reduction of CrC and CrG compared to inter-CpG regions (**Fig. 5D, tables S10, S13**). Similarly, ArG was more frequent and CrG was reduced near TSSs compared to distal regions (**Fig. 5E, tables S10, S13**). Strand-specific patterns were evident 500 bp downstream of the TSS: ArA and ArC preferred on the non-template strand; CrA, CrC, and CrG on the template strand (**Fig. 5F**; **tables S14, S13**). These strand biases were absent in low expressed genes (**Fig. 5G, tables S14, S13**). Additional analyses showed rA decrease with accompanying rC increment in inter-CpG regions and on the template strand downstream of TSSs in highly expressed genes (**fig. S12, table S13**).

Together, these results reveal that rNMP incorporation in RNH2A-KO cells is shaped by sequence, transcriptional activity, strand identity, and genomic features such as CpG islands and proximity to TSS.

### In human telomers rNMPs show more frequent incorporation exhibiting distinct patterns

We analyzed rNMP incorporation in human telomeres by extracting ribose-seq reads containing >3 consecutive telomeric repeats (TTAGGG/AATCCC) from dsDNA Fragmentase (F)-fragmented libraries, thereby overcoming limitations in repeat-region alignment (see Methods). Reads with ≤3 repeats were assigned to telomeric-like units in the genome. rNMPs were abundantly present in telomeres across all cell types (**table S15**). In wild-type RNase H2 cells, the G-strand of telomeres (telomeric repeats) showed elevated rA incorporation, in contrast to telomeric-like units in the genome, which showed rG dominance at G positions (**Fig. 6A**). This strand-specific rA enrichment was consistent across CD4^+^T, hESC-H9, and HEK293T libraries. No such G-strand difference was observed in RNH2A-KO cells, where both telomeric and genomic units displayed dominant rG incorporation (**Fig. 6A**). Dinucleotide heatmaps (NR format) further revealed a distinct TrA pattern on the telomeric G-strand in wild-type cells, while ArG was more common in genomic telomeric-like units (**Fig. 6B**). These differences were absent in RNH2A-KO libraries, which showed uniform ArG and GrG patterns on the G-strand and CrC on the C-strand across both regions (**Fig. 6B**). The rNMP EF in telomeres was consistently high, averaging >5 in wild-type and >2 in RNH2A-KO libraries, based on rNMP counts and estimated telomere lengths (**table S15** and **Fig. 6C**). These findings indicate elevated rNMP incorporation and distinct G-strand preferences in telomeres, likely reflecting both increased incorporation and reduced removal in wild-type cells.

**Figure. 6.**
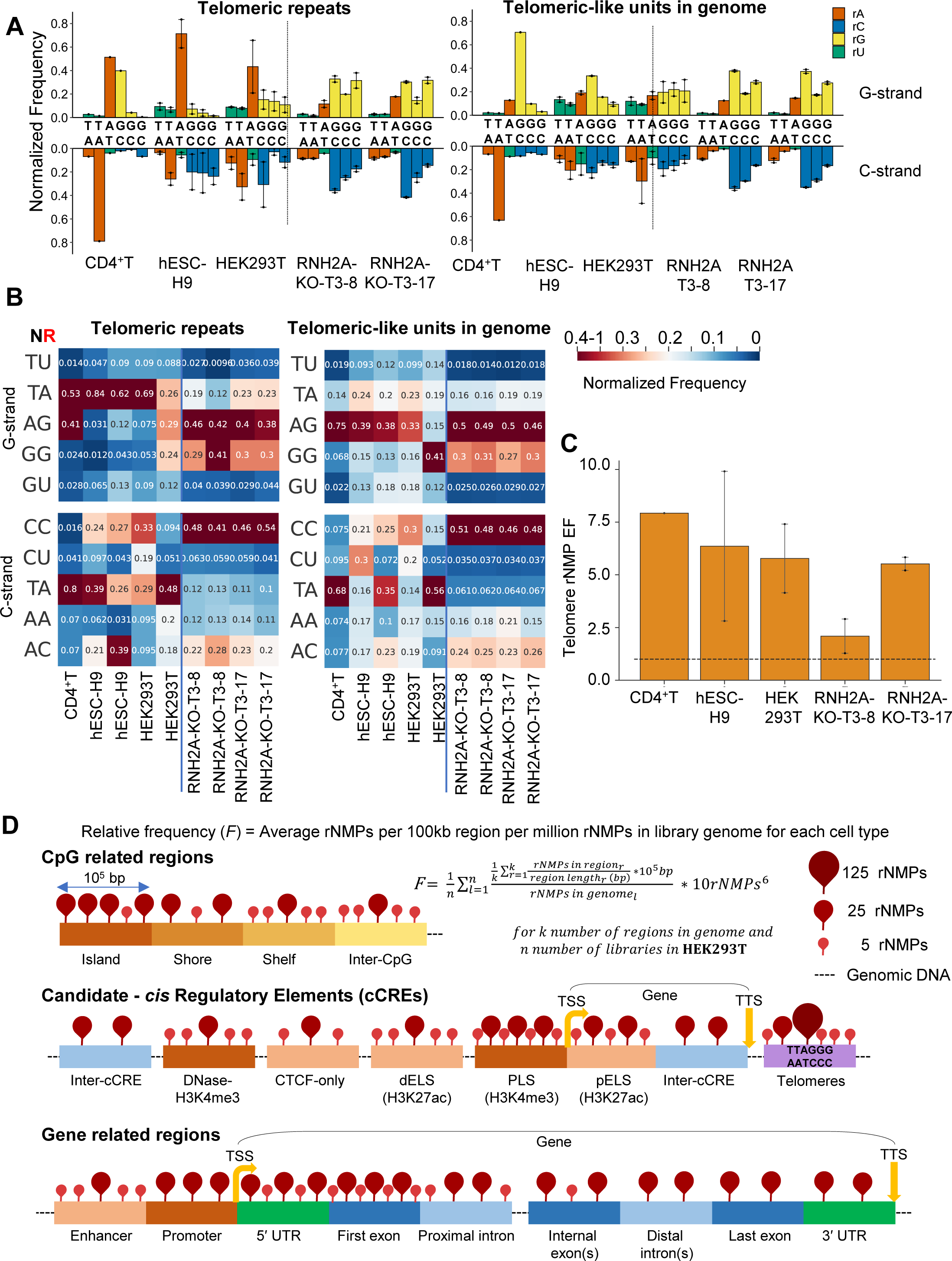
TrA signature characterizes rNMP incorporation in human telomeres. (**A**) Bar graphs showing the mean and standard error of normalized frequencies (*y*-axis) of rNMPs embedded in telomeric repeats (sequencing reads containing >3 telomeric repeat units, TTAGGG/CCCTAA) and isolated telomeric-like units in the genome (≤3 consecutive repeats), for libraries fragmented by the dsDNA Fragmentase. The top graph represents the G strand (TTAGGG) and the bottom graph the C strand (CCCTAA), with nucleotides on the *x*-axis. Bars are grouped by cell type, with RNH2A-KO genotypes separated by a vertical dotted line. (**B**) Heatmaps showing normalized frequencies for NR dinucleotides (rA, rC, rG, and rU paired with upstream dA, dC, dG, or dT) on the G and C strands for rNMPs embedded in telomeric repeats and telomeric-like units in the genome. Frequencies are normalized to the number of dinucleotide occurrences in the telomeric repeat sequence for the G and C strand, respectively. The color scale (legend on the right) ranges from white (0.20) to light red/red (0.20 to 0.4–1) and dark blue/light blue for (0–0.20). (**C**) EF of rNMPs in telomeres calculated based on telomere lengths estimated from whole genome sequencing reads (**table S15** and Methods) for libraries fragmented by the dsDNA Fragmentase. (**D**) Schematic representations of rNMP distribution across annotated genomic features in wild-type HEK293T cells. Top panel: CpG island (dark orange), shore and shelf (lighter shades of orange), and inter-CpG (yellow); each region represents 100-kb genomic segments. Middle panel: cCREs, including PLS (dark orange), dELS and pELS (light orange) with respect to the TSS, regions with CTCF-only signature (light orange), DNase-H3K4me3 signature (orange), inter-cCRE (light blue), and telomeres (purple). Bottom panel: gene-associated regions, including enhancer (1–5 kb upstream of TSS, light orange), promoter (0–1 kb upstream of TSS, orange), 5′ and 3′ UTRs (green), exons (blue) and introns (light blue). Exons are categorized as first, last, and internal exons, and introns as proximal and distal introns to show proximity to TSSs and TTSs. Pin size and color intensity (pink/darkness) indicate rNMP counts per 100 kb per million, as defined by the formula shown in the figure. See data in **table S16**.

## Discussion

We demonstrate that embedded rNMPs are abundant and non-randomly distributed across the human nuclear genome, forming conserved, reproducible patterns across diverse human cell types. Ribose-seq and the Ribose-Map toolkit provided a high-resolution map of the human nuclear *ribome*, revealing that rNMPs preferentially accumulate in C/G-rich regions, especially CpG islands, PLSs (H3K4me3), promoters, 5′ UTRs, and first exons (**Fig. 6D, fig. S13, table S16**). This enrichment was consistent across fragmentation strategies and cell types, indicating biological relevance rather than technical artifacts. These patterns reveal a structural footprint of transcriptional activity, chromatin accessibility, and polymerase usage, suggesting that embedded rNMPs function as a novel form of epigenetic mark, distinct from established DNA or histone modifications, yet capable of both reflecting and influencing gene regulatory states.

We identified rNMP-enriched zones (REZs), broad, strand-symmetric domains with elevated rNMP incorporation that colocalize with CpG islands and R-loop regions. The enrichment of rNMPs within CpG islands and promoters of highly transcribed regions, together with the presence of R-loops reported in these regions (*48–50*), suggests that the human nuclear ribome is shaped by genome organization and transcriptional activity. Strong enrichment near TSSs, particularly within 1 kb downstream, and the absence of enrichment near TTSs point to a specific association with transcription initiation. rNMP levels scale with gene expression and inversely correlates with DNA methylation at TSSs. Hypomethylated CpG islands are a hallmark of transcriptionally active promoters (*51, 52*), consistent with rNMPs marking transcriptionally permissive chromatin. These observations raise the possibility of a feedback loop, in which transcription and R-loop formation promote rNMP incorporation, which in turn may reinforce transcription. Future work is needed to determine whether embedded rNMPs also play a causal role in promoting DNA hypomethylation, a characteristic of RNase H2 mutations in AGS (*53*).

The observed rNMP enrichment near TSSs, particularly for rC and rG, and the associated dinucleotide biases may reflect incorporation by DNA polymerases of rNMPs locally recycled via salvage pathways from degraded RNA in R-loops, which form near TSSs due to the accumulation of hyper-negative supercoiling on the DNA upstream of RNA polymerases in highly transcribed genes (*54, 55*) (**Fig. 4E**). Notably, rNMP enrichment near the TSS is lower in hESC-H9 cells, which are characterized by low CpG methylation (*56, 57*) but also more efficient de novo, rather than salvage, nucleotide synthesis (*58–60*). Interestingly, rNMPs were also enriched in highly methylated TSSs in certain cell lines, potentially reflecting the proximity of hypo- and hypermethylated domains in both linear and 3D genome architecture (*61–63*). In such contexts, R-loop processing in nearby low-methylation regions may promote rNMP incorporation into adjacent highly methylated DNA.

The increased rNMP enrichment in REZs in RNH2A-KO cells suggests that RNase H2 normally cleaves rNMPs in these regions, reflecting an active balance between incorporation and removal. Indeed, RNase H2 may already be positioned near REZs to process the RNA component of R-loops. Consistent with our findings, RNase H2A binding has been shown to depend on active transcription, with enrichment at strong promoters of protein-coding genes and at TSSs of active small nuclear RNAs (*64*). The mechanistic relevance of rNMPs to transcription is further supported by their strong correlation with gene expression in RNH2A-KO cells, more so than in wild-type, and by the greater enrichment on the template strand. Therefore, we propose that RNase H2 by cleaving at rNMPs plays a significant role in relieving torsional stress of replication and then transcription near the TSS of highly transcribed genes particularly along the leading strand to facilitate rapid and efficient transcription after leading strand replication (**Fig. 4F**). In the absence of RNase H2, Top1 may help resolve torsional stress by removing rNMPs. In yeast, Top1 has been proposed to excise rNMPs from the nascent leading strand, alleviating replication-induced supercoiling (*65, 66*), whereas the lagging strand is less affected due to its discontinuous synthesis (*66*). In human cells, origin firing near TSSs promotes co-directional replication with transcription (*67–69*). The template-biased rNMP enrichment and reduced rC embedment on the non-template strand in RNH2A-KO cells, particularly downstream of TSSs, is suggestive of Top1 processing of the non-template leading strand of highly expressed genes. This activity would relieve negative supercoiling behind RNA and DNA polymerases, supporting transcriptional restart and elongation (**fig. S14**). Alternatively, the DEAD-box RNA helicase DDX3X, which has been shown to exhibit RNase H2-like activity on both RNA:DNA hybrids and single rNMPs (*70, 71*), could potentially target rNMPs in the absence of RNase H2 function.

Among individual rNMPs, rG showed the strongest enrichment downstream of TSSs in RNH2A-KO cells, increasing with gene expression, whereas its concentration declined or remained unchanged in wild-type cells, suggesting selective RNase H2 cleavage at rGs, particularly at highly transcribed TSSs. In contrast, rC and rU levels were reduced on the non-template strand in RNH2A-KO but not wild-type cells, supporting a model where Top1 targets these rNMPs specifically on the non-template strand of highly transcribed genes (**Fig. 4F** and **fig. S14**). This interplay may help regulate transcriptional output, as gene expression distribution was somewhat narrower in RNH2A-KO than wild-type cells.

When normalized to dNMPs, rA, rC, and rG were enriched, while rU was least abundant, a pattern inconsistent with rNTP/dNTP pool ratios, suggesting that DNA polymerase activity, chromatin structure, and/or local sequence context contribute to shaping incorporation patterns. The general rNMP composition in nuclear DNA of RNH2A-KO cells (C/G > A > U) mirrors that of human mitochondrial DNA, where RNase H2 is inactive (*37*), implying minimal influence of other nuclear rNMP-processing enzymes like Top1 on overall composition. Sequence preference analysis revealed enrichment of ArA, ArC, and ArG in CpG islands and near TSSs of highly expressed genes, with a relative depletion of CrG. These patterns parallel polymerase-specific rNMP signatures described in yeast, where Pol ε favors ArN and Pol *δ* prefers CrN (*31, 32*), suggesting higher Pol ε activity in CpG islands and near TSSs, and dominant Pol *δ* activity in inter-CpG regions. Strand-specific preferences further support Pol ε activity on the non-template (leading) strand of highly expressed genes.

Finally, we found that telomeres are rNMP incorporation hotspots (**Fig. 6D**), consistent with the known accumulation of RNA:DNA hybrids in these regions, including TERRA-associated R-loops (*72*). In wild-type cells, the telomeric G-strand, synthesized by telomerase, exhibited elevated rA incorporation in TrA motifs, a pattern absent from telomeric-like units elsewhere in the genome, indicating telomerase specificity. This signature was diminished in RNH2A-KO cells, where rNMPs incorporated by replicative polymerases likely obscure telomerase-derived signals. These results suggest that rA incorporated by telomerase is inefficiently removed by RNase H2 and may contribute structurally to telomeric DNA.

Altogether, our study reveals that the human nuclear ribome is not random but highly specific, shaped by DNA sequence, transcription, polymerase activity, and repair. RNase H2 maintains rNMP homeostasis, and its loss reveals a transcription-coupled rNMP footprint. These findings establish DNA-embedded rNMPs as a new type of epigenetic mark, arising not from classical writer/eraser machinery, but from DNA synthesis and selective incision, that links genome sequence, transcriptional activity, and gene regulation. By uncovering a non-canonical mode of DNA marking, our work expands the landscape of known epigenetic features and opens new directions for understanding transcription-coupled genome architecture in human cells and other biological systems.

## Materials and Methods

### Human cell line and genomic DNA preparation

#### Human primary activated CD4^+^T cells

CD4^+^T cells were isolated from human buffy coats of five health donors (NY Blood Center) as previously described (*73*) and the isolated CD4^+^T cells were pooled and activated by phytohemagglutinin (5 ng/ml) and IL-2 (5 ng/ml) for 3 days. The total cellular DNAs of the activated CD4^+^T cells were isolated using a DNA extraction kit (Promega Wizard) for analysis.

#### hESC-H9

Human Embryonic Stem Cells (hESC) line H9, passage 22, were purchased from WiCell Research Institute, University of Wisconsin (line WA09, lot # WB66595). hESC was maintained and expanded in feeder-free culture conditions with mTeSR basal medium (STEMCELL Technologies Inc., Vancouver, Canada, catalog no. 85850) on 6-well plates culture-treated coated with Corning Matrigel hESC-qualified Matrix, LDEV free (Corning Inc., Corning, NY, USA, catalog no. 35427) in a humidified chamber in a 5% CO_2_-air mixture at 37 °C. The culture medium was changed daily, and regions of differentiation were removed by aspiration. Cells were passed as small aggregates every four to five days at ratios of 1:3 to 1:6 using a gentle dissociation reagent (Corning Inc., catalog no. 3010). Cells were harvested at passage 28 as a single-cell suspension for downstream applications.

#### HEK293T

Human embryonic kidney T (HEK293T) RNH2A wild-type and KO (RNH2A-KO) cells were provided by the Pursell group at Tulane University (*37, 74*). Briefly, cells were grown in Dulbecco’s modification of Eagle’s medium (DMEM) containing 4.5 g/l glucose, L-glutamine, and sodium pyruvate (Corning catalog no. 10-013-CV) with 10% fetal bovine serum (Sigma-Aldrich catalog no. F0926-500ML) and 1× penicillin–streptomycin (Gibco catalog no. 15140122). Cells were grown at 37 °C in a 5% CO_2_-humified incubator. For the construction of the RNH2A-KO clones see (*37, 74*). HEK-293T RNH2A-KO T3-8 and T3-17 have three distinct frameshift mutations consistent with all three alleles being modified in hypotriploid 293T cells, respectively. The RNH2A-KO T3-8 has an insertion of G at position −1; deletion of five bases at position +2 to +6; complex alteration with deletion of three bases from position −1 to +2, deletion of two bases from position +5 to +6, and CC > TT at position +8 and +9. The RNH2A-KO T3-17 has the deletion of 14 bases from position +7 to −7; the deletion of six bases from position −5 to 1; and the deletion of 25 bases from position −3 to +22. All positions are indicated with respect to the Cas9 cleavage site on the reference strand.

Genomic DNA from hESC-H9 and HEK293T as well as RNH2A-KO cells was extracted using the Qiagen DNeasy Blood & Tissue Kit (catalog no. 69504).

### Denaturing gel electrophoresis of RNase HII treated genomic DNA with analysis, and simulation of breaks in DNA

5 units of RNase HII (catalog no. M0288L) were added to 250 ng of genomic DNA to induce cleavage at rNMP embedded and incubated for 16 hours at 37 °C. After incubation, 95% Formamide, 1mM EDTA was added to the reaction mixture and 1kb plus DNA ladder (catalog no. N3200L) and was incubated at 95 °C for 10 minutes to induce denaturation of double stranded DNA and cooled down to 4 °C for 5 minutes. 0.8% agarose gel electrophoresis was run at 2V/cm for 10 hours. After running the gel, staining was performed with 1:10,000 SYBR Gold (catalog no. S11494) for 20 minutes. The gel was then imaged using Axygen system using UV302.

We made 4 replicates of gels. Each gel was analyzed from the bottom of the well to the end of the gel to exclude any artifacts from gel edges using gelpy software. The portion of the gel analyzed is indicated with thin black horizontal lines on the right side of the gel image. Each lane in the gel had 100-110 pixels of width (left to right) and 1900-2000 pixels in length (top to bottom).

Each lane is defined by a length *l* and width *w* in pixels. For each position along the length, the intensity is computed as the sum of *w* pixel values across the width.

For normalization of stain intensity in each strip, we converted the intensity in each strip as a percentage of the total intensity in the lane. The percent stain intensity shows the moving average intensity in 50 strips for every strip in the lane. To estimate the fragment length corresponding to each strip, we use an exponential decay curve model:

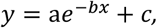

where *y* corresponds to the fragment lengths and *x* corresponds to the location of the strip in the lane. We derived the constants *a, b*, and *c* by fitting the fragment lengths of ladder bands against their respective locations, represented by the highest intensity of each band in the ladder.

To calculate the number of fragments for each corresponding fragment length, *l*_*i*_ at each strip *i*, we first calculated the number of DNA nucleotides N in each lane using the formula

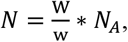

where *W* is the weight of DNA loaded in each lane, for our study its 250 ng, *w* is the average weight of DNA nucleotide, i.e., 330 * 10^9^ ngs/mole and *N*_*A*_ is the number of nucleotides in 1 mole, 6.023 *10^23^.

Let *P*_*i*_ be the percent stain intensity in each strip *i* in a lane, the number of fragments *N*_*i*_ in the strip is calculated using the formula:

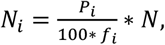

where *f*_*i*_ is the length of fragments at each corresponding strip *i*, predicted using the exponential model, and *N* is the total DNA nucleotides in each lane area as calculated above.

To calculate the median fragment length for treated and untreated sample lanes, we analyzed the top 95% of fragment data (i.e., starting from the bottom of the well). This approach minimizes variation caused by background staining, which is typically more pronounced in regions containing smaller fragments. Within this 95% data subset, we identified the pixel position at which the cumulative fragment count reached 50% of the total, representing the median fragment length. These values are shown in **fig. S1** as blue circles (untreated) and brown triangles (treated).

To estimate the number of rNMPs incorporated in the DNA, we developed a simulation-based method that infers the number of strand breaks by introducing random breaks into the untreated fragment length distribution until the resulting median matches that of the treated sample. For each cell type and gel, we used the distribution of DNA fragment sizes in both untreated (control) and treated samples and extracted the median fragment length from each. These values served as the initial (untreated) median and goal (treated) median fragment lengths. We initiated the simulation with an assumed population of 10 million DNA fragments following the empirical untreated distribution. Starting with an initial guess of N= 2*10^6^, we introduced N random breaks across the fragments. Break points were selected independently and uniformly across the DNA sequence, under the constraint that all resulting fragments must be at least the minimum observed in the untreated gel. After each simulation, we computed the median of the resulting fragment size distribution. If this simulated median fell within 1 nucleotide of the empirically obtained median from the treated sample, we accepted N as the estimated number of breaks and retained the corresponding fragment distribution. If not, we adjusted N using a binary search strategy, iteratively decreasing or increasing the number of breaks based on whether the simulated median was lower or higher than the goal, respectively. The search was terminated when the simulated median was within 1 nucleotide of the target, or the updated value of N differed by less than 1% from the previous estimate. This procedure was repeated independently 10 times for each cell type and gel, producing estimated break counts and corresponding simulated fragment distributions. To estimate the number of DNA bases per ribonucleotide incorporation, we computed the total number of bases in the fragments resulting from the simulated breaks and divided it by the number of breaks introduced. The final value reported is the mean across the 10 simulated replicates. Further details, including pseudocode and implementation, are provided in the Supplementary Materials and associated GitHub repository.

### Ribose-seq library preparation

The libraries in this study were prepared using the ribose-seq method (*31, 37, 75*). To construct libraries with human cell lines, we improved the ribose-seq protocol by (i) introducing NEBNext® dsDNA Fragmentase® (F) (New England Biolabs catalog no. M0348L) to fully fragment the human genomic DNA samples; (ii) optimizing the removal of linear ssDNA process to retain more rNMPs; (iii) applying a new size selection method using HighPrep™ PCR Clean-up System (MagBio Genomics catalog no. AC-60050) to increase the yield of captured rNMPs (*37*). Part of these libraries was used to study rNMPs embedded in human mitochondrial DNA (*37*).

All the commercial enzymes applied in the ribose-seq protocol were used according to the manufacturer’s recommended instructions. Genomic DNA samples were fragmented using dsDNA Fragmentase (F) or a combination of restriction enzymes (RE) to generate small DNA fragments. Various sets of restriction enzymes were applied to fragment human genomic DNA samples, as indicated in table S1. The used combinations were (i) RE1: [HpyCH4V] + [Hpy116II, Eco53KI, RsaI and StuI]; (ii) RE2: [AleI, AluI and PvuII] + [DraI, HaeIII and SspI]; (iii) RE3: [CviKl-1] + [Mly, MscI and MslI].

Following fragmentation with restriction endonucleases, the fragmented DNA was purified by QIAquick PCR Purification Kit (Qiagen catalog no. 28104). The fragments were tailed with dATP (New England Biolabs catalog no. N0440S) by using Klenow Fragment (3′→5′ exo-) (New England Biolabs catalog no. M0212L) for 30 min at 37 °C and purified by using a QIAquick PCR Purification Kit. In case of fragmentation with NEBNext® dsDNA Fragmentase®, NEBNext End Repair Module (New England Biolabs catalog no. E6050L) was performed before dA-tailing to convert fragmented DNA to blunt-ended DNA having 5′ phosphates and 3′ hydroxyls. Following dA-tailing and purification, a partially double-stranded adapter (Adapter.L1 - Adapter.L8 with Adapter.S (*31*), **table. S3**) was annealed with the DNA fragments by T4 DNA ligase (New England Biolabs catalog no. M0202L) incubating overnight at 16 °C. Following overnight ligation, the products were purified using the HighPrep™ PCR Clean-up System. The adapter-annealed fragments were then treated with working concentration of 0.3 M NaOH for 2 h at 55 °C to denature the DNA strands and cleave the 3′ site of embedded rNMP sites resulting in 2′,3′-cyclic phosphate, and 2′-phosphate termini. After the treatment with Alkali, neutralization using 2 M HCl and purification using HighPrep™ RNA Elite Clean-up System (MagBio Genomics catalog no. RC90050) was performed. Further purification steps were performed using HighPrep™ RNA Elite Clean-up System, except for the size selection step.

The single-stranded DNA (ssDNA) fragments were then incubated with 1 μM *Arabidopsis thaliana* tRNA ligase (AtRNL), 50 mM Tris–HCl pH 7.5, 40 mM NaCl, 5 mM MgCl_2_, 1 mM DTT, and 300 μM ATP for 1 h at 30 °C to ligate the 5′-phosphate and 3′-OH of ssDNA fragments containing an rNMP, followed by bead purification. The ssDNA fragments were treated with T5 Exonuclease (New England Biolabs catalog no. M0633L) for 30 min at 37°C to degrade the unligated linear ssDNA fragments. After purification, the circular fragments were incubated with 1 μM 2′-phosphotransferase (Tpt1), 20 mM Tris–HCl pH 7.5, 5 mM MgCl_2_, 0.1 mM DTT, 0.4% Triton X-100 and 10 mM NAD^+^ for 1 h at 30 °C to remove the 2′-phosphate at the ligation junction.

Following purification, the circular ssDNA fragments were amplified with two steps of PCR to create a library. Both PCR began with an initial denaturation at 98 °C for 30 s, then continued with PCR cycles having denaturation at 98 °C for 10 s, primer annealing at 65 °C for 30 s, and DNA extension at 72 °C for 30 s. These PCR steps were performed for 6 cycles in the first PCR round and 11 cycles in the second PCR round. Lastly, there is a final extension reaction at 72 °C for 2 min for both PCR rounds. The first round of PCR was performed to extend the sequences of the Illumina adapter for TruSeq CD Index primers. The primers (PCR.1 and PCR.2) used for the first round were the same for all libraries. The second round of PCR was performed to insert specific indexes i7 and i5 into each library (*31*). Both PCR rounds were performed with Q5-High Fidelity polymerase (New England Biolabs catalog no. M0491L) for 6 and 11 cycles, respectively. Following the PCR cycles, DNA fragments between 250 and 700 bp were purified using the HighPrep™ PCR Clean-up System. The resulting ribose-seq libraries were mixed at equimolar concentrations and normalized to 4 nM. The libraries were sequenced on an Illumina Next 500 or HiSeq X Ten in the Molecular Evolution Core Facility at the Georgia Institute of Technology or Admera Health.

### Preliminary data processing and alignment

For the ribose-seq libraries, the sequencing reads consist of an eight-nucleotide UMI, a three-nucleotide molecular barcode, the tagged nucleotide (the nucleotide tagged during ribose-seq from which the position of the rNMP is determined), and the sequence directly downstream from the tagged nucleotide. The UMI corresponds to sequence position 1–6 and 10–11, the molecular barcode corresponds to position 7–9, the tagged nucleotide corresponds to position 12, and the tagged nucleotides downstream of the sequence corresponds to positions 13+ of the raw FASTQ sequences. The rNMP is the reverse complement of the tagged nucleotide. Before aligning the sequencing reads to the reference genome, the reads were trimmed based on sequencing quality and the custom ribose-seq adaptor sequence using cutadapt 1.16 (−q 15 -m 62 -a “AGTTGCGACACGGATCTATCA”). In addition, to ensure accurate alignment to the reference genome, reads containing fewer than 50 nucleotides of genomic DNA (those nucleotides located downstream from the tagged nucleotide) after trimming were discarded. Following quality control, the Alignment and Coordinate Modules of the Ribose-Map toolkit (*41, 76, 77*) were used to process and analyze the reads. The Alignment Module de-multiplexed the trimmed reads by the appropriate molecular barcode, aligned the reads to the reference genome (GrCh38) (*78*) using Bowtie 2 (*79, 80*), and de-duplicated the aligned reads using UMI-tools (*81*). Based on the alignment results, the Coordinate Module filtered the reads to retain only those with a mapping quality score of at least 30 (probability of misalignment <0.001) and calculated the chromosomal coordinates and per-nucleotide counts of rNMPs.

Due to the efficient removal of rNMPs by RNase H2, nuclear libraries of wild-type cells generally had a much lower number of reads compared to the mitochondrial libraries of the same cells, and thus had a higher number of background reads that originated from the capture of restriction enzyme ends likely by residual activity of T4 DNA ligase. To improve the quality of analysis and filter out any background reads or artifacts, we filtered out those reads that showed restriction enzyme cleavage sites at rNMP embedment sites, which removes any linear single strand DNA-only fragments being captured. We then also filter out rNMP mismatches between the nucleotide in the reference genome versus the sequencing reads. The final file generated is of BED format and all further analysis is performed this bed file after filtration of restriction enzyme site and mismatched rNMPs.

### Random control rNMP set creation and ribonucleotide distribution histogram analysis

We created a random control rNMP set by distributing an equal number of randomly assigned rNMP sites across the autosomal chromosomes for each library, matching the observed frequencies of biological rNMPs at each site. For a given library, let *N*_*c*_ denote the number of ribonucleotides detected on the chromosome *c*, where *c* ∈ {1,2, …, 22}. We draw *N*_1_ + … + *N*_22_ positions from the set of all non-gap positions (A, C, G, T nucleotides) on chromosomes 1 to 22, ensuring uniform rNMP frequencies per site. If multiple ribonucleotides were detected at the same position, the random control set matched both the number of sites and the number of ribonucleotides per site to the biological rNMP data in each library.

We then generated 100 kb discrete, equally spaced bins of the GRCh38 human genome assembly in the following way. Each chromosome was separately partitioned into 100-KB bins starting from the 5′ end of the chromosome to the 3′ end on the positive strand and bins on negative strand are made to align exactly with those on the positive strand, with the last bin on the 3′ end to be of variable length depending on the chromosome. Bins fully within masked/gap regions of GRCh38 (represented by N’s) do not contain any ribonucleotides; hence, these bins were filtered out from further analysis so that they do not contribute to non-biological zero-inflation. The remaining bins are called “valid” bins. As noted above, even valid bins may partially contain some gaps (N’s) and the last bin on each chromosome may be of length less than 100 kb; hence, we perform length normalization of ribonucleotide counts present in each bin as mentioned under mono and dinucleotide sequence context correlations.

Ribonucleotide Distribution Histogram Analysis was conducted as follows. Let {*R*_*i*_} denote the number of ribonucleotides within the *i*^th^ bin of the sample being analyzed and let {*S*_*i*_} denote the number of ribonucleotides within the *i*^th^ bin of the corresponding random sample. Note that the indices *i* range over only indices of bins in chromosomes 1-22. We then created a histogram of the values {*R*_*i*_} (in orange) and {*S*_*i*_} (in green). The *x*-axis limit was set to the 99^th^ percentile of the {*R*_*i*_} histogram to exclude outliers from the visualizations and enable clear distinction between the distributions of both biological rNMPs and random control rNMP sets.

### Mono and dinucleotide sequence context correlations

We determined the Pearson correlations between the rNMP count fraction and mono and di dNMP count fraction in 1 kb, 10 kb, 100 kb and 1 Mb discrete bins across Watson and Crick strand of autosomal chromosomes. Respective width of discrete bins was created and filtered using the method mentioned under ribonucleotide distribution histogram analysis in Methods. Now let {*R*_*i*_} denote the frequency of ribonucleotides within the *i*^th^ bin of the sample being analyzed and let {*A*_*i*_},{*C*_*i*_},{*G*_*i*_},{*T*_*i*_} denote the number of the respective nucleotides within the *i*^th^ bin. We normalized all quantities rNMPs and dNMPs by the dividing by *A*_*i*_, *C*_*i*_, *G*_*i*_, *T*_*i*_ before computing the correlations. That is, define *R*_*n,i*_ *= R*_*i*_ */ (A*_*i*_ + *C*_*i*_ + *G*_*i*_ + *T*_*i*_) and *A*_*n,i*_ *= A*_*i*_ */ (A*_*i*_ + *C*_*i*_ + *G*_*i*_ + *T*_*i*_) and similarly for *C*_*n,i*_, *G*_*n,i*,_ and *T*_*n,i*_ (subscript ‘n’ indicates ‘normalized’). Then to compute each correlation for the sample and nucleotide A, C, G, T we computed the Pearson correlation coefficients: corr({*R*_*n,i*_, *A*_*n,i*_}), corr({*R*_*n,i*_, *C*_*n,i*_}), corr({*R*_*n,i*_, *G*_*n,i*_}), and corr({*R*_*n,i*_, *T*_*n,i*_}). An analogous procedure was conducted to compute the correlations for dinucleotides.

### rNMP enrichment factor calculation in different genomic regions

Whether analyzing annotation set elements, 100 bp bins around Transcription start sites (TSS), or correlation with expression and methylation levels, the rNMP enrichment Factor (rNMP EF) calculated throughout the study was defined as the ratio of rNMP per base in region of interest and rNMP per base in the genome. The ribonucleotide Enrichment Factor was calculated as

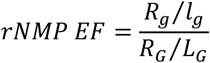

where *R*_*g*_ is number of ribonucleotides in a specific region of the genome with *l*_*g*_ bases, and *R*_*G*_ is total number of ribonucleotides in the genome with *L* bases.

For rNMP heatmaps, correlation with expression and methylation, we analyzed rNMPs on both strands, whereas while studying distribution of rNMPs around TSS, CpG centers and Ribonucleotide Enriched Zones (REZs) we analyzed rNMP EF on separate strands.

### rNMP enrichment distribution analysis around transcription start site (TSS) and CpG centers

The bin size used for studying distribution around TSS and CpG is 100 bases and aligned for both top and bottom strands in non-overlapping (discrete) steps. We calculated rNMP EF for each rNMP in each library with region length as 100 bases followed by filtering those rNMPs 3 kb around the reference point (TSS or CpG) and calculating average of rNMP EF for 100 base discrete bins on each strand separately. Let *i* denote the bin observed from 5′ to 3′ direction across a 3 kb window centered on the reference point, using 100 bp bin widths. This results in 60 bins in total, 30 upstream and 30 downstream of the reference point. The rNMP EF for each library for *i*^th^ bin is calculated as

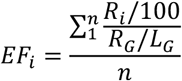

where *R*_*i*_ is count of rNMPs in *i*^th^ bin, *R*_*G*_ is total number of rNMPs in the genome with *L* bases. Here *n* denotes number of regions to be visualized for distribution, e.g. protein coding genes, n = 19,022 genes (*82*) in chromosomes 1 – 22.

When visualizing distributions in discrete bins around the reference point for all cell types in a single plot, the mean *EFi* of the libraries corresponding to each cell type is used to generate the distribution line. When observing each cell type separately, for example, low, moderate, and high expression, the mean and standard error of the *EFi* values from the respective libraries are used to generate the distribution line and error shading in the plot.

A similar method was utilized for studying each rNMP base EF around TSS, defined as

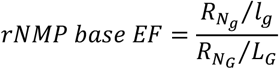

where N ∈ {A, C, G, U} and 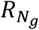 is number of specific rNMP base in an observed region g with length *l*_*g*_ and 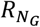 is number of specific rNMP base in the nuclear genome, G (chr 1-22) with length *L*_*G*_.

The percentage of rNMP bases are calculated to specific observed regions, as follows

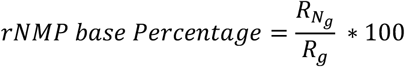

where N ∈ {A, C, G, U} and 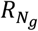 is number of specific rNMP base in an observed region and *R*_*g*_ is the number of all rNMP bases in corresponding region.

### Whole-genome bisulfite sequencing and data processing

The methylome libraries for bisulfite sequencing were made with an in-house Illumina sequencer-compatible protocol (*83*). The extracted DNA was fragmented by S-series Focused-ultrasonicator (Covaris, Woburn, MA) using the “200-bp target peak size protocol.” Fragmented DNA was then size selected (200–600 bp) with an Agencourt AMPure XP bead-based (#A63880, Beckman Coulter, Brea, CA) size selection protocol. The DNA end repair step was performed with End-It DNA End-Repair Kit (ER81050, Epicentre, Madison, WI). After the end-repair step, A-tailing (M0202, New England Biolabs, Ipswich, MA) and ligation steps were performed to ligate the methylated adaptors. Bisulfite treatment of gDNA was performed using the MethylCode Bisulfite Conversion Kit (MECOV50, ThermoFisher). Purified gDNA was treated with CT conversion reagent in a thermocycler for 10 min at 98 °C, followed by 2.5 h at 640 °C. Bisulfite-treated DNA fragments remain single-stranded as they are no longer complementary. Low-cycle (4–8) PCR amplification was performed with Kapa HiFi Uracil Hot start polymerase enzyme (#KK2801, KAPA Biosystems, Wilmington, MA) which can tolerate uracil residues. The final library fragments contain thymines and cytosines in place of the original unmethylated cytosine and methylated cytosines, respectively.

The methylome libraries were diluted and loaded onto an Illumina HiSeq 2500 or HiSeqX system for sequencing using 150 bp paired end reads. We generated over 900 million reads per sample and a standardized WGBS data analysis pipeline was employed for analyzing the resulting data. Overall, we first performed quality and adapter trimming using TrimGalore v0.6.1 (Babraham Institute) with paired-end mode and quality threshold of 20 and trimming 8 bases on both ends for both reads. Subsequently, reads were mapped to the reference genome (GrCh38) using Bismark v0.24.0 (*84*). Following deduplication using Bismark, we generated a total of 3.75 billion reads (equivalent to 561 Giga base pairs of raw sequence data) and obtained per-sample average coverage depths 8-13 x covering > 89% of the CpGs in the autosomal chromosomes (chr 1-chr22). The data, comprising the counts of methylated and unmethylated cytosines in each C-context at individual cytosines, were generated as cytosine report files using Bismark methylation extractor with –bedGraph –cytosine_report –CX_context options. These outputs were used in downstream analyses to calculate the mean methylation level of specific genomic regions.

### RNA sequencing and data processing

RNA samples for HEK293T WT and RNH2A-KO (T3-8 and T3-17) cells were extracted using RNeasy Mini Kit (QIAGEN, Cat: 74104) with on-column DNase treatment. Samples were sent to Admera Health for RNA library prep using PolyA Selection & NEBNext Ultra II Directional kit followed by sequencing with NovaSeq X Plus with 10B flow cell (PE150). The fastq reads were trimmed for any residual Illumina adapters and filtered to keep reads above the quality threshold of 15 and length 50 bp. Alignment of trimmed reads was done on GRCh38 genome using bowtie2 to generate SAM (Sequence Alignment Map) and indexed BAM (Binary Alignment Map) file. Gene exon counts were obtained by featureCounts (*v*.2.0.3) and merged for each gene by custom scripts. Each RNA-seq data set was normalized using FeatureCorr package to log2 Transcript per million (TPM) levels. The clustering of the libraries were checked for outliers, and each genotype library was clustered with same genotypes.

### Correlation trend analysis of rNMP enrichment with expression and methylation

For studying rNMP enrichment correlation with expression levels, we used two regions around the genes, +/− 1 kb around TSS and gene length region from TSS to transcription termination site (TTS). For canonical protein coding genes obtained from rNMP EF was calculated for these regions of protein coding genes and matched with RNA-seq expression levels using Ensembl gene annotations. The RNA-seq datasets used for CD4^+^T and hESC-H9 are GSE139242 (*85*), and GSE175070 (*86*), respectively. obtained from GEO datasets. The expression levels of protein coding genes for each cell type were divided into 10 groups of increasing percentiles: 0-10, 11-20, … 91-100. Let *g* denote gene and *EF*_g_ be the rNMP EF for the gene in each library. For each cell type, the average *EF*_*g*_ was calculated. Using this average EF_g_ we further calculated average *EF*_*G*_, for each group of genes defined by the increasing percentile of 10. The blue and red lines in transcription trend analysis (**Fig. 3D**) connect *EF*_*G*_ from each group to the next with increasing percentile eventually reflecting trend of ribonucleotide embedment with respect to increasing expression levels for genes. The blue lines represent average rNMP EF near TSS (+/− 1 kb) and red indicate average rNMP EF in gene region (TSS to TTS). We used linear fit model, Pearson and Spearman correlation methods implemented in R, to assess linear trends and correlation between rNMP incorporation and gene expression.

A similar analysis was conducted for rNMP Enrichment in +/− 1 kb around protein coding gene TSSs and methylation percentages. CD4^+^T cells whole genome bisulfite sequencing (WGBS) was performed and for hESC-H9 and HEK293T, we utilized GSE16256 (*87, 88*) and GSE97814 (*89*) datasets for correlation and trend analysis. For HEK293T-RNH2A-KOs, we used the same dataset for HEK293T to observe any differences between wildtype RNase H2 and RNH2A-KO genotypes. Furthermore, for analysis of rNMP enrichment with methylation, we separated the CpG islands based on their vicinity to TSS by 3 kb using custom code and bed tools to observe the trends for CpG near TSS and CpG, not near TSS groups.

### Neighboring mono and dinucleotide context preference

In this analysis the goal was to examine the local nucleotide context of the rNMPs, rather than global correlations as in the nucleotide correlation analysis. To this end, we computed the local nucleotide frequencies of each sample as follows. For a single sample being analyzed, let {*X*_*i*_} denote the position of each rNMP in the sample where *i* indexes each rNMP in the sample. Note that we allow *X*_*i*_ to contain duplicate positions for different *i* if multiple rNMPs were detected at the same position. For each *X*_*i*_ we counted the frequency of each nucleotide A, C, G, and T, 50 bases around each rNMP index *i* while excluding the position of the rNMP. We summed up these counts over all rNMP indices *i* to obtain total counts *f*_*A*_, *f*_*C*_, *f*_*G*_, *f*_*T*_. We converted these to frequencies by dividing each by the sum: *F*_*A*_ = *f*_*A*_ */(f*_*A*_ *+ f*_*C*_ *+ f*_*G*_ *+ f*_*T*_) and similarly for *F*_*C*_, *F*_*G*_, *F*_*T*_. We obtained background frequencies by counting the number of each nucleotide in the whole region being considered (either the single chromosome or all chromosomes) to obtain *b*_*A*_, *b*_*C*_, *b*_*G*_, *b*_*T*_. We then converted them to frequencies in an analogous manner to obtain *B*_*A*_, *B*_*C*_, *B*_*G*_, *B*_*T*_. Finally, to obtain the frequency ratio as shown in **Fig. 1A**, we computed the quotients *Q*_*A*_ *= F*_*A*_*/B*_*A*_ and similarly for *Q*_*C*_, *Q*_*G*_, *Q*_*T*_. Note that for dinucleotides (AA, AC, …, TG, TT), the steps are identical except we counted overlapping dinucleotides (e.g., the sequence AGA contributes one count of AG and one count of GA).

### Visualization of rNMP enriched zone (REZs) relative to CpG islands

For visualization and analysis of REZs, we used KaryoploteR (*90*) along with custom scripts (*37*). REZs were identified based on the rNMP Enrichment Factor (EF), calculated for 500 kb discrete bins across both DNA strands genome-wide. A region was defined as a REZ if more than 80% of libraries (i.e., at least 23 out of 26) had an rNMP EF of ≥1.2 for five consecutive bins. Thus, the REZs highlighted in blue in Fig. 3 represent genomic regions with rNMP EF ≥1.2 over a contiguous 2.5 Mb stretch in most libraries. CpG islands were annotated on the karyotype plots using varying transparency: regions with high CpG island density appear as darker red, while regions with lower density appear lighter, allowing for visual comparison with REZ distribution.

### DNA enrichment factor calculation and visualization

DNA extracted from the CD4^+^T cells fragmented using different enzyme sets used in this study for ribose-seq was also sequenced using Illumina HiSeq X Ten Platform. The fastq reads were trimmed for any residual Illumina adapters and filtered to keep reads above the quality threshold of 15 and length 50 bp. Alignment of trimmed reads was done on GRCh38 genome using bowtie2 to generate SAM (Sequence Alignment Map) and indexed BAM (Binary Alignment Map) file, which was further used to obtain the DNA-seq enrichment factor (EF), similar to the rNMP EF using customized scripts. The DNA-seq EF was then visualized with CpG islands in the center of the karyotype plot.

### Relative fraction for each rNMP with base A, C, G, or U

We computed the relative fraction of each rNMP with base A, C, G, or U to its corresponding dNMP with base A, C, G, or T, respectively for each chromosome strand and aggregated them by cell type. Concretely, first we split each chromosome strand into discrete 1Mb windows. Then, within each window, we computed the fraction of each rNMP with base A, C, G, or U with respect to the total number of rNMPs *f*_*r*_, and similar fraction for each dNMP with base A, C, G, or T *f*_*d*_. The fraction of rNMPs with base A, C, G, or U was then divided by the fraction of dNMPs with base A, C, G, or T, respectively, to obtain the relative fraction for each rNMP with base A, C, G, or U as follows

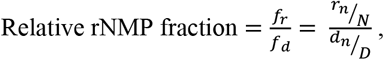

where *r*_*n*_ is number of rNMP with base A, C, G, or U with n ∈ {A,C,G,U} and *N* = *r*_*A*_ + *r*_*C*_ + *r*_*G*_ + *r*_*U*_ and *d*_*n*_ is number of dNMP with base A, C, G, or T with n ∈ {A,C,G,T}and *D* = *r*_*A*_ + *r*_*C*_ + *r*_*G*_ + *r*_*T*_

For each cell type, we displayed mean rNMP fraction for 1Mb window for Watson and Crick strand in Chromosome 19 (**Fig. 5B**).

### rNMP composition and nucleotide preference heatmaps

Because each library can have different coverage, we used percentage of rNMPs in the nuclear genome to determine the rate of embedment in different cell types and genotypes. We then visualized these percentages using horizontal bar graphs generated using customized R scripts. The Composition Module of Ribose-Map was used to obtain raw and normalized percentages of each rNMP base (A, C, G, U/T) in nuclear DNA of all cell types and genotypes. The normalized percentages are the raw rNMP counts normalized on the nucleotide counts in the corresponding (nuclear or mitochondrial) reference genome.

To generate the mononucleotide heatmaps, in every nuclear and mitochondrial ribose-seq library, the frequency of each type of rNMP (*F*_*N*_: *F*_*A*_, *F*_*C*_, *F*_*G*_, *or F*_*U*_) was calculated as a ratio of percentage of each rNMP base divided by the percentage of each dNTP base in the reference genome. Normalized frequency for each rNMP base was calculated using the frequency of the respective rNMP base divided by the sum of frequencies of each rNMP base (*31, 32, 34, 91*). The sum of all normalized frequencies in the mononucleotide heatmaps sums up to 1.

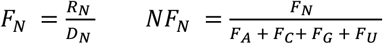

where R_N_ is the percentage of ribo-nucleotides where N ∈ {A, C, G, U}, where D_N_ is the percentage of di-nucleotides in reference genome where N ∈ {A, C, G, T}.

For dinucleotide heatmaps, normalized frequency was calculated for each deoxyribonucleotide immediately upstream or downstream of rNMP embedment site for each base such that the sum of NR (upstream) or RN (downstream) frequencies for each R would sum up to 1 (*31, 91*). For upstream dinucleotide NR, the formula is as follows:

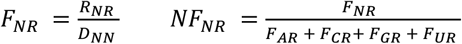

For trinucleotide heatmaps, normalized frequency was calculated for combinations around rNMP (NNR, NRN, RNN) such that the sum of each combination for each R base would sum up to 1. For upstream trinucleotide NNR, the formula is as follows:

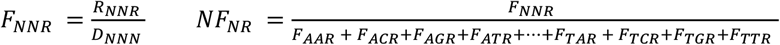

For mono- and di-nucleotide the total number of combinations for each R are 4, where the frequency at random for each combination (R, NR or RN) is 0.25 (1/4). Whereas for trinucleotides the total combinations for each R are 16 making the random base frequency as 0.0625 (1/16).

### rNMP embedment in telomeres

Human telomeres consist of highly conserved tandem repeats of the sequence “TTAGGG” on the G-strand and “CCCTAA” on the C-strand (*92*). We searched for these repeat units in the GRCh38 human genome and found that the longest uninterrupted “TTAGGG” or “CCCTAA” stretches in the reference genome contain only three repeat units. Therefore, any read beginning with four or more consecutive “TTAGGG” or “CCCTAA” repeats can be confidently identified as originating from telomeric DNA. After removing UMIs and PCR duplicates, we located such telomeric reads and identified rNMP incorporation at the first base of each read. As a comparison, we also retrieved rNMPs aligned within telomeric-like units present elsewhere in the genome, defined as rNMPs located within “TTAGGG” or “CCCTAA” motifs of fewer than four repeats in the GRCh38 reference genome. The Python3 scripts used to identify rNMPs in telomeres and telomeric-like units are available on GitHub.

For calculating rNMP EF in telomeres, we first use DNA-seq data for each cell type in this study and get telomeric reads using same method defined above for capturing rNMPs in telomeres. We obtain the ratio of DNA-seq reads aligned using bowtie2 to human reference genome (GrCh38) post trim galore processing with default parameters. Using these parameters and genome length based on known sex of cell lines, we estimated total telomere length in all chromosomes for each cell type. This length of telomeres is then used to calculate the rNMP Enrichment factor.

### Statistical Analysis

We use the two-sided Mann Whitney *U-*test and Kolmogorov-Smirnov test for comparison of random and biological rNMP distribution (**fig. S2**) and two-sided Mann Whitney *U*-test for rNMP EF in genomic annotation sets vs. reference regions. We use two-sided Mann Whitney *U-* test for rNMP EF difference found in template and non-template strand for comparison of low expressed with moderate/high expressed genes (**Fig. 3E**). We use a two-sided Mann Whitney *U-* test for comparison of rNMP EF in the template and non-template strand in each cell type of protein coding genes (**fig. S6**).

For mono-, di- and tri-nucleotide heatmaps, we use one-sided Mann Whitney *U*-test for obtaining preferred nucleotides, i.e., nucleotide combinations that are significantly greater than random frequency of 0.25 for mono- and di-nucleotides and 0.0625 for tri-nucleotides. We highlight those combinations that are significantly different between two groups, e.g., CpG islands and inter-CpG regions using two-sided Mann Whitney *U*-test, while reporting both paired and unpaired test *p*-values.

## Acknowledgments

We thank S. Biliya, N. Djeddar, and A. Bryksin from the Molecular Evolution Core for advice and support with high-throughput sequencing, and the Partnership for an Advanced Computing Environment (PACE) for their research cyberinfrastructure resources and services at the Georgia Institute of Technology, and Thomas Layman for the help with WGBS libraries. We acknowledge G. Gibson, M. Borodovsky, M. Lu, and T. Warner for critical reading of the manuscript, and all members of the Storici laboratory for assistance and feedback on this study. The authors used ChatGPT 4o (OpenAI) to assist with language refinement during manuscript preparation.

## Funding

This work was supported by the National Institutes of Health NIH R01 ES026243 (F.S.), the Howard Hughes Medical Institute Faculty Scholars Award HHMI 55108574 (F.S.), the Mathers Foundation AWD-002589 (F.S.), the W.M. Keck Foundation (F.S.), the Tom and Marie Patton Distinguished Professor at Georgia Tech (F.S.); the National Science Foundation grants NSF CCF-2107267 and NSF DMS-2054321 (N.J.); the NIH AI136581 and AI162633 (B.K.), the MH116695 (R.F.S.); the NIH R01 CA284633 (B.D.F.), the Mark Foundation for Cancer Research (B.D.F.); the NIH R01MH103517 and NSF EF-2021635 (S.V.Y.).

## Author contributions

Conceptualization: D.L.K., T.Y., P.X., T.C., N.J., and F.S.; Data curation: D.L.K., T.C., P.X., F.M.-F., and M.S.; Formal analysis: D.L.K., T.C., P.X., F.M.-F., N.J., and F.S.; Funding acquisition: R.F.S., Z.F.P., B.K., B.D.F., S.V.Y., N.J., and F.S.; Investigation: D.L.K., T.Y., T.C., P.X., F.M.-F., N.J., and F.S.; Methodology: DLK conducted most of the bioinformatic analysis with help from T.C., P.X., F.M.-F., A.M., M.S, and A.G.; T.Y. performed most experiments with help from Y.Lee, S.R., G.N., Y.J., SM, G.N., Y.Lu, S.T., and J.A.L.; Project administration: N.J. and F.S.; Resources: R.F.S., Z.F.P., B.K., N.J., and F.S; Software: D.L.K., T.C., P.X., F.M.-F.; Supervision: A.M., N.J., and F.S.; Validation: D.L.K.; Visualization: D.L.K., T.C., P.X., Y.Lee, F.M.-F., N.J., and F.S.; Writing, original draft: D.L.K. with main input from N.J., and F.S.; Writing, review & editing: all authors.

## Competing interests

We have a patent related to this study: Storici, F., Hesselberth, J.R., and Koh, K. D. Methods to detect Ribonucleotides in deoxyribonucleic acids. GTRC-6522, 2013; U.S. 10,787,703 B1 Sep. 29, 2020. https://uspto.report/patent/grant/10,787,703

## Data and materials availability

All raw and processed data generated for this study, including ribose-seq libraries for all cell lines, DNA-seq and bisulfite-seq for CD4^+^T, RNA-seq for HEK293T wild-type and RNH2A-KO cells has been submitted to Geodataset. All unique/stable materials generated in this study are available from the lead contact.

## Supplementary Materials

Figs. S1 to S14

Tables S1 to S16

