## Supplementary Tables and Figures for "Human *ribomes* reveal DNA-embedded ribonucleotides as a new type of epigenetic mark"

**The PDF file includes:**

Supplementary Tables S1-16
Supplementary Figures S1-14

table S1.
Estimated rNMPs per human nuclear diploid genome across four gel runs, based on newly generated fragments in RNase HII-treated vs. untreated lanes by cell type in gels presented in **fig. S1.**

| **Gel no.** | **$CD4^{+}T$** | **hESC-H9** | **HEK293T** | **HEK293T-RNH2A-KO-T3-8** | **HEK293T-RNH2A-KO-T3-8** | **Mean (KO RNase H2)** |
|---|---|---|---|---|---|---|
| Gel 1 | 767029.197 | 720709.217 | 774673.353 | 993764.4 | 1047303.93 | 1020534.16 |
| Gel 2 | 718090.95 | 709551.601 | 741854.869 | 828589.796 | 821325.128 | 824957.462 |
| Gel 3 | 795701.121 | 717629.246 | 784913.593 | 1098566.98 | 964968.989 | 1031767.98 |
| Gel 4 | 790008.43 | 766000.879 | 815245.145 | 1002235.26 | 997856.773 | 1000046.02 |
| Median | 778518.82 | 719169.231 | 779793.473 | 997999.83 | 981412.881 | 1010290.09 |

**table S2.**
Human nuclear ribose-seq library information including cell type, genotype, library name, fragmentation method/set and respective annotation, UMI barcode and number of cycles for PCR1 and 2 in ribose-seq.

| Cell type | Geno type | Library | Fragmentation | RE set | Barcode | PCR 1 | PCR 2 |
|---|---|---|---|---|---|---|---|
| CD4$^{+}$T | WT | FS185 | NEBNext dsDNA Fragmentase | F | CCG | 10 | 7 |
| | | FS187 | [NEBNext dsDNA Fragmentase] + [HpyCH4V] + [Hpy166II, Eco53KI, RsaI, StuI] | FRE1 | TGT | 6 | 11 |
| | | FS188 | [NEBNext dsDNA Fragmentase] + [HpyCH4V] + [Hpy166II, Eco53KI, RsaI, StuI] | FRE1 | CTG | 6 | 11 |
| | | FS189 | [NEBNext dsDNA Fragmentase] + [HpyCH4V] + [Hpy166II, Eco53KI, RsaI, StuI] | FRE1 | GAC | 6 | 11 |
| | | FS186 | [HpyCH4V] + [Hpy166II, Eco53KI, RsaI, StuI] | RE1 | TGA | 10 | 7 |
| hESC-H9 | WT | FS197 | NEBNext dsDNA Fragmentase | F | GCT | 6 | 11 |
| | | FS198 | NEBNext dsDNA Fragmentase | F | GAC | 6 | 11 |
| | | FS201 | [NEBNext dsDNA Fragmentase] + [HpyCH4V] + [Hpy166II, Eco53KI, RsaI, StuI] | FRE1 | AGC | 6 | 11 |
| | | FS199 | [HpyCH4V] + [Hpy166II, Eco53KI, RsaI, StuI] | RE1 | GAA | 6 | 11 |
| | | FS331 | [AluI, PvuII, AleI] + [SspI, DraI, HaeIII] | RE2 | GAA | 6 | 11 |
| | | FS303 | [Cvikl-1] + [MlyI, MscI, MslI] | RE3 | GCT | 6 | 11 |
| HEK293T | WT | FS326 | NEBNext dsDNA Fragmentase | F | GCT | 6 | 11 |
| | | FS391 | NEBNext dsDNA Fragmentase | F | CTG | 6 | 11 |
| | | FS203 | [NEBNext dsDNA Fragmentase] + [HpyCH4V] + [Hpy166II, Eco53KI, RsaI, StuI] | FRE1 | CCG | 6 | 11 |
| | | FS333 | [AluI, PvuII, AleI] + [SspI, DraI, HaeIII] | RE2 | TGT | 6 | 11 |
| | | FS305 | [Cvikl-1] + [MlyI, MscI, MslI] | RE3 | GAC | 6 | 11 |
| | RNH2A-KO T3-8 | FS327 | NEBNext dsDNA Fragmentase | F | TGT | 6 | 11 |
| | | FS392 | NEBNext dsDNA Fragmentase | F | AGC | 6 | 11 |
| | | FS300 | [HpyCH4V] + [Hpy166II, Eco53KI, RsaI, StuI] | RE1 | TGA | 6 | 11 |
| | | FS306 | [Cvikl-1] + [MlyI, MscI, MslI] | RE3 | TGA | 6 | 11 |
| | | FS309 | [AluI, PvuII, AleI] + [SspI, DraI, HaeIII] | RE2 | GAA | 6 | 11 |
| | RNH2A-KO T3-17 | FS329 | NEBNext dsDNA Fragmentase | F | AGC | 6 | 11 |
| | | FS393 | NEBNext dsDNA Fragmentase | F | GAA | 6 | 11 |
| | | FS301 | [HpyCH4V] + [Hpy166II, Eco53KI, RsaI, StuI] | RE1 | CTG | 6 | 11 |
| | | FS307 | [AluI, PvuII, AleI] + [SspI, DraI, HaeIII] | RE2 | TGT | 6 | 11 |
| | | FS310 | [Cvikl-1] + [MlyI, MscI, MslI] | RE3 | TGT | 6 | 11 |

**table S3.**

Information on oligonucleotides used in this study. Name, length, and sequence of oligonucleotides used in this study are presented. All bold letters in the PCR primers indicate the specific sequence of index used in sequencing. P and Am indicate end modifications of phosphate and amino modifier, respectively. All oligonucleotides were desalted, except those marked with an asterisk (*), which were HPLC purified. All oligonucleotides were synthesized by Integrated DNA Technologies.

| Name | Size | Sequence |
|---|---|---|
| Adapter.L1 * | 65 | 5’ P-NNC CGN NNN NNA GAT CGG AAG AGC GTC GTG TAG GGA AAG AGT GTT GAT AGA TCC GTG TCG CAA C*T |
| Adapter.L2 * | 65 | 5’ P-NNT GAN NNN NNA GAT CGG AAG AGC GTC GTG TAG GGA AAG AGT GTT GAT AGA TCC GTG TCG CAA C*T |
| Adapter.L3 * | 65 | 5’ P-NNG ACN NNN NNA GAT CGG AAG AGC GTC GTG TAG GGA AAG AGT GTT GAT AGA TCC GTG TCG CAA C*T |
| Adapter.L5 * | 65 | 5’ P-NNG CTN NNN NNA GAT CGG AAG AGC GTC GTG TAG GGA AAG AGT GTT GAT AGA TCC GTG TCG CAA C*T |
| Adapter.L6 * | 65 | 5’ P-NNA GCN NNN NNA GAT CGG AAG AGC GTC GTG TAG GGA AAG AGT GTT GAT AGA TCC GTG TCG CAA C*T |
| Adapter.L8 * | 65 | 5’ P-NNT GTN NNN NNA GAT CGG AAG AGC GTC GTG TAG GGA AAG AGT GTT GAT AGA TCC GTG TCG CAA C*T |
| Adapter.S * | 25 | 5’ P-GTT GCG ACA CGG ATC TAT CAA CAC T -Am 3’ |
| PCR.1 | 54 | 5’ GTG ACT GGA GTT CAG ACG TGT GCT CTT CCG ATC TTG ATA GAT CCG TGT CGC AAC |
| PCR.2 | 20 | 5’ ACA CTC TTT CCC TAC ACG AC |
| PCR.701 | 53 | 5’ CAA GCA GAA GAC GGC ATA CGA GAT **CGA GTA AT**G TGA CTG GAG TTC AGA CGT GT |
| PCR.702 | 53 | 5’ CAA GCA GAA GAC GGC ATA CGA GAT **TCT CCG GA**G TGA CTG GAG TTC AGA CGT GT |
| PCR.705 | 53 | 5' CAA GCA GAA GAC GGC ATA CGA GAT **TTC TGA AT**G TGA CTG GAG TTC AGA CGT GT |
| PCR.707 | 53 | 5' CAA GCA GAA GAC GGC ATA CGA GAT **AGC TTC AG**G TGA CTG GAG TTC AGA CGT GT |
| PCR.712 | 53 | 5' CAA GCA GAA GAC GGC ATA CGA GAT **CTA TCG CT**G TGA CTG GAG TTC AGA CGT GT |
| PCR.501 | 57 | 5’ AAT GAT ACG GCG ACC GAG ATC TAC AC**T ATA GCC T**AC ACT CTT TCC CTA CAC GAC |
| PCR.502 | 57 | 5’ AAT GAT ACG GCG ACC GAG ATC TAC AC**A TAG AGG C**AC ACT CTT TCC CTA CAC GAC |
| PCR.503 | 57 | 5’ AAT GAT ACG GCG ACC GAG ATC TAC AC**C CTA TCC T**AC ACT CTT TCC CTA CAC GAC |
| PCR.504 | 57 | 5’ AAT GAT ACG GCG ACC GAG ATC TAC AC**G GCT CTG A**AC ACT CTT TCC CTA CAC GAC |
| PCR.505 | 57 | 5’ AAT GAT ACG GCG ACC GAG ATC TAC AC**A GGC GAA G**AC ACT CTT TCC CTA CAC GAC |
| PCR.506 | 57 | 5’ AAT GAT ACG GCG ACC GAG ATC TAC AC**T AAT CTT A**AC ACT CTT TCC CTA CAC GAC |
| PCR.507 | 57 | 5’ AAT GAT ACG GCG ACC GAG ATC TAC AC**C AGG ACG T**AC ACT CTT TCC CTA CAC GAC |
| PCR.508 | 57 | 5’ AAT GAT ACG GCG ACC GAG ATC TAC AC**G TAC TGA C**AC ACT CTT TCC CTA CAC GAC |

**table S4.**

Ribose-seq distribution comparison *p*-values of rNMP (biological) data and random control set using Mann-Whitney *U* and Kolmogorov-Smirnov test presented in **fig. S2.** Listed are also rNMP counts and fold change in standard deviation of each ribose-seq library in this study. Percentages of individual rNMPs in the nucleus are also shown.

| Library | rNMPs in nucleus | rNMPs 1kb downstream of TSS | fold change in SD of rNMPs vs random set | Mann Whitney *p*-value | Kolmogorov–Smirnov test *p*-value | rAMP (%) | Mean of rAMP (%) / St dev (%) | rCMP (%) | Mean of rCMP (%) / St dev (%) | rGMP (%) | Mean of rGMP (%) / St dev (%) | rUMP (%) | Mean of rUMP (%) / St dev (%) |
|---|---|---|---|---|---|---|---|---|---|---|---|---|---|
| FS185-CD4$^{+}$T | 1250079 | 21096 | 1.42 | 5.17E-229 | 0 | 42.23 | 37.02 / 13.43 | 20.73 | 22.8 / 9.2 | 26.29 | 23.14 / 5.11 | 10.76 | 17.04 / 4.79 |
| FS187-CD4$^{+}$T | 963479 | 6184 | 0.54 | 7.78E-19 | 0 | 57.4 | | 12.35 | | 17.25 | | 13 | |
| FS188-CD4$^{+}$T | 357159 | 4828 | 0.37 | 6.25E-15 | 0 | 34.67 | | 16.6 | | 28.78 | | 19.95 | |
| FS189-CD4$^{+}$T | 424992 | 4955 | 0.29 | 0.052158518 | 0 | 26.78 | | 34.58 | | 18.25 | | 20.4 | |
| FS186-CD4$^{+}$T | 1569753 | 13095 | 1.02 | 1.28E-14 | 0 | 24.04 | | 29.74 | | 25.13 | | 21.09 | |
| FS197-hESC-H9 | 969423 | 12216 | 1.07 | 3.87E-121 | 0 | 25.6 | 29.98 / 9.98 | 24.33 | 24.91 / 11 | 29.41 | 25.55 / 13.99 | 20.67 | 19.56 / 6.27 |
| FS198-hESC-H9 | 1486071 | 15374 | 1.13 | 7.58E-35 | 0 | 26.13 | | 40.02 | | 15.47 | | 18.38 | |
| FS201-hESC-H9 | 1368990 | 11704 | 0.89 | 2.93E-07 | 0 | 22.94 | | 26.5 | | 27.42 | | 23.13 | |
| FS199-hESC-H9 | 875200 | 5377 | 0.68 | 0.001171938 | 0 | 25.02 | | 6.91 | | 50.09 | | 17.98 | |
| FS331-hESC-H9 | 4184439 | 47094 | 2.03 | 1.88E-29 | 0 | 30.48 | | 20.97 | | 20.55 | | 28 | |
| FS303-hESC-H9 | 762387 | 9687 | 1.04 | 2.52E-76 | 0 | 49.73 | | 30.72 | | 10.35 | | 9.2 | |
| FS326-HEK293T | 271552 | 2592 | 0.06 | 6.18E-19 | 0 | 26.6 | 29.62 / 13.16 | 24.61 | 34.95 / 17.56 | 28.21 | 21.51 / 12.2 | 20.58 | 19.99 / 6.87 |
| FS391-HEK293T | 6842523 | 63754 | 2.87 | 9.06E-127 | 0 | 33.36 | | 23.4 | | 21.27 | | 21.97 | |
| FS203-HEK293T | 1028550 | 12251 | 0.91 | 1.20E-36 | 0 | 24.15 | | 22.9 | | 38.25 | | 14.7 | |
| FS333-HEK293T | 447012 | 13987 | 0.99 | 2.39E-281 | 0 | 49.73 | | 40.29 | | 10.26 | | 30.09 | |
| FS305-HEK293T | 1037489 | 17154 | 1.74 | 0 | 0 | 14.27 | | 63.57 | | 9.54 | | 12.62 | |
| FS327-HEK293T-RNH2A-KO-T3-8 | 5990007 | 85757 | 3.47 | 0 | 0 | 25.88 | 27.39 / 1.13 | 44.04 | 36.7 / 4.68 | 25.8 | 31.50 / 3.84 | 4.27 | 4.4 / 0.18 |
| FS392-HEK293T-RNH2A-KO-T3-8 | 6772739 | 139154 | 6 | 0 | 0 | 26.91 | | 38.5 | | 30.34 | | 4.24 | |
| FS300-HEK293T-RNH2A-KO-T3-8 | 41819863 | 807556 | 10.25 | 0 | 0 | 27.13 | | 32.28 | | 36.27 | | 4.32 | |
| FS306-HEK293T-RNH2A-KO-T3-8 | 20724824 | 388710 | 8.34 | 0 | 0 | 28.38 | | 34.41 | | 32.69 | | 4.52 | |
| FS309-HEK293T-RNH2A-KO-T3-8 | 5494700 | 98348 | 4.49 | 0 | 0 | 28.64 | | 34.28 | | 32.41 | | 4.67 | |
| FS329-HEK293T-RNH2A-KO T3-17 | 16502973 | 308983 | 7.63 | 0 | 0 | 30.36 | 31.04 / | 40.54 | 35.4 / | 25.02 | 29.51 / | 4.08 | 4.05 / |
| FS393-HEK293T-RNH2A-KO T3-17 | 10167570 | 184329 | 9.16 | 0 | 0 | 30.71 | | 36.13 | | 29.21 | | 3.95 | |
| FS301-HEK293T-RNH2A-KO T3-17 | 53079330 | 942003 | 10.87 | 0 | 0 | 27.04 | | 38.96 | | 29.81 | | 4.19 | |
| FS307-HEK293T-RNH2A-KO T3-17 | 24690567 | 343770 | 6.43 | 0 | 0 | 34.55 | | 29.22 | | 32.2 | | 4.03 | |

| | | | | | | | | | | | | | |
|---|---|---|---|---|---|---|---|---|---|---|---|---|---|
| FS310-HEK293T-RNH2A-KO T3-17 | 949701 | 11069 | 1.05 | 2.35E-91 | 0 | 32.56 | 2.79 | 32.14 | 4.7 | 31.31 | 2.78 | 3.99 | 0.09 |

**table S5.**

Mann-Whitney *U*-test *p*-values for rNMP EF in genomic annotation sets vs. reference region specific to data in heatmaps shown in **Fig. 1B**. For CpG related annotations, inter-CpG is used as a reference, for candidate *cis*-regulatory elements, inter-cCRE, and for gene related regions, distal introns are used as a reference. *P*-values < 0.05 are highlighted in red.

| Annotation Set | Comparisons | $CD4^+T$ | hESC-H9 | HEK293T | HEK293T-RNH2A-KO-T3-8 | HEK293T-RNH2A-KO-T3-17 |
|---|---|---|---|---|---|---|
| CpG related regions | Island vs Inter-CpG | 0.30952 | 0.13203 | 0.00794 | 0.00794 | 0.00794 |
| | Shore vs Inter-CpG | 0.30952 | 0.09307 | 0.03175 | 0.00794 | 0.00794 |
| | Shelf vs Inter-CpG | 0.69913 | 0.39394 | 0.30952 | 0.54762 | 0.30952 |
| Candidate - *cis* Regulatory Elements (cCREs) | dELS (H3K27ac) vs Inter-cCRE | 0.69913 | 1 | 0.84127 | 0.09524 | 0.22222 |
| | PLS (H3K4me3) vs Inter-cCRE | 0.30952 | 0.06494 | 0.03175 | 0.00794 | 0.00794 |
| | pELS (H3K27ac) vs Inter-cCRE | 0.30952 | 0.01515 | 0.03175 | 0.00794 | 0.00794 |
| | DNase-H3K4me3 vs Inter-cCRE | 0.58874 | 0.13203 | 0.69048 | 0.22222 | 0.30952 |
| | CTCF-only vs Inter-cCRE | 0.06494 | 0.00433 | 0.00794 | 0.00794 | 0.00794 |
| Candidate - *cis* Regulatory Elements (cCREs) | Enhancer vs distal intron(s) | 0.93723 | 0.48485 | 0.69048 | 0.09524 | 0.09524 |
| | Promoter vs distal intron(s) | 0.24026 | 0.17965 | 0.05556 | 0.00794 | 0.00794 |
| | 5' UTR vs distal intron(s) | 0.30952 | 0.30952 | 0.15079 | 0.00794 | 0.00794 |
| | First exon vs distal intron(s) | 0.30952 | 0.30952 | 0.09524 | 0.00794 | 0.00794 |
| | Proximal intron vs distal intron(s) | 0.30952 | 0.30952 | 0.22222 | 0.01587 | 0.05556 |
| | Internal exon(s) vs distal intron(s) | 0.93723 | 0.93723 | 0.69048 | 0.22222 | 0.42063 |
| | Last exon vs distal intron(s) | 0.69913 | 0.24026 | 0.84127 | 0.54762 | 0.69048 |
| | 3' UTR vs distal intron(s) | 0.81818 | 0.13203 | 0.84127 | 0.69048 | 0.69048 |

**table S6.**

Association statistics of rNMPs 1 kb around TSS and respective methylation for 10 percentile expression groups of respective cell types specific to data presented in **Fig. 3B**. Methylation percentages used for RNH2A-KOs is that of HEK293T WT for comparison.

| Region | Statistics | $CD4^+T$ | hESC-H9 | HEK293T | HEK293T-RNH2A-KO-T3-8 | HEK293T-RNH2A-KO-T3-8 |
|---|---|---|---|---|---|---|
| 1kb around all protein coding genes TSSs | linear model ($R^2$) | 0.8762 | 0.9040 | 0.7715 | 0.6340 | 0.5314 |
| | Pearson's *R* | -0.9434 | -0.9564 | -0.8927 | -0.8214 | -0.7639 |
| | Pearson *p*-value | 4.20E-05 | 1.50E-05 | 5.09E-04 | 3.57E-03 | 1.01E-02 |
| | Spearman's' *rho* | -0.9879 | -0.5515 | -0.9515 | -0.9273 | -0.9152 |
| | Spearman's *p*-value | 0.0000 | 1.04E-01 | 0.0000 | 1.30E-04 | 4.67E-04 |

**table S7.**

Association statistics of rNMPs 1 kb around TSS and in gene region (TSS – TTS) with expression ($log_2$TPM) for 10 percentile expression groups of respective cell types specific to data presented in **Fig. 3D**.

| Region | Statistics | $CD4^+T$ | hESC-H9 | HEK293T | HEK293T-RNH2A-KO-T3-8 | HEK293T-RNH2A-KO-T3-8 |
|---|---|---|---|---|---|---|
| +/- 1kb TSS | linear model ($R^2$) | 0.8758 | 0.9124 | 0.9452 | 0.9400 | 0.9227 |
| | Pearson's *R* | 0.9432 | 0.9603 | 0.9753 | 0.9729 | 0.9650 |
| | Pearson *p*-value | 4.26E-05 | 1.04E-05 | 1.57E-06 | 2.27E-06 | 6.27E-06 |
| | Spearman's' *rho* | 0.8909 | 0.9515 | 0.9394 | 1.0000 | 0.9758 |
| | Spearman's *p*-value | 0.0014 | 0.0000 | 0.0000 | 0.0000 | 0.0000 |
| gene (TSS to TTS) | linear model ($R^2$) | -0.0558 | 0.6793 | 0.4117 | 0.1332 | 0.2985 |
| | Pearson's *R* | -0.2479 | 0.8455 | 0.6907 | 0.4791 | 0.6135 |
| | Pearson *p*-value | 0.4898 | 0.0021 | 0.0270 | 0.1612 | 0.0592 |
| | Spearman's' *rho* | -0.4182 | 0.7818 | 0.6242 | 0.4061 | 0.6000 |
| | Spearman's *p*-value | 0.2324 | 0.0117 | 0.0602 | 0.2474 | 0.0731 |

**table S8.**

Two-sided Mann Whitney *U*-test *p*-values for rNMP EF difference between template and non-template strand for 1 kb downstream of TSS and between TSS to TTS among three predefined Low, Moderate (mod) and High expression groups for each cell type with Low Expression group as reference for comparison, visually represented in **Fig. 3F**. Significance levels: * ($0.05 \geq p > 0.01$), ** ($0.01 \geq p > 0.001$), *** ($p < 0.001$).

| Cell type | Expression comparison groups | | Two-sided Mann Whitney *U* test *p*-values | |
|---|---|---|---|---|
| | group1 | group2 | 1kb downstream of TSS | TSS-TTS |
| CD4T | low | mod | 0.421 | 1 |
| CD4T | low | high | 0.31 | 0.841 |
| hESC-H9 | low | mod | 0.937 | 0.394 |
| hESC-H9 | low | high | 1 | 0.589 |
| HEK293T-WT | low | mod | 0.548 | 0.69 |
| HEK293T-WT | low | high | 1 | 0.548 |
| RNH2A-KO-T3-8 | low | mod | 0.016 | 0.016 |
| RNH2A-KO-T3-8 | low | high | 0.032 | 0.008 |
| RNH2A-KO-T3-17 | low | mod | 0.114 | 0.008 |
| RNH2A-KO-T3-17 | low | high | 0.029 | 0.008 |

#### table S9.

Pearson's *R*, *p*-value, and linear fit slope for each rNMP base (A, C, G and U) counts, EF and percentage in relation to 10 expression percentile groups in each cell type specific to data presented in **Fig. 4A-C, figs. S8,9**. Expression levels used for RNH2A-KOs are those of HEK293T WT.

| Both Strands | | | | | | | | | | | | | | |
|---|---|---|---|---|---|---|---|---|---|---|---|---|---|---|
| | Stats | Ribo | 0-1 kb down stream of TSS | | | | 4-5 kb down stream of TSS | | | | 3-2 kb up stream of TSS | | | |
| | | | CD4$^+$T | hESC-H9 | HEK WT | KO avg | CD4$^+$T | hESC-H9 | HEK WT | KO avg | CD4$^+$T | hESC-H9 | HEK WT | KO avg |
| rNMP counts | *R* | rA | 0.866 | 0.601 | 0.898 | 0.892 | -0.224 | 0.405 | 0.866 | 0.806 | -0.291 | 0.298 | 0.886 | 0.827 |
| | | rC | 0.645 | 0.800 | 0.973 | 0.967 | -0.805 | 0.707 | 0.680 | 0.696 | -0.231 | 0.921 | 0.682 | 0.760 |
| | | rG | 0.621 | 0.803 | 0.909 | 0.938 | -0.855 | 0.765 | 0.780 | 0.749 | -0.512 | 0.686 | 0.856 | 0.793 |
| | | rU | 0.672 | 0.538 | 0.857 | 0.883 | -0.049 | 0.799 | 0.740 | 0.745 | 0.511 | 0.626 | 0.835 | 0.767 |
| | *p*-value | rA | 1.19E-03 | 6.61E-02 | 4.20E-04 | 5.28E-04 | 5.35E-01 | 2.45E-01 | 1.19E-03 | 4.84E-03 | 4.14E-01 | 4.02E-01 | 6.35E-04 | 3.18E-03 |
| | | rC | 4.41E-02 | 5.49E-03 | 2.29E-06 | 4.80E-06 | 4.92E-03 | 2.21E-02 | 3.05E-02 | 2.53E-02 | 5.20E-01 | 1.57E-04 | 2.99E-02 | 1.08E-02 |
| | | rG | 5.53E-02 | 5.20E-03 | 2.71E-04 | 6.10E-05 | 1.63E-03 | 9.89E-03 | 7.75E-03 | 1.26E-02 | 1.31E-01 | 2.85E-02 | 1.59E-03 | 6.24E-03 |
| | | rU | 3.32E-02 | 1.09E-01 | 1.54E-03 | 7.04E-04 | 8.92E-01 | 5.53E-03 | 1.44E-02 | 1.34E-02 | 1.32E-01 | 5.26E-02 | 2.61E-03 | 9.59E-03 |
| | slope | rA | 0.023 | 0.017 | 0.040 | 0.541 | -0.003 | 0.012 | 0.040 | 0.408 | -0.005 | 0.007 | 0.035 | 0.359 |
| | | rC | 0.024 | 0.056 | 0.122 | 1.357 | -0.024 | 0.027 | 0.048 | 0.648 | -0.005 | 0.038 | 0.034 | 0.662 |
| | | rG | 0.016 | 0.036 | 0.060 | 1.455 | -0.019 | 0.030 | 0.038 | 0.811 | -0.014 | 0.021 | 0.032 | 0.782 |
| | | rU | 0.016 | 0.010 | 0.031 | 0.080 | 0.000 | 0.024 | 0.019 | 0.049 | 0.004 | 0.016 | 0.027 | 0.052 |
| rNMP EF | *R* | rA | 0.7901 | 0.7008 | 0.8815 | 0.9045 | -0.2393 | 0.6569 | 0.7005 | 0.8105 | -0.372 | 0.372 | 0.740 | 0.861 |
| | | rC | 0.6436 | 0.8686 | 0.9381 | 0.9771 | -0.8037 | 0.6427 | 0.6993 | 0.6480 | 0.053 | 0.922 | 0.684 | 0.756 |
| | | rG | 0.7404 | 0.6938 | 0.9088 | 0.9582 | -0.8850 | 0.8165 | 0.4820 | 0.7155 | -0.340 | 0.550 | 0.550 | 0.757 |
| | | rU | 0.6223 | 0.7125 | 0.8600 | 0.8903 | -0.2093 | 0.8204 | 0.7750 | 0.6794 | -0.363 | 0.499 | 0.510 | 0.863 |
| | *p*-value | rA | 6.54E-03 | 2.40E-02 | 7.45E-04 | 3.24E-04 | 5.05E-01 | 3.91E-02 | 2.41E-02 | 4.45E-03 | 2.90E-01 | 2.90E-01 | 1.44E-02 | 1.38E-03 |
| | | rC | 4.46E-02 | 1.11E-03 | 5.94E-05 | 1.17E-06 | 5.09E-03 | 4.50E-02 | 2.44E-02 | 4.28E-02 | 8.85E-01 | 1.49E-04 | 2.90E-02 | 1.15E-02 |
| | | rG | 1.43E-02 | 2.61E-02 | 2.71E-04 | 1.26E-05 | 6.64E-04 | 3.95E-03 | 1.58E-01 | 2.00E-02 | 3.36E-01 | 9.97E-02 | 9.95E-02 | 1.12E-02 |
| | | rU | 5.47E-02 | 2.08E-02 | 1.41E-03 | 5.53E-04 | 5.62E-01 | 3.64E-03 | 8.47E-03 | 3.07E-02 | 3.02E-01 | 1.42E-01 | 1.32E-01 | 1.29E-03 |
| | slope | rA | 0.047 | 0.021 | 0.048 | 0.059 | -0.009 | 0.023 | 0.043 | 0.039 | -0.015 | 0.009 | 0.031 | 0.038 |
| | | rC | 0.082 | 0.081 | 0.178 | 0.131 | -0.069 | 0.032 | 0.052 | 0.049 | 0.003 | 0.051 | 0.044 | 0.053 |
| | | rG | 0.075 | 0.048 | 0.141 | 0.153 | -0.063 | 0.050 | 0.031 | 0.071 | -0.020 | 0.033 | 0.034 | 0.070 |
| | | rU | 0.063 | 0.033 | 0.114 | 0.059 | -0.007 | 0.053 | 0.053 | 0.032 | -0.013 | 0.022 | 0.030 | 0.040 |
| rNMP Percentage | *R* | rA | 0.235 | -0.295 | -0.658 | -0.690 | 0.623 | -0.527 | -0.465 | -0.469 | 0.129 | -0.413 | -0.644 | -0.686 |
| | | rC | -0.020 | 0.314 | 0.561 | -0.111 | -0.660 | 0.030 | 0.582 | -0.009 | 0.238 | 0.550 | 0.726 | -0.368 |
| | | rG | -0.298 | 0.056 | -0.202 | 0.862 | -0.037 | -0.163 | 0.127 | 0.659 | -0.679 | 0.273 | 0.548 | 0.805 |
| | | rU | 0.156 | -0.342 | -0.456 | -0.695 | 0.315 | 0.496 | -0.319 | -0.508 | 0.436 | -0.313 | -0.672 | -0.255 |
| | *p*-value | rA | 5.14E-01 | 4.08E-01 | 3.88E-02 | 2.74E-02 | 5.43E-02 | 1.17E-01 | 1.76E-01 | 1.72E-01 | 7.23E-01 | 2.35E-01 | 4.44E-02 | 2.84E-02 |
| | | rC | 9.57E-01 | 3.77E-01 | 9.16E-02 | 7.61E-01 | 3.79E-02 | 9.35E-01 | 7.78E-02 | 9.81E-01 | 5.08E-01 | 9.97E-02 | 1.73E-02 | 2.96E-01 |
| | | rG | 4.03E-01 | 8.78E-01 | 5.75E-01 | 1.35E-03 | 9.19E-01 | 6.52E-01 | 7.26E-01 | 3.82E-02 | 3.08E-02 | 4.46E-01 | 1.01E-01 | 4.93E-03 |
| | | rU | 6.67E-01 | 3.33E-01 | 1.85E-01 | 2.58E-02 | 3.75E-01 | 1.45E-01 | 3.69E-01 | 1.34E-01 | 2.08E-01 | 3.79E-01 | 3.34E-02 | 4.76E-01 |
| | slope | rA | 0.001 | -0.001 | -0.001 | -0.002 | 0.006 | -0.002 | -0.003 | -0.002 | 0.075 | -0.278 | -0.348 | -0.263 |
| | | rC | 0.000 | 0.002 | 0.003 | 0.000 | -0.007 | 0.000 | 0.003 | 0.000 | 0.113 | 0.332 | 0.328 | -0.050 |
| | | rG | -0.002 | 0.000 | -0.001 | 0.002 | 0.000 | -0.001 | 0.001 | 0.002 | -0.495 | 0.098 | 0.224 | 0.331 |
| | | rU | 0.001 | -0.001 | -0.001 | 0.000 | 0.002 | 0.003 | -0.001 | 0.000 | 0.307 | -0.152 | -0.204 | -0.017 |
| Non-template Strand | | | | | | | | | | | | | | |
| | Stats | Ribo | 0-1 kb down stream of TSS | | | | 4-5 kb down stream of TSS | | | | 3-2 kb up stream of TSS | | | |
| | | | CD4$^+$T | hESC-H9 | HEK WT | KO avg | CD4$^+$T | hESC-H9 | HEK WT | KO avg | CD4$^+$T | hESC-H9 | HEK WT | KO avg |
| rNMP counts | *R* | rA | 0.823 | 0.426 | 0.699 | 0.857 | -0.193 | -0.412 | 0.672 | 0.696 | -0.099 | -0.529 | 0.872 | 0.868 |
| | | rC | 0.760 | 0.611 | 0.942 | 0.912 | -0.680 | 0.401 | 0.654 | 0.552 | -0.152 | 0.438 | 0.793 | 0.775 |
| | | rG | 0.628 | 0.704 | 0.891 | 0.933 | -0.905 | 0.366 | 0.744 | 0.704 | -0.771 | 0.278 | 0.830 | 0.781 |
| | | rU | 0.573 | 0.639 | 0.859 | 0.771 | -0.349 | 0.800 | 0.732 | 0.552 | -0.186 | 0.345 | 0.828 | 0.699 |
| | *p*-value | rA | 3.44E-03 | 2.20E-01 | 2.44E-02 | 1.51E-03 | 5.93E-01 | 2.36E-01 | 3.34E-02 | 2.54E-02 | 7.85E-01 | 1.16E-01 | 1.02E-03 | 1.12E-03 |
| | | rC | 1.07E-02 | 6.03E-02 | 4.50E-05 | 2.36E-04 | 3.06E-02 | 2.51E-01 | 4.04E-02 | 9.82E-02 | 6.75E-01 | 2.06E-01 | 6.21E-03 | 8.49E-03 |
| | | rG | 5.21E-02 | 2.32E-02 | 5.48E-04 | 8.21E-05 | 3.21E-04 | 2.98E-01 | 1.36E-02 | 2.29E-02 | 9.08E-03 | 4.37E-01 | 2.99E-03 | 7.71E-03 |
| | | rU | 8.31E-02 | 4.69E-02 | 1.45E-03 | 9.03E-03 | 3.23E-01 | 5.42E-03 | 1.60E-02 | 9.81E-02 | 6.06E-01 | 3.28E-01 | 3.12E-03 | 2.45E-02 |
| | slope | rA | 0.016 | 0.008 | 0.010 | 0.224 | -0.002 | -0.006 | 0.017 | 0.134 | -0.001 | -0.008 | 0.022 | 0.188 |
| | | rC | 0.018 | 0.018 | 0.049 | 0.490 | -0.010 | 0.007 | 0.024 | 0.225 | -0.002 | 0.007 | 0.026 | 0.388 |
| | | rG | 0.015 | 0.029 | 0.031 | 0.849 | -0.010 | 0.006 | 0.019 | 0.352 | -0.010 | 0.004 | 0.014 | 0.384 |
| | | rU | 0.008 | 0.012 | 0.023 | 0.044 | -0.002 | 0.013 | 0.011 | 0.017 | -0.001 | 0.005 | 0.017 | 0.026 |
| rNMP EF | *R* | rA | 0.798 | 0.582 | 0.540 | 0.840 | -0.186 | -0.400 | 0.611 | 0.716 | -0.360 | -0.458 | 0.649 | 0.862 |
| | | rC | 0.815 | 0.886 | 0.961 | 0.929 | -0.720 | 0.398 | 0.615 | 0.520 | 0.066 | 0.665 | 0.739 | 0.763 |
| | | rG | 0.613 | 0.820 | 0.786 | 0.940 | -0.919 | 0.376 | 0.315 | 0.651 | -0.724 | 0.349 | 0.458 | 0.736 |
| | | rU | 0.588 | 0.667 | 0.889 | 0.760 | -0.736 | 0.868 | 0.720 | 0.474 | -0.206 | 0.617 | 0.507 | 0.769 |
| | *p*-value | rA | 5.71E-03 | 7.75E-02 | 1.07E-01 | 2.37E-03 | 6.07E-01 | 2.52E-01 | 6.06E-02 | 2.00E-02 | 3.07E-01 | 1.83E-01 | 4.23E-02 | 1.33E-03 |
| | | rC | 4.03E-03 | 6.46E-04 | 9.47E-06 | 1.04E-04 | 1.88E-02 | 2.54E-01 | 5.85E-02 | 1.23E-01 | 8.57E-01 | 3.58E-02 | 1.47E-02 | 1.03E-02 |
| | | rG | 5.94E-02 | 3.64E-03 | 7.01E-03 | 5.18E-05 | 1.74E-04 | 2.85E-01 | 3.75E-01 | 4.17E-02 | 1.80E-02 | 3.23E-01 | 1.83E-01 | 1.53E-02 |
| | | rU | 7.35E-02 | 3.52E-02 | 5.83E-04 | 1.07E-02 | 1.52E-02 | 1.13E-03 | 1.88E-02 | 1.67E-01 | 5.68E-01 | 5.76E-02 | 1.35E-01 | 9.40E-03 |
| | slope | rA | 0.060 | 0.018 | 0.026 | 0.050 | -0.008 | -0.013 | 0.040 | 0.024 | -0.025 | -0.017 | 0.041 | 0.039 |
| | | rC | 0.143 | 0.056 | 0.165 | 0.095 | -0.072 | 0.017 | 0.057 | 0.037 | 0.006 | 0.035 | 0.059 | 0.065 |
| | | rG | 0.105 | 0.087 | 0.171 | 0.181 | -0.073 | 0.024 | 0.028 | 0.064 | -0.038 | 0.022 | 0.029 | 0.067 |

| | | | | | | | | | | | | | | |
|---|---|---|---|---|---|---|---|---|---|---|---|---|---|---|
| | | **rU** | 0.075 | 0.049 | 0.168 | 0.064 | -0.040 | 0.050 | 0.056 | 0.021 | -0.011 | 0.029 | 0.041 | 0.045 |
| **rNMP Percentage** | ***R*** | **rA** | 0.358 | -0.762 | -0.121 | -0.792 | 0.684 | -0.408 | 0.063 | -0.467 | 0.644 | -0.573 | -0.010 | -0.664 |
| | | **rC** | 0.023 | 0.062 | 0.134 | -0.688 | -0.320 | 0.424 | 0.846 | -0.239 | 0.031 | 0.318 | -0.164 | 0.456 |
| | | **rG** | -0.467 | 0.029 | 0.022 | 0.906 | -0.341 | -0.522 | -0.353 | 0.710 | -0.486 | 0.378 | 0.239 | 0.456 |
| | | **rU** | 0.241 | 0.455 | -0.196 | -0.633 | 0.280 | 0.521 | -0.423 | -0.524 | -0.083 | 0.019 | -0.014 | 0.127 |
| | ***p*-value** | **rA** | 3.09E-01 | 1.04E-02 | 7.39E-01 | 6.30E-03 | 2.90E-02 | 2.42E-01 | 8.62E-01 | 1.73E-01 | 4.46E-02 | 8.34E-02 | 9.78E-01 | 3.63E-02 |
| | | **rC** | 9.50E-01 | 8.65E-01 | 7.13E-01 | 2.79E-02 | 3.68E-01 | 2.22E-01 | 2.05E-03 | 5.06E-01 | 9.32E-01 | 3.70E-01 | 6.51E-01 | 1.85E-01 |
| | | **rG** | 1.74E-01 | 9.36E-01 | 9.51E-01 | 3.00E-04 | 3.34E-01 | 1.21E-01 | 3.17E-01 | 2.15E-02 | 1.55E-01 | 2.82E-01 | 5.06E-01 | 1.85E-01 |
| | | **rU** | 5.02E-01 | 1.86E-01 | 5.87E-01 | 4.92E-02 | 4.33E-01 | 1.23E-01 | 2.23E-01 | 1.20E-01 | 8.20E-01 | 9.58E-01 | 9.70E-01 | 7.27E-01 |
| | **slope** | **rA** | 0.291 | -0.409 | -0.053 | -0.190 | 0.715 | -0.358 | 0.049 | -0.215 | 0.839 | -0.641 | -0.006 | -0.311 |
| | | **rC** | 0.019 | 0.049 | 0.107 | -0.312 | -0.578 | 0.455 | 0.482 | -0.069 | 0.033 | 0.322 | -0.116 | 0.129 |
| | | **rG** | -0.506 | 0.019 | 0.013 | 0.533 | -0.411 | -0.493 | -0.225 | 0.363 | -0.805 | 0.301 | 0.131 | 0.174 |
| | | **rU** | 0.195 | 0.341 | -0.068 | -0.031 | 0.274 | 0.396 | -0.306 | -0.078 | -0.068 | 0.018 | -0.009 | 0.008 |
| **Template Strand** | | | | | | | | | | | | | | |
| | **Stats** | **Ribo** | **0-1 kb down stream of TSS** | | | | **4-5 kb down stream of TSS** | | | | **3-2 kb up stream of TSS** | | | |
| | | | **CD4$^+$T** | **hESC-H9** | **HEK WT** | **KO avg** | **CD4$^+$T** | **hESC-H9** | **HEK WT** | **KO avg** | **CD4$^+$T** | **hESC-H9** | **HEK WT** | **KO avg** |
| **rNMP counts** | ***R*** | **rA** | 0.495 | 0.645 | 0.893 | 0.901 | -0.140 | 0.676 | 0.942 | 0.854 | -0.205 | 0.667 | 0.730 | 0.736 |
| | | **rC** | 0.356 | 0.789 | 0.967 | 0.966 | -0.667 | 0.713 | 0.656 | 0.776 | -0.238 | 0.905 | 0.250 | 0.712 |
| | | **rG** | 0.023 | 0.275 | 0.794 | 0.909 | -0.718 | 0.839 | 0.772 | 0.781 | -0.215 | 0.625 | 0.844 | 0.802 |
| | | **rU** | 0.606 | -0.180 | 0.743 | 0.828 | 0.211 | 0.643 | 0.651 | 0.843 | 0.632 | 0.608 | 0.744 | 0.790 |
| | ***p*-value** | **rA** | 1.46E-01 | 4.39E-02 | 5.01E-04 | 3.75E-04 | 7.00E-01 | 3.19E-02 | 4.61E-05 | 1.66E-03 | 5.70E-01 | 3.50E-02 | 1.65E-02 | 1.53E-02 |
| | | **rC** | 3.13E-01 | 6.69E-03 | 5.17E-06 | 5.49E-06 | 3.52E-02 | 2.07E-02 | 3.94E-02 | 8.28E-03 | 5.08E-01 | 3.22E-04 | 4.87E-01 | 2.09E-02 |
| | | **rG** | 9.50E-01 | 4.41E-01 | 6.05E-03 | 2.67E-04 | 1.95E-02 | 2.40E-03 | 8.93E-03 | 7.71E-03 | 5.52E-01 | 5.31E-02 | 2.12E-03 | 5.25E-03 |
| | | **rU** | 6.33E-02 | 6.18E-01 | 1.37E-02 | 3.08E-03 | 5.58E-01 | 4.49E-02 | 4.13E-02 | 2.22E-03 | 4.99E-02 | 6.23E-02 | 1.37E-02 | 6.55E-03 |
| | **slope** | **rA** | 0.008 | 0.009 | 0.029 | 0.317 | -0.001 | 0.019 | 0.023 | 0.274 | -0.003 | 0.016 | 0.014 | 0.172 |
| | | **rC** | 0.007 | 0.037 | 0.073 | 0.867 | -0.014 | 0.020 | 0.024 | 0.423 | -0.003 | 0.031 | 0.007 | 0.274 |
| | | **rG** | 0.000 | 0.006 | 0.029 | 0.605 | -0.010 | 0.024 | 0.019 | 0.459 | -0.005 | 0.017 | 0.018 | 0.398 |
| | | **rU** | 0.008 | -0.002 | 0.008 | 0.036 | 0.002 | 0.011 | 0.007 | 0.032 | 0.005 | 0.011 | 0.011 | 0.026 |
| **rNMP EF** | ***R*** | **rA** | 0.492 | 0.617 | 0.751 | 0.934 | -0.156 | 0.846 | 0.655 | 0.835 | -0.063 | 0.618 | 0.526 | 0.789 |
| | | **rC** | 0.142 | 0.794 | 0.890 | 0.967 | -0.566 | 0.678 | 0.688 | 0.736 | 0.009 | 0.819 | 0.402 | 0.706 |
| | | **rG** | 0.577 | 0.135 | 0.670 | 0.958 | -0.660 | 0.931 | 0.456 | 0.765 | -0.017 | 0.514 | 0.591 | 0.767 |
| | | **rU** | 0.481 | 0.320 | 0.742 | 0.797 | 0.310 | 0.617 | 0.664 | 0.733 | -0.219 | 0.237 | 0.327 | 0.903 |
| | ***p*-value** | **rA** | 1.48E-01 | 5.74E-02 | 1.23E-02 | 7.46E-05 | 6.67E-01 | 2.01E-03 | 3.98E-02 | 2.66E-03 | 8.62E-01 | 5.68E-02 | 1.19E-01 | 6.67E-03 |
| | | **rC** | 6.95E-01 | 6.05E-03 | 5.54E-04 | 4.90E-06 | 8.82E-02 | 3.11E-02 | 2.79E-02 | 1.53E-02 | 9.81E-01 | 3.72E-03 | 2.49E-01 | 2.25E-02 |
| | | **rG** | 8.09E-02 | 7.11E-01 | 3.40E-02 | 1.29E-05 | 3.77E-02 | 8.91E-05 | 1.86E-01 | 9.93E-03 | 9.64E-01 | 1.29E-01 | 7.17E-02 | 9.62E-03 |
| | | **rU** | 1.60E-01 | 3.68E-01 | 1.41E-02 | 5.76E-03 | 3.84E-01 | 5.75E-02 | 3.63E-02 | 1.60E-02 | 5.43E-01 | 5.11E-01 | 3.56E-01 | 3.42E-04 |
| | **slope** | **rA** | 0.033 | 0.024 | 0.070 | 0.069 | -0.010 | 0.059 | 0.046 | 0.053 | -0.004 | 0.034 | 0.021 | 0.036 |
| | | **rC** | 0.021 | 0.106 | 0.191 | 0.168 | -0.066 | 0.048 | 0.047 | 0.061 | 0.001 | 0.066 | 0.029 | 0.041 |
| | | **rG** | 0.045 | 0.009 | 0.111 | 0.125 | -0.053 | 0.076 | 0.034 | 0.079 | -0.002 | 0.043 | 0.038 | 0.073 |
| | | **rU** | 0.050 | 0.016 | 0.059 | 0.054 | 0.026 | 0.055 | 0.050 | 0.043 | -0.014 | 0.015 | 0.019 | 0.035 |
| **rNMP Percentage** | ***R*** | **rA** | 0.089 | -0.158 | 0.295 | -0.228 | 0.710 | -0.140 | -0.214 | -0.400 | -0.204 | -0.377 | -0.257 | -0.632 |
| | | **rC** | 0.002 | 0.390 | 0.511 | 0.425 | -0.884 | 0.141 | 0.096 | 0.445 | 0.061 | 0.716 | 0.024 | -0.637 |
| | | **rG** | -0.482 | -0.138 | -0.389 | -0.183 | -0.259 | 0.530 | 0.297 | 0.253 | -0.716 | -0.278 | 0.284 | 0.876 |
| | | **rU** | 0.379 | -0.328 | -0.762 | -0.713 | 0.601 | -0.520 | -0.226 | -0.570 | 0.662 | -0.253 | -0.106 | 0.175 |
| | ***p*-value** | **rA** | 8.08E-01 | 6.63E-01 | 4.09E-01 | 5.27E-01 | 2.13E-02 | 7.00E-01 | 5.52E-01 | 2.52E-01 | 5.71E-01 | 2.83E-01 | 4.73E-01 | 4.99E-02 |
| | | **rC** | 9.95E-01 | 2.65E-01 | 1.31E-01 | 2.21E-01 | 6.78E-04 | 6.98E-01 | 7.92E-01 | 1.97E-01 | 8.67E-01 | 1.99E-02 | 9.49E-01 | 4.75E-02 |
| | | **rG** | 1.59E-01 | 7.05E-01 | 2.67E-01 | 6.13E-01 | 4.71E-01 | 1.15E-01 | 4.04E-01 | 4.81E-01 | 2.00E-02 | 4.36E-01 | 4.27E-01 | 8.84E-04 |
| | | **rU** | 2.80E-01 | 3.55E-01 | 1.04E-02 | 2.07E-02 | 6.61E-02 | 1.23E-01 | 5.30E-01 | 8.55E-02 | 3.69E-02 | 4.81E-01 | 7.71E-01 | 6.28E-01 |
| | **slope** | **rA** | 0.056 | -0.085 | 0.115 | -0.056 | 0.784 | -0.068 | -0.163 | -0.162 | -0.195 | -0.336 | -0.236 | -0.223 |
| | | **rC** | 0.002 | 0.386 | 0.314 | 0.145 | -1.131 | 0.078 | 0.099 | 0.187 | 0.037 | 0.756 | 0.018 | -0.248 |
| | | **rG** | -0.420 | -0.105 | -0.218 | -0.039 | -0.209 | 0.300 | 0.173 | 0.067 | -0.545 | -0.229 | 0.268 | 0.455 |
| | | **rU** | 0.361 | -0.196 | -0.211 | -0.050 | 0.556 | -0.310 | -0.109 | -0.092 | 0.702 | -0.192 | -0.050 | 0.016 |

**table S10.**

One-sided Mann-Whitney $U$-test $p$-values for mono (R) and upstream dinucleotide (NR) frequencies in the following regions of the human nuclear genome: whole nuclear genome, near (0-500 bp) and away (4-5 kb) from TSS, in CpG islands and inter-CpG regions specific to data presented in **Fig. 5C-E**.

| **Nucleotide** | **Whole genome** | | | | | **CpG islands** | | **Inter-CpG** | | **TSS, +/- 0-500 bp** | | **TSS, +/- 4-5 kb** | |
|---|---|---|---|---|---|---|---|---|---|---|---|---|---|
| R or NR | $CD4^+$ T | hESC-H9 | HEK293T-WT | HEK293T-RNH2A-KO-T3-8 | HEK293T-RNH2A-KO-T3-17 | HEK293T-RNH2A-KO-T3-8 | HEK293T-RNH2A-KO-T3-17 | HEK293T-RNH2A-KO-T3-8 | HEK293T-RNH2A-KO-T3-17 | HEK293T-RNH2A-KO-T3-8 | HEK293T-RNH2A-KO-T3-17 | HEK293T-RNH2A-KO-T3-8 | HEK293T-RNH2A-KO-T3-17 |
| A | 0.353 | 0.867 | 0.963 | 0.998 | 0.748 | 0.998 | 0.963 | 0.998 | 0.748 | 0.748 | 0.004 | 0.998 | 0.748 |
| C | 0.681 | 0.025 | 0.004 | 0.004 | 0.004 | 0.004 | 0.004 | 0.004 | 0.004 | 0.004 | 0.004 | 0.004 | 0.004 |
| G | 0.681 | 0.174 | 0.328 | 0.004 | 0.004 | 0.004 | 0.004 | 0.004 | 0.004 | 0.004 | 0.004 | 0.004 | 0.004 |
| U | 1.000 | 0.999 | 0.998 | 0.998 | 0.998 | 0.998 | 0.998 | 0.998 | 0.998 | 0.998 | 0.998 | 0.998 | 0.998 |
| AA | 0.353 | 0.025 | 0.328 | 0.004 | 0.004 | 0.004 | 0.004 | 0.004 | 0.004 | 0.004 | 0.004 | 0.004 | 0.004 |
| CA | 0.985 | 0.984 | 0.059 | 0.004 | 0.004 | 0.004 | 0.004 | 0.004 | 0.004 | 0.004 | 0.004 | 0.004 | 0.004 |
| GA | 0.110 | 0.174 | 0.748 | 0.059 | 0.328 | 0.059 | 0.059 | 0.059 | 0.059 | 0.998 | 0.998 | 0.059 | 0.059 |
| TA | 0.907 | 0.999 | 0.963 | 0.998 | 0.998 | 0.998 | 0.998 | 0.998 | 0.998 | 0.998 | 0.998 | 0.998 | 0.998 |
| AC | 0.019 | 0.001 | 0.963 | 0.004 | 0.004 | 0.004 | 0.004 | 0.004 | 0.004 | 0.004 | 0.004 | 0.004 | 0.004 |
| CC | 0.907 | 0.534 | 0.059 | 0.004 | 0.004 | 0.004 | 0.004 | 0.004 | 0.004 | 0.004 | 0.004 | 0.004 | 0.004 |
| GC | 0.999 | 0.999 | 0.998 | 0.998 | 0.998 | 0.998 | 0.998 | 0.998 | 0.998 | 0.998 | 0.998 | 0.998 | 0.998 |
| TC | 0.681 | 0.867 | 0.748 | 0.998 | 0.998 | 0.998 | 0.998 | 0.998 | 0.998 | 0.998 | 0.998 | 0.998 | 0.998 |
| AG | 0.000 | 0.174 | 0.328 | 0.004 | 0.004 | 0.004 | 0.004 | 0.004 | 0.004 | 0.004 | 0.004 | 0.004 | 0.004 |
| CG | 0.985 | 0.867 | 0.059 | 0.059 | 0.059 | 0.004 | 0.004 | 0.004 | 0.004 | 0.328 | 0.328 | 0.004 | 0.004 |
| GG | 0.110 | 0.534 | 0.998 | 0.748 | 0.963 | 0.998 | 0.998 | 0.998 | 0.963 | 0.998 | 0.998 | 0.998 | 0.963 |
| TG | 1.000 | 0.999 | 0.998 | 0.998 | 0.998 | 0.998 | 0.998 | 0.998 | 0.998 | 0.998 | 0.998 | 0.998 | 0.998 |
| AU | 0.002 | 0.025 | 0.748 | 0.998 | 0.998 | 0.998 | 0.998 | 0.998 | 0.998 | 0.998 | 0.998 | 0.998 | 0.998 |
| CU | 0.353 | 0.867 | 0.328 | 0.004 | 0.004 | 0.004 | 0.004 | 0.004 | 0.004 | 0.004 | 0.004 | 0.004 | 0.004 |
| GU | 0.681 | 0.984 | 0.963 | 0.998 | 0.998 | 0.998 | 0.998 | 0.998 | 0.998 | 0.998 | 0.998 | 0.998 | 0.998 |
| TU | 0.999 | 0.999 | 0.748 | 0.998 | 0.998 | 0.998 | 0.998 | 0.998 | 0.998 | 0.998 | 0.998 | 0.998 | 0.998 |

**table S11.**

One-sided Mann-Whitney *U*-test *p*-values for downstream dinucleotide (RN) frequencies in whole nuclear genome specific to data presented in **fig. S10C.**

| **Nuc** | **Whole Genome** | | | | |
|---|---|---|---|---|---|
| RN | $CD4^{+}T$ | hESC-H9 | HEK293T | HEK293T-RNH2A-KO-T3-8 | HEK293T-RNH2A-KO-T3-17 |
| A | 0.999 | 0.867 | 0.963 | 0.998 | 0.998 |
| C | 0.907 | 0.867 | 0.963 | 0.004 | 0.004 |
| G | 0.353 | 0.534 | 0.748 | 0.059 | 0.328 |
| U | 0.002 | 0.174 | 0.059 | 0.004 | 0.004 |
| AA | 1.000 | 0.984 | 0.998 | 0.748 | 0.748 |
| AC | 0.110 | 0.867 | 0.748 | 0.998 | 0.998 |
| AG | 0.353 | 0.174 | 0.748 | 0.004 | 0.004 |
| AT | 0.999 | 0.867 | 0.748 | 0.748 | 0.748 |
| CA | 1.000 | 0.984 | 0.998 | 0.998 | 0.998 |
| CC | 0.019 | 0.867 | 0.059 | 0.004 | 0.004 |
| CG | 0.002 | 0.001 | 0.059 | 0.004 | 0.004 |
| CT | 1.000 | 0.999 | 0.748 | 0.998 | 0.998 |
| GA | 0.999 | 0.984 | 0.998 | 0.998 | 0.998 |
| GC | 0.110 | 0.025 | 0.748 | 0.998 | 0.998 |
| GG | 0.000 | 0.025 | 0.004 | 0.004 | 0.004 |
| GT | 0.907 | 0.984 | 0.328 | 0.004 | 0.004 |
| UA | 0.999 | 0.867 | 0.963 | 0.998 | 0.998 |
| UC | 0.907 | 0.867 | 0.963 | 0.004 | 0.004 |
| UG | 0.353 | 0.534 | 0.748 | 0.059 | 0.328 |
| UT | 0.002 | 0.174 | 0.059 | 0.004 | 0.004 |

**table S12.**

One-sided Mann Whitney *U*-test *p*-values for trinucleotides (NNR, NRN and RNN) for whole nuclear genome specific to data presented in **fig. S11.**

| Whole Genome | | | | | | | | |
|---|---|---|---|---|---|---|---|---|
| NNR | RNH2A-KO-T3-8 | RNH2A-KO-T3-17 | NRN | RNH2A-KO-T3-8 | RNH2A-KO-T3-17 | RNN | RNH2A-KO-T3-8 | RNH2A-KO-T3-17 |
| AAA | 0.998 | 0.998 | AAA | 0.963 | 0.963 | AAA | 0.998 | 0.998 |
| CAA | 0.004 | 0.004 | AAC | 0.328 | 0.328 | AAC | 0.998 | 0.998 |
| GAA | 0.004 | 0.004 | AAG | 0.004 | 0.004 | AAG | 0.998 | 0.998 |
| TAA | 0.748 | 0.748 | AAT | 0.004 | 0.004 | AAT | 0.998 | 0.998 |
| ACA | 0.004 | 0.004 | CAA | 0.998 | 0.998 | ACA | 0.998 | 0.998 |
| CCA | 0.004 | 0.059 | CAC | 0.004 | 0.059 | ACC | 0.748 | 0.748 |
| GCA | 0.004 | 0.004 | CAG | 0.004 | 0.004 | ACG | 0.748 | 0.748 |
| TCA | 0.059 | 0.004 | CAT | 0.004 | 0.004 | ACT | 0.998 | 0.998 |
| AGA | 0.328 | 0.748 | GAA | 0.998 | 0.998 | AGA | 0.004 | 0.004 |
| CGA | 0.059 | 0.059 | GAC | 0.998 | 0.963 | AGC | 0.004 | 0.004 |
| GGA | 0.059 | 0.004 | GAG | 0.004 | 0.004 | AGG | 0.004 | 0.004 |
| TGA | 0.998 | 0.998 | GAT | 0.004 | 0.004 | AGT | 0.004 | 0.004 |
| ATA | 0.998 | 0.998 | TAA | 0.998 | 0.998 | ATA | 0.963 | 0.748 |
| CTA | 0.998 | 0.998 | TAC | 0.998 | 0.998 | ATC | 0.004 | 0.004 |
| GTA | 0.998 | 0.998 | TAG | 0.998 | 0.998 | ATG | 0.004 | 0.004 |
| TTA | 0.998 | 0.998 | TAT | 0.998 | 0.998 | ATT | 0.059 | 0.004 |
| AAC | 0.998 | 0.998 | ACA | 0.059 | 0.748 | CAA | 0.998 | 0.998 |
| CAC | 0.004 | 0.004 | ACC | 0.004 | 0.004 | CAC | 0.963 | 0.998 |
| GAC | 0.004 | 0.004 | ACG | 0.059 | 0.059 | CAG | 0.998 | 0.998 |
| TAC | 0.059 | 0.328 | ACT | 0.004 | 0.059 | CAT | 0.998 | 0.998 |
| ACC | 0.004 | 0.004 | CCA | 0.004 | 0.004 | CCA | 0.004 | 0.004 |
| CCC | 0.004 | 0.004 | CCC | 0.004 | 0.004 | CCC | 0.004 | 0.059 |
| GCC | 0.004 | 0.004 | CCG | 0.004 | 0.004 | CCG | 0.004 | 0.004 |
| TCC | 0.004 | 0.004 | CCT | 0.004 | 0.004 | CCT | 0.328 | 0.328 |
| AGC | 0.998 | 0.998 | GCA | 0.998 | 0.998 | CGA | 0.998 | 0.963 |
| CGC | 0.328 | 0.963 | GCC | 0.998 | 0.998 | CGC | 0.004 | 0.328 |
| GGC | 0.998 | 0.998 | GCG | 0.998 | 0.998 | CGG | 0.059 | 0.328 |
| TGC | 0.998 | 0.998 | GCT | 0.998 | 0.998 | CGT | 0.963 | 0.998 |
| ATC | 0.998 | 0.998 | TCA | 0.998 | 0.998 | CTA | 0.963 | 0.748 |
| CTC | 0.998 | 0.998 | TCC | 0.998 | 0.998 | CTC | 0.004 | 0.004 |
| GTC | 0.998 | 0.998 | TCG | 0.998 | 0.998 | CTG | 0.004 | 0.004 |
| TTC | 0.998 | 0.998 | TCT | 0.998 | 0.998 | CTT | 0.748 | 0.004 |
| AAG | 0.998 | 0.998 | AGA | 0.059 | 0.059 | GAA | 0.998 | 0.998 |
| CAG | 0.004 | 0.004 | AGC | 0.328 | 0.748 | GAC | 0.328 | 0.328 |
| GAG | 0.059 | 0.004 | AGG | 0.004 | 0.004 | GAG | 0.328 | 0.328 |
| TAG | 0.004 | 0.004 | AGT | 0.004 | 0.004 | GAT | 0.998 | 0.998 |
| ACG | 0.004 | 0.004 | CGA | 0.004 | 0.004 | GCA | 0.998 | 0.998 |
| CCG | 0.004 | 0.004 | CGC | 0.004 | 0.004 | GCC | 0.748 | 0.748 |
| GCG | 0.059 | 0.963 | CGG | 0.004 | 0.004 | GCG | 0.059 | 0.004 |
| TCG | 0.004 | 0.004 | CGT | 0.004 | 0.004 | GCT | 0.998 | 0.998 |
| AGG | 0.998 | 0.998 | GGA | 0.963 | 0.748 | GGA | 0.004 | 0.004 |
| CGG | 0.004 | 0.059 | GGC | 0.998 | 0.998 | GGC | 0.004 | 0.004 |
| GGG | 0.059 | 0.004 | GGG | 0.004 | 0.004 | GGG | 0.004 | 0.004 |
| TGG | 0.963 | 0.963 | GGT | 0.328 | 0.748 | GGT | 0.004 | 0.004 |

| | | | | | | | | |
|---|---|---|---|---|---|---|---|---|
| ATG | 0.998 | 0.998 | TGA | 0.998 | 0.998 | GTA | 0.998 | 0.963 |
| CTG | 0.998 | 0.998 | TGC | 0.998 | 0.998 | GTC | 0.004 | 0.004 |
| GTG | 0.998 | 0.998 | TGG | 0.998 | 0.998 | GTG | 0.004 | 0.004 |
| TTG | 0.998 | 0.998 | TGT | 0.998 | 0.998 | GTT | 0.998 | 0.963 |
| AAU | 0.998 | 0.998 | AUA | 0.998 | 0.998 | UAA | 0.998 | 0.998 |
| CAU | 0.998 | 0.998 | AUC | 0.998 | 0.998 | UAC | 0.998 | 0.998 |
| GAU | 0.998 | 0.998 | AUG | 0.998 | 0.998 | UAG | 0.998 | 0.998 |
| TAU | 0.998 | 0.998 | AUT | 0.998 | 0.998 | UAT | 0.998 | 0.998 |
| ACU | 0.004 | 0.004 | CUA | 0.004 | 0.004 | UCA | 0.004 | 0.004 |
| CCU | 0.004 | 0.004 | CUC | 0.004 | 0.004 | UCC | 0.004 | 0.004 |
| GCU | 0.004 | 0.004 | CUG | 0.004 | 0.004 | UCG | 0.004 | 0.004 |
| TCU | 0.004 | 0.004 | CUT | 0.004 | 0.004 | UCT | 0.004 | 0.004 |
| AGU | 0.998 | 0.998 | GUA | 0.998 | 0.998 | UGA | 0.998 | 0.998 |
| CGU | 0.998 | 0.998 | GUC | 0.998 | 0.998 | UGC | 0.004 | 0.004 |
| GGU | 0.998 | 0.998 | GUG | 0.998 | 0.998 | UGG | 0.328 | 0.059 |
| TGU | 0.998 | 0.998 | GUT | 0.998 | 0.998 | UGT | 0.963 | 0.963 |
| ATU | 0.998 | 0.998 | TUA | 0.998 | 0.998 | UTA | 0.998 | 0.998 |
| CTU | 0.998 | 0.998 | TUC | 0.998 | 0.998 | UTC | 0.963 | 0.998 |
| GTU | 0.998 | 0.998 | TUG | 0.998 | 0.998 | UTG | 0.998 | 0.998 |
| TTU | 0.998 | 0.998 | TUT | 0.998 | 0.998 | UTT | 0.998 | 0.998 |

**table S13.**

Two-sided unpaired and paired Mann-Whitney *U*-test *p*-values for mono (R) and upstream dinucleotide (NR) frequencies in regions of the human nuclear genome. Significant comparisons between HEK293T-RNH2A-KO libraries (two-tailed Mann–Whitney *U*-test) are marked with asterisks in **Fig. 5D-G, fig. S12**. (*). Significance levels: * ($0.05 \geq p > 0.01$), ** ($0.01 \geq p > 0.001$), *** ($p < 0.001$).

| Nucleotide (R or NR) | CpG Islands vs Inter-CpG | | TSS, +/- 0-500 bp vs +/- 4-5 kb | | TSS, high expression 0-500 bp, template vs non-template strand | | TSS, low expression 0-500 bp, template vs non-template strand | |
|---|---|---|---|---|---|---|---|---|
| | unpaired | paired | unpaired | paired | unpaired | paired | unpaired | paired |
| A | 0.019 | 0.002 | 0.247 | 0.002 | 0.005 | 0.002 | 0.353 | 0.002 |
| C | 0.052 | 0.002 | 0.684 | 0.002 | 0.123 | 0.002 | 0.853 | 0.322 |
| G | 0.912 | 1.000 | 0.353 | 0.004 | 0.912 | 0.084 | 0.481 | 0.002 |
| U | 0 | 0.002 | 0 | 0.002 | 0.005 | 0.014 | 0.436 | 0.105 |
| AA | 0.001 | 0.002 | 0.436 | 0.010 | 0.023 | 0.004 | 0.436 | 0.084 |
| CA | 0.075 | 0.002 | 0.796 | 0.049 | 0.005 | 0.002 | 0.353 | 0.010 |
| GA | 0.002 | 0.002 | 0.043 | 0.002 | 0.481 | 0.432 | 0.684 | 0.275 |
| TA | 0 | 0.002 | 0.353 | 0.084 | 0.853 | 0.846 | 0.684 | 0.432 |
| AC | 0.003 | 0.002 | 0.19 | 0.004 | 0 | 0.002 | 0.579 | 0.193 |
| CC | 0.004 | 0.002 | 0.393 | 0.020 | 0 | 0.002 | 0.631 | 0.064 |
| GC | 0 | 0.002 | 0.004 | 0.002 | 0.529 | 0.014 | 0.353 | 0.020 |
| TC | 0 | 0.002 | 0 | 0.002 | 0.436 | 0.232 | 0.089 | 0.375 |
| AG | 0 | 0.002 | 0 | 0.002 | 0.247 | 0.002 | 0.971 | 0.846 |
| CG | 0 | 0.002 | 0 | 0.002 | 0.002 | 0.002 | 0.353 | 0.131 |
| GG | 0.019 | 0.002 | 0.043 | 0.002 | 0.165 | 0.004 | 0.143 | 0.064 |
| TG | 0 | 0.002 | 0 | 0.002 | 0.015 | 0.020 | 0.247 | 0.105 |
| AU | 0.105 | 0.084 | 0.481 | 0.232 | 0.481 | 0.557 | 0.912 | 0.770 |
| CU | 0.023 | 0.002 | 0.529 | 0.492 | 0.123 | 0.131 | 0.853 | 0.492 |
| GU | 0.631 | 0.105 | 0.631 | 0.625 | 0.043 | 0.064 | 0.631 | 0.770 |
| TU | 0 | 0.002 | 0.035 | 0.064 | 0.19 | 0.049 | 0.043 | 0.020 |

**table S14.**

One-sided Mann Whitney *U*-test *p*-values for mono (R) and upstream dinucleotide (NR) frequencies near high and low expression TSS for template and non-template strand, specific to data presented in **Fig. 5F-G**.

| **Nucleotide** | **TSS, high expression 0-500 bp, template strand** | | **TSS, high expression 0-500 bp, non-template strand** | | **TSS, low expression 0-500 bp, template strand** | | **TSS, low expression 0-500 bp, non-template strand** | |
|---|---|---|---|---|---|---|---|---|
| R or NR | RNH2A-KO-T3-8 | RNH2A-KO-T3-17 | RNH2A-KO-T3-8 | RNH2A-KO-T3-17 | RNH2A-KO-T3-8 | RNH2A-KO-T3-17 | RNH2A-KO-T3-8 | RNH2A-KO-T3-17 |
| A | 0.998 | 0.748 | 0.059 | 0.004 | 0.998 | 0.748 | 0.998 | 0.748 |
| C | 0.004 | 0.004 | 0.004 | 0.004 | 0.004 | 0.004 | 0.004 | 0.004 |
| G | 0.004 | 0.004 | 0.004 | 0.004 | 0.004 | 0.004 | 0.004 | 0.004 |
| U | 0.998 | 0.998 | 0.998 | 0.998 | 0.998 | 0.998 | 0.998 | 0.998 |
| AA | 0.004 | 0.004 | 0.004 | 0.004 | 0.059 | 0.059 | 0.004 | 0.059 |
| CA | 0.004 | 0.004 | 0.004 | 0.004 | 0.004 | 0.004 | 0.004 | 0.004 |
| GA | 0.998 | 0.998 | 0.998 | 0.998 | 0.059 | 0.059 | 0.059 | 0.059 |
| TA | 0.998 | 0.998 | 0.998 | 0.998 | 0.998 | 0.998 | 0.998 | 0.998 |
| AC | 0.004 | 0.004 | 0.004 | 0.004 | 0.004 | 0.004 | 0.004 | 0.004 |
| CC | 0.004 | 0.004 | 0.004 | 0.004 | 0.004 | 0.004 | 0.004 | 0.004 |
| GC | 0.998 | 0.998 | 0.998 | 0.998 | 0.998 | 0.998 | 0.998 | 0.998 |
| TC | 0.998 | 0.998 | 0.998 | 0.998 | 0.998 | 0.998 | 0.998 | 0.998 |
| AG | 0.004 | 0.004 | 0.004 | 0.004 | 0.004 | 0.004 | 0.004 | 0.004 |
| CG | 0.328 | 0.328 | 0.998 | 0.998 | 0.004 | 0.004 | 0.004 | 0.004 |
| GG | 0.998 | 0.998 | 0.963 | 0.963 | 0.963 | 0.748 | 0.328 | 0.748 |
| TG | 0.998 | 0.998 | 0.998 | 0.998 | 0.998 | 0.998 | 0.998 | 0.998 |
| AU | 0.998 | 0.998 | 0.998 | 0.963 | 0.998 | 0.998 | 0.998 | 0.963 |
| CU | 0.004 | 0.004 | 0.004 | 0.004 | 0.004 | 0.004 | 0.004 | 0.004 |
| GU | 0.998 | 0.998 | 0.998 | 0.998 | 0.998 | 0.998 | 0.998 | 0.998 |

**table S15.**

Telomere rNMP EF estimation. For each library fragmented using the dsDNA Fragmentase, estimated telomere length using DNA-seq read counts, rNMP counts in telomers, and respective enrichment in telomeres are listed. The latter is also represented in **Fig 6C.**

| Library | Gender | Total genomic reads from DNA-seq | Total telomeric reads from DNA-seq | Genome length (bp) | rNMPs in nuclear genome (ribose-seq) | Est. telomere length (bp) | Total rNMPs in telomeres | rNMP EF | rNMPs on G-strand | rNMPs on C-strand | G/C strand ratio |
|---|---|---|---|---|---|---|---|---|---|---|---|
| FS185-CD4$^{+}$T | Male | 718,300,444 | 39,144 | 2,981,635,677 | 1,250,079 | 162,485 | 540 | 7.93 | 206 | 334 | 0.6 |
| FS197-hESC-H9 | Female | 764,564,708 | 67,729 | 3,031,042,417 | 969,423 | 268,505 | 851 | 9.91 | 717 | 134 | 5.4 |
| FS198-hESC-H9 | Female | 764,564,708 | 67,729 | 3,031,042,417 | 1,486,071 | 268,505 | 370 | 2.81 | 221 | 149 | 1.5 |
| FS326-HEK293T | Female | 817,573,193 | 30,083 | 3,031,042,417 | 271,552 | 111,529 | 74 | 7.41 | 64 | 10 | 6.4 |
| FS391-HEK293T | Female | 817,573,193 | 30,083 | 3,031,042,417 | 6,842,523 | 111,529 | 731 | 2.90 | 369 | 362 | 1.0 |
| FS327-RNH2A-KO T3-8 | Female | 671,977,196 | 14,599 | 3,031,042,417 | 5,990,007 | 65,851 | 759 | 5.83 | 93 | 666 | 0.4 |
| FS392-RNH2A-KO T3-8 | Female | 671,977,196 | 14,599 | 3,031,042,417 | 6,772,739 | 65,851 | 610 | 4.15 | 284 | 326 | 0.9 |
| FS329-RNH2A-KO T3-17 | Female | 841,702,458 | 36,037 | 3,031,042,417 | 16,502,973 | 129,772 | 904 | 1.28 | 355 | 549 | 0.6 |
| FS393-RNH2A-KO T3-17 | Female | 841,702,458 | 36,037 | 3,031,042,417 | 10,167,570 | 129,772 | 2,268 | 5.21 | 1,201 | 1,067 | 1.1 |

**table S16.**

Average relative frequency (*F*) for rNMP counts per 100 kb per million of genomic elements of CpG related regions, gene related regions, candidate-*cis* regulatory elements (cCREs), and telomeres. Avg *F*, standard deviation (SD), and standard error of mean (SEM) for each cell type is calculated using all libraries, except for telomeres, where rNMPs are detected only in libraries fragmented by the dsDNA Fragmentase. A schematic summary is represented in **Fig. 6D**, and **fig. S13.**

| **Annotation Set** | **Set elements** | **CD4$^+$T** | | | **hESC-H9** | | | **HEK293T** | | | **HEK293T-RNH2A-KO avg.** | | |
|---|---|---|---|---|---|---|---|---|---|---|---|---|---|
| | | **Average** | **SD** | **SEM** | **Average** | **SD** | **SEM** | **Average** | **SD** | **SEM** | **Average** | **SD** | **SEM** |
| CpG related regions | Island | 68 | 29 | 11 | 59 | 19 | 8 | 105 | 60 | 27 | 119 | 23 | 8 |
| | Shore | 49 | 8.8 | 3.2 | 50 | 5.3 | 2.2 | 56 | 11.7 | 5.2 | 66 | 6.9 | 2.5 |
| | Shelf | 44 | 4.3 | 1.8 | 45 | 2.7 | 1.1 | 46 | 3.3 | 1.5 | 52 | 3.9 | 1.4 |
| | Inter-CpG | 42 | 3.6 | 1.6 | 44 | 2.6 | 1.1 | 44 | 2.9 | 1.3 | 50 | 3.4 | 1.2 |
| Gene related regions | Enhancer | 45 | 3.7 | 1.4 | 45 | 2.5 | 1.0 | 46 | 3.8 | 1.7 | 50 | 3.1 | 1.1 |
| | Promoter | 60 | 14.1 | 5.3 | 52 | 8.7 | 3.6 | 74 | 32.5 | 14.5 | 78 | 9.4 | 3.3 |
| | 5' UTR | 64 | 20.4 | 7.5 | 51 | 10.7 | 4.4 | 84 | 46.2 | 20.7 | 90 | 15.3 | 5.4 |
| | First exon | 67 | 24.7 | 8.6 | 54 | 14.7 | 6.0 | 97 | 61.6 | 27.5 | 104 | 19.1 | 6.8 |
| | Proximal intron | 52 | 10.2 | 3.4 | 50 | 6.7 | 2.7 | 61 | 17.1 | 7.7 | 67 | 7.2 | 2.5 |
| | Internal exon(s) | 42 | 7.9 | 3.1 | 45 | 4.1 | 1.7 | 54 | 10.2 | 4.6 | 59 | 8.2 | 2.9 |
| | Distal intron(s) | 43 | 5.8 | 2.2 | 45 | 3.4 | 1.4 | 48 | 5.9 | 2.6 | 56 | 5.8 | 2.0 |
| | Last exon | 41 | 6.8 | 2.6 | 43 | 3.3 | 1.3 | 50 | 8.0 | 3.6 | 57 | 6.6 | 2.3 |
| | 3' UTR | 40 | 6.0 | 2.4 | 42 | 3.1 | 1.3 | 49 | 6.5 | 2.9 | 55 | 5.5 | 2.0 |
| Candidate - *cis* regulatory elements (cCREs) | dELS (H3K27ac) | 42 | 3.8 | 1.4 | 44 | 2.6 | 1.1 | 45 | 4.3 | 1.9 | 50 | 2.5 | 0.9 |
| | PLS (H3K4me3) | 70 | 22.9 | 8.2 | 57 | 13.6 | 5.5 | 97 | 55.2 | 24.7 | 101 | 16.1 | 5.7 |
| | pELS (H3K27ac) | 54 | 12.2 | 4.2 | 53 | 7.0 | 2.8 | 67 | 21.6 | 9.7 | 75 | 8.4 | 3.0 |
| | DNase-H3K4me3 | 41 | 4.1 | 1.5 | 42 | 2.9 | 1.2 | 44 | 5.4 | 2.4 | 50 | 3.3 | 1.2 |
| | CTCF-only | 36 | 0.7 | 0.3 | 38 | 2.0 | 0.8 | 37 | 3.2 | 1.4 | 38 | 1.1 | 0.4 |
| | Inter-cCRE | 43 | 4.6 | 1.5 | 44 | 2.2 | 0.9 | 48 | 7.3 | 3.2 | 52 | 3.2 | 1.1 |
| Telomeres | Telomeres | 266 | 0 | 0.00 | 210 | 117 | 47.81 | 170 | 74 | 33.22 | 136 | 58 | 18.21 |

**Supplementary figure legends**

**fig. S1. Abundant rNMPs in the human nuclear genome** (**A-D**) Four replicates of gel electrophoresis of heat shock and formamide denatured genomic DNA after no treatment and treatment with RNase HII. Ticks on the left edge of the gel image mark the positions of ladder bands, with corresponding molecular weights indicated in kilobases (kb). Lane M, 1 kb Plus DNA Ladder from NEB (CAT# N3200L); 1-5 left, genomic DNA from CD4$^+$T, hESC-H9, HEK293T, HEK293T RNH2A-KO-T3-8, or HEK293T RNH2A-KO-T3-17 not treated with RNase HII; 1-5 right, genomic DNA from the same cell types but treated with RNase HII. To the right of the gel image, percent intensity distributions are presented for untreated (left distribution panel) and treated (right panels) for 1. CD4$^+$T (orange), 2. hESC-H9 (green), 3. HEK293T (red), 4. HEK293T RNH2A-KO-T3-8 (pink) and 5. HEK293T RNH2A-KO-T3-17 (purple). Median fragment lengths are denoted for untreated (round, blue) and treated (triangle, brown) genomic DNA in the third panel and shown also in a table in A). See data in **table S1**. All panels are aligned with the analyzed gel image. Black horizontal lines on the right edge of the gel indicate the region analyzed, from the bottom of the wells to just above the end of the gel.

**fig. S2. rNMPs of the human nuclear ribome are found in CG rich sequences.** Heatmaps showing the relative frequency of mono and dinucleotide dNMP ±50 bp around rNMP-embedment sites, normalized to their genomic frequency (chr1 to chr22). Frequencies >1 (enriched) are shown in red, and <1 (depleted) in blue, with exact values indicated in each cell. Each column represents a different library. See data in **table S4**.

**fig. S3. rNMP embedment correlates with C/G-rich content and positive AT skew across genomic scales.** Heatmaps showing Pearson's correlation coefficient (R) for rNMP embedment frequencies with respect to genomic mono and dinucleotide fraction on the same strand of (**A**) 1 kb, (**B**) 10 kb, (**C**) 100 kb and (**D**) 1 Mb discrete bins spanning chr1- chr22. The cell types and the specific ribose-seq library names are shown under the heatmaps. Positive correlation is shown in green, negative correlation in purple, and no correlation in white. (**E**) Violin boxplots of rNMP Enrichment Factor (rNMP EF) (*y*-axis) with increasing A-T skew and G-C skew are shown in the top and bottom panels. The *x*-axis represents skew values grouped in 10 percentiles of data for 10 kb discrete bins in the genome, and labels represent mean skew value in each group. Darker brown represents positive skew, whereas lighter brown color represents negative skew.

**fig. S4. rNMPs around 1kb of CpG centers distant from TSS overlap with enhancer and intronic regions.** Heatmaps showing percentage of located within ±1 kb of CpG island centers that are more than 3 kb away from transcription start sites (TSSs), overlapping with candidate *cis*-regulatory elements (cCREs) and gene-related regions of protein-coding genes. Each column represents a ribose-seq library (labeled at the bottom), and each row corresponds to a genomic region category (labeled on the right). Percentages greater than 1% are shown in red, and less than 1% are shown in blue.

**fig. S5. No C/G strand bias in whole genome sequencing; rNMP-enriched zones (REZs) colocalize with R-loop regions.** (**A**) Karyotype-style line plot showing DNA enrichment factor (DNA EF, *y* axis) on the Watson (W, +) and Crick (C, -) strands across genomic coordinates (*x*-axis, in megabases, Mb) for chr1 – chr22 of DNA from CD4$^+$T cells. Fragmentation methods are color-coded: dsDNA Fragmentase (F) (red), RE1 (orange), RE2 (blue) and RE3 (green). CpG islands are shown in the center (transparent red bars: regions with high CpG island density appear as darker red, while regions with lower density appear lighter) (**B**) Karyotype-style line plots showing rNMP enrichment factor (rNMP EF, *y* axis) on the Watson (W, +) and Crick (C, -) strands with R-loop regions from RLBase (transparent orange bars) and (**C**) RIAN-seq (transparent purple bars) across genomic coordinates (*x*-axis, in megabases, Mb) for chr1 – chr22. Cell types are color-coded: CD4$^{+}$T (orange), hESC-H9 (green), HEK293T (red), HEK293T RNH2A-KO-T3-8 (pink), and HEK293T RNH2A-KO-T3-17 (purple).

**fig. S6. Properties of rNMP enrichment and distribution around gene regions.** (**A**) Line plots showing mean rNMP EF (*y*-axis) in 100-bp bins, ± 3 kb (*x*-axis) around TSSs and TTSs, for the non-template (top) and template (bottom) strands of non-coding genes. Cell types are color-coded: CD4$^{+}$T (orange), hESC-H9 (green), HEK293T (red), HEK293T RNH2A-KO-T3-8 (pink), and HEK293T RNH2A-KO-T3-17 (purple). (**B**) Violin boxplots of rNMP EF (*y*-axis) for regions ± 1 kb around TSS (red) and across gene bodies (TSS to TTS, blue) plotted against increasing gene expression levels, grouped into deciles (x-axis, left to right) for each cell type. (**C**) Bar plots showing rNMP EF in non-template (green) and template (purple) strands. Each dot represents a library for the corresponding cell type. Asterisks indicate significance levels from the Mann-Whitney *U*-test (*: $0.05 \geq p > 0.01$; **: $0.01 \geq 0.001$; ***: $0.001 \geq p$) (**D**) Line plots showing mean rNMP EF (*y*-axis) in 100 bp bins ± 3 kb (*x*-axis) around TSSs of protein coding genes, on the non-template (green) and template (purple) strands. Horizontal and vertical dotted lines indicate EF = 1 and TSS position, respectively.

**fig. S7. Association of rNMP enrichment with methylation data.** (**A**) Line plots showing methylation percentage (*y*-axis) in 100-bp bins, ± 3 kb (*x*-axis) around TSSs, for the non-template (top) and template (bottom) strands of non-coding genes. Cell types are color-coded: CD4$^{+}$T (orange), hESC-H9 (green), HEK293T (red), HEK293T RNH2A-KO-T3-8 (pink), and HEK293T RNH2A-KO-T3-17 (purple). (**B**) Violin boxplots of rNMP EF (*y*-axis) for regions ± 1 kb around TSS (red) plotted against increasing methylation percentages, grouped into deciles (*x*-axis, left to right) for each cell type.

**fig. S8**. **rC and rG embedment correlates more strongly with gene expression within 0–1 kb downstream of TSS than at upstream or distal downstream regions.** Mean rNMP EF (*y*-axis) within (**A**) 1 kb downstream, (**B**) 2-3 kb upstream and (**C**) 4-5 kb downstream of TSS for each rNMP: rA (red), rC (blue), rG (yellow), and rU (green) plotted across increasing expression levels grouped into deciles (*x*-axis, left to right) for each cell type. Each group contains an equal number of TSSs. Linear regression lines for each rNMP are shown, with Pearson's *R*, *p*-value, and linear regression slope summarized in **table S9** for each cell type.

**fig. S9**. **Reduced rC embedment on the non-template strand in RNH2A-KO cells within 0–1 kb downstream of TSS.** Mean rNMP EF (*y*-axis) within (**A**) 1 kb downstream, (**B**) 2-3 kb upstream and (**C**) 4-5 kb downstream of TSS for separate strands non-template (top) and template (bottom), for each rNMP: rA (red), rC (blue), rG (yellow), and rU (green) plotted across increasing expression levels grouped into deciles (*x*-axis, left to right) for each cell type. Each group contains an equal number of TSSs. Linear regression lines for each rNMP are shown, with Pearson's *R*, *p*-value, and linear regression slope summarized in **table S9** for each cell type.

**fig. S10**. **Raw rNMP composition, rNTP/dNTP ratios and di-nucleotide RN frequencies in all cell types.** (**A**) Bar graph showing mean and standard error of percentages of rNMPs rA (red), rC (blue), rG (yellow), and rU (green), without normalization to the background genomic frequency of dNMPs for libraries from the indicated cell type. (**B**) Bar graph showing rNTP pool to dNTP pool ratios in whole cells of $CD4^{+}T$, hESC-H9, HEK293T WT and RNH2A-KO from (*3*). (**C**) Heatmap displaying the normalized frequency of RN dinucleotides (each rNMP followed by dA, dC, dG, or dT) across all ribose-seq libraries (columns). Libraries of different cell types are separated by thick vertical blue lines; the two RNH2A-KO mutants are separated by a dashed vertical blue line. Genomic frequencies for each dinucleotide are shown on the left. The legend (top right) indicates frequency representation: white for 0.25; light red to red for 0.25 to 0.5–1; dark blue to light blue for 0-0.25. One-sided Mann-Whitney *U*-test *p*-values are presented in **table S11**.

**fig. S11**. **Specific trinucleotide patterns reveal the preference of nucleotides upstream of rNMP sites.** Heatmaps displaying the normalized frequency NNR, RNR, RNN trinucleotides of each rNMPs (rA, rC, rG, or rU) across RNH2A-KO libraries (columns). Libraries of the two RNH2A-KO mutants are separated by a dashed vertical blue line. Genomic frequencies for each trinucleotide are shown on the left. The legend (top right) indicates frequency representation: white for 0.0625; light red to red for greater than 0.0625; dark blue to light blue for 0-0.0625. One-sided Mann-Whitney *U*-test *p*-values are presented in **table S12**.

**fig. S12**. **rA increases and rC decreases in CpG islands and in the template strand downstream of the TSS of highly expressed genes in RNH2A-KO cells.** Heatmaps displaying the normalized frequency for rNMPs (rA, rC, rG, and rU) across RNH2A-KO libraries (columns). Libraries of the RNH2A-KO mutants are separated by a dashed vertical blue line. Genomic frequencies for each mononucleotide are shown on the left. The legend (top right) indicates frequency representation: white for 0.25; light red to red for 0.25 to 0.5–1; dark blue to light blue for 0-0.25. The mononucleotide heatmaps highlight preferences in: (**A**) CpG islands and inter-CpG regions, (**B**) 0 - 500 bp and 4-5 kb around TSS, (**C**) non-template and template strands, 500 bases downstream of TSSs of highly expressed genes (>75th percentile), and (**D**) non-template and template strands, 500 bases downstream of TSSs of low expressed genes (<25th percentile). Comparisons between RNH2A-KO libraries (two-tailed Mann–Whitney *U*-test) are marked with asterisks (*). Significance levels: * ($0.05 \geq p > 0.01$), ** ($0.01 \geq p > 0.001$). Two-sided unpaired and paired Mann-Whitney *U*-test *p*-values are presented in **table S13**.

**fig. S13. rNMPs preferentially accumulate in CpG islands, PLS regions, near promoters, and telomeres.** Schematic representations of rNMP distribution across annotated genomic features in (**A**) $CD4^{+}T$, (**B**) hESC-H9 and (**C**) RNH2A-KO libraries. Top panel: CpG island (dark orange), shore and shelf (lighter shades of orange), and inter-CpG (yellow); each region represents 100-kb genomic segments. Middle panel: cCREs, including PLS (dark orange), dELS and pELS (light orange) with respect to the TSS, regions with CTCF-only signature (light orange), DNase-H3K4me3 signature (orange), inter-cCRE (light blue), and telomeres (purple). Bottom panel: gene-associated regions, including enhancer (1–5 kb upstream of TSS, light orange), promoter (0–

1 kb upstream of TSS, orange), 5′ and 3′ UTRs (green), exons (blue) and introns (light blue). Exons are categorized as first, last, and internal exons, and introns as proximal and distal introns to show proximity to TSSs and TTSs. Pin size and color intensity (pink/darkness) indicate rNMP counts per 100 kb per million rNMPs in the nuclear genome of each library, as defined by the formula shown in the figure. See data in **table S16**.

**fig. S14 Model for the epigenetic role of rNMPs near TSSs in RNH2A-KO cells.** In the absence of RNase H2, embedded rNMPs, particularly rCs and rUs, are targeted by Top1 along gene bodies, from the transcription start site (TSS) to the transcription termination site (TTS), with enrichment downstream of the TSS. This activity relieves torsional stress generated during replication and transcription, particularly on the leading strand, thereby facilitating transcriptional restart.

### fig. S1

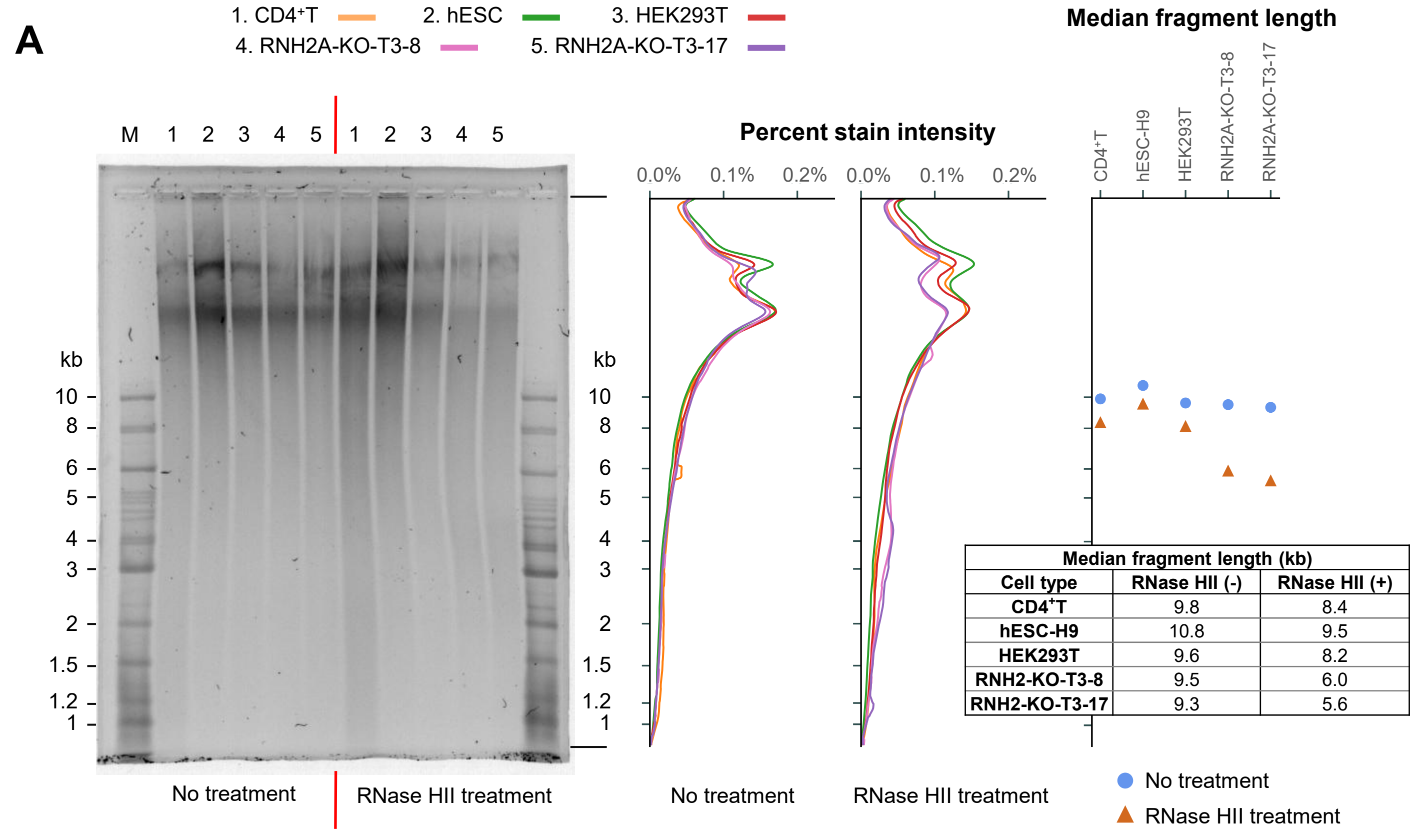


| Median fragment length (kb) | | |
|---|---|---|
| **Cell type** | **RNase HII (-)** | **RNase HII (+)** |
| **CD4⁺T** | 9.8 | 8.4 |
| **hESC-H9** | 10.8 | 9.5 |
| **HEK293T** | 9.6 | 8.2 |
| **RNH2-KO-T3-8** | 9.5 | 6.0 |
| **RNH2-KO-T3-17** | 9.3 | 5.6 |

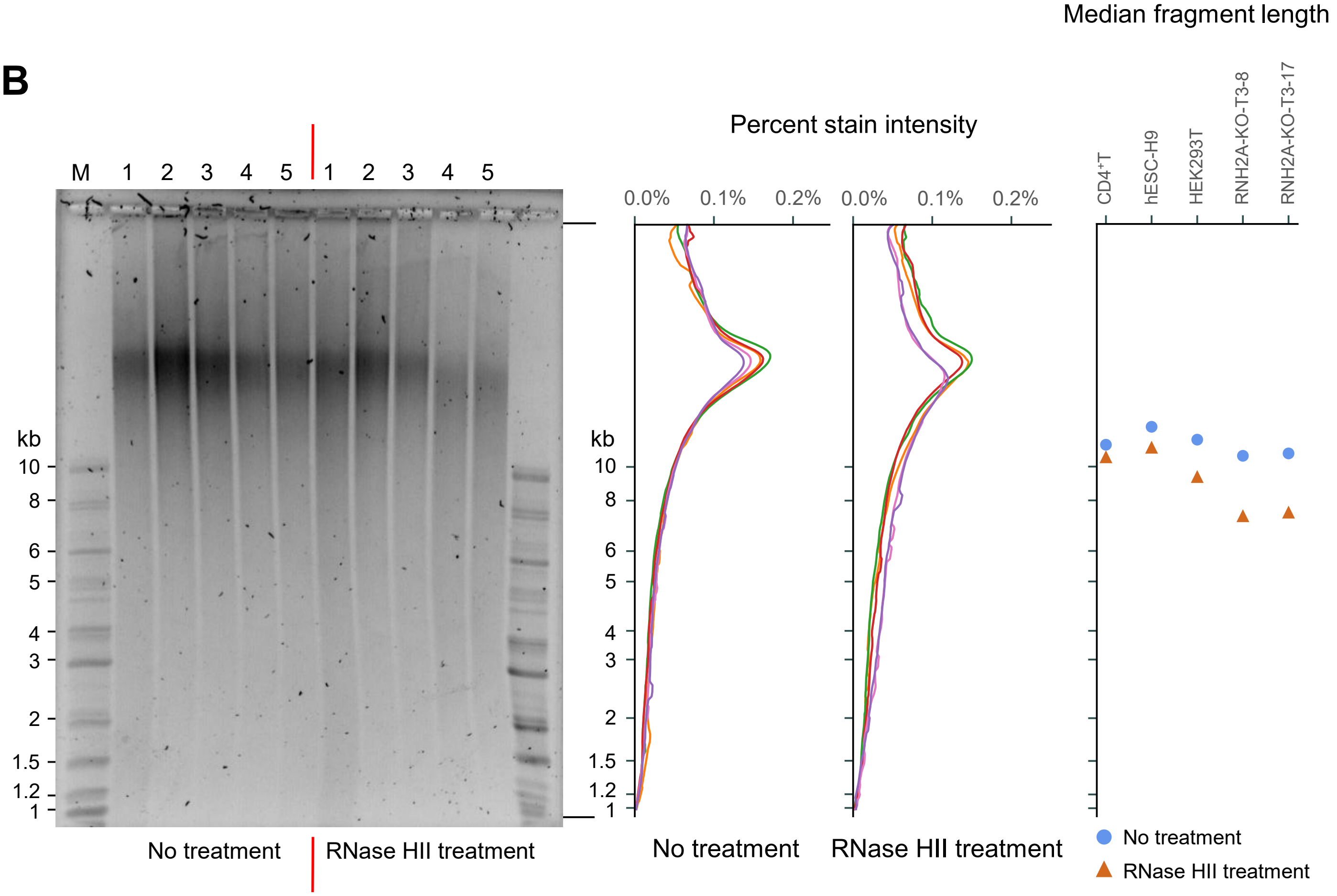

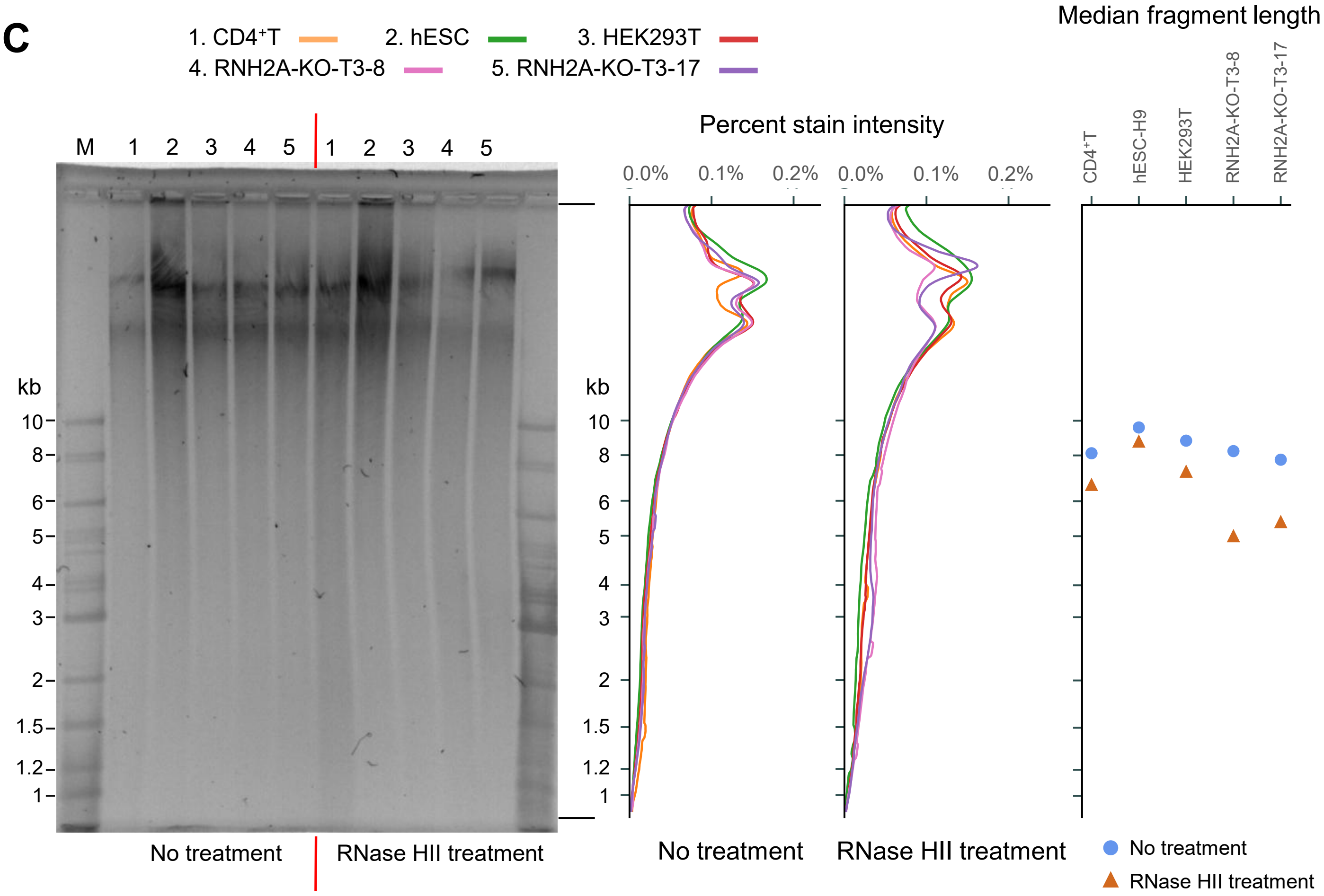
C
1. CD4⁺T
2. hESC
3. HEK293T
4. RNH2A-KO-T3-8
5. RNH2A-KO-T3-17
M 1 2 3 4 5 1 2 3 4 5
kb
10
8
6
5
4
3
2
1.5
1.2
1
No treatment
RNase HII treatment
Percent stain intensity
0.0% 0.1% 0.2%
0.0% 0.1% 0.2%
No treatment
RNase HII treatment
Median fragment length
CD4⁺T
hESC-H9
HEK293T
RNH2A-KO-T3-8
RNH2A-KO-T3-17
No treatment
RNase HII treatment

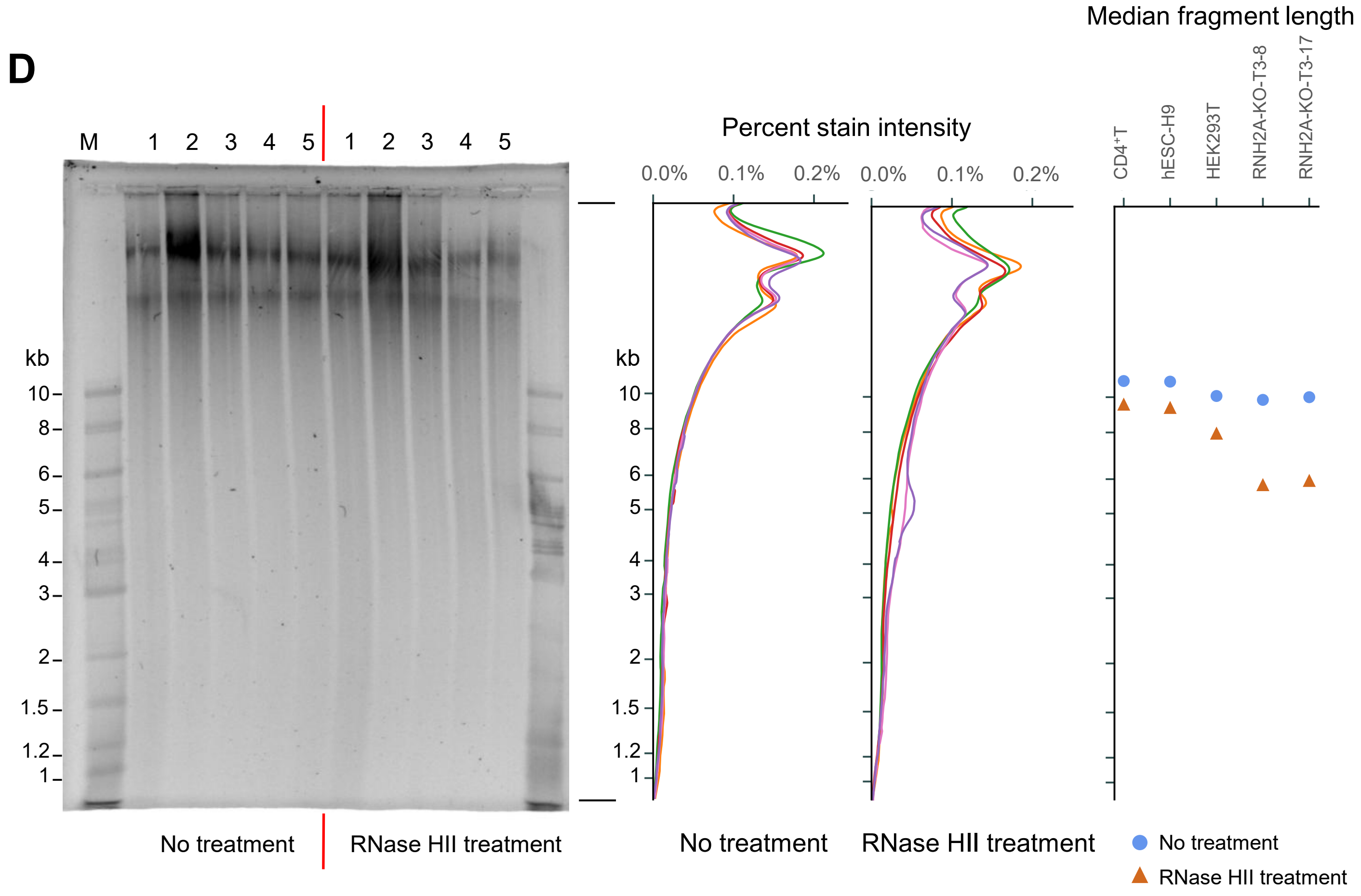
D
M 1 2 3 4 5 1 2 3 4 5
kb
10
8
6
5
4
3
2
1.5
1.2
1
No treatment
RNase HII treatment
Percent stain intensity
0.0% 0.1% 0.2%
0.0% 0.1% 0.2%
No treatment
RNase HII treatment
Median fragment length
CD4⁺T
hESC-H9
HEK293T
RNH2A-KO-T3-8
RNH2A-KO-T3-17
No treatment
RNase HII treatment

**fig. S2**

**dNMP frequency +/- 50 bp around rNMPs**

Relative frequency: 2 — 1.5 — 1 — 0.5 — 0

| CD4⁺T | | | | | hESC-H9 | | | | | | HEK293T | | | | | RNH2A-KO-T3-8 | | | | | RNH2A-KO-T3-17 | | | | |
|---|---|---|---|---|---|---|---|---|---|---|---|---|---|---|---|---|---|---|---|---|---|---|---|---|---|
| FS185-F | FS187-FRE1 | FS188-FRE1 | FS189-FRE1 | FS186-RE1 | FS197-F | FS198-F | FS201-FRE1 | FS199-RE1 | FS331-RE2 | FS303-RE3 | FS326-F | FS391-F | FS203-FRE1 | FS333-RE2 | FS305-RE3 | FS327-F | FS392-F | FS300-RE1 | FS309-RE2 | FS306-RE3 | FS329-F | FS393-F | FS301-RE1 | FS307-RE2 | FS310-RE3 |
| 0.91 | 1.05 | 0.99 | 1.00 | 0.96 | 0.96 | 1.00 | 1.00 | 1.03 | 1.02 | 0.98 | 0.99 | 0.96 | 0.95 | 0.88 | 0.90 | 0.94 | 0.95 | 0.90 | 0.97 | 0.94 | 0.92 | 0.98 | 0.90 | 1.00 | 1.02 |
| 1.15 | 0.94 | 1.06 | 1.04 | 1.07 | 1.12 | 1.07 | 1.05 | 0.96 | 0.94 | 1.06 | 1.06 | 1.08 | 1.07 | 1.10 | 1.19 | 1.15 | 1.14 | 1.18 | 1.13 | 1.17 | 1.18 | 1.09 | 1.21 | 1.08 | 1.05 |
| 1.17 | 1.05 | 1.12 | 1.08 | 1.04 | 1.10 | 1.09 | 1.08 | 1.11 | 1.13 | 1.12 | 1.05 | 1.03 | 1.09 | 1.12 | 1.11 | 1.05 | 1.16 | 1.15 | 1.14 | 1.14 | 1.13 | 1.16 | 1.14 | 1.10 | 1.11 |
| 0.86 | 0.96 | 0.88 | 0.91 | 0.96 | 0.88 | 0.89 | 0.91 | 0.92 | 0.93 | 0.90 | 0.93 | 0.96 | 0.94 | 0.97 | 0.90 | 0.92 | 0.84 | 0.86 | 0.84 | 0.84 | 0.86 | 0.85 | 0.86 | 0.88 | 0.87 |

| CD4⁺T | | | | | hESC-H9 | | | | | | HEK293T | | | | | RNH2A-KO-T3-8 | | | | | RNH2A-KO-T3-17 | | | | |
|---|---|---|---|---|---|---|---|---|---|---|---|---|---|---|---|---|---|---|---|---|---|---|---|---|---|
| FS185-F | FS187-FRE1 | FS188-FRE1 | FS189-FRE1 | FS186-RE1 | FS197-F | FS198-F | FS201-FRE1 | FS199-RE1 | FS331-RE2 | FS303-RE3 | FS326-F | FS391-F | FS203-FRE1 | FS333-RE2 | FS305-RE3 | FS327-F | FS392-F | FS300-RE1 | FS309-RE2 | FS306-RE3 | FS329-F | FS393-F | FS301-RE1 | FS307-RE2 | FS310-RE3 |
| 0.84 | 1.08 | 0.98 | 1.00 | 0.92 | 0.93 | 1.02 | 1.01 | 1.04 | 1.05 | 0.93 | 1.00 | 0.94 | 0.89 | 0.81 | 0.80 | 0.87 | 0.88 | 0.80 | 0.92 | 0.86 | 0.83 | 0.94 | 0.79 | 0.98 | 1.01 |
| 1.00 | 0.99 | 1.03 | 1.04 | 1.01 | 1.07 | 1.03 | 1.01 | 0.95 | 0.92 | 1.02 | 1.06 | 1.04 | 0.99 | 0.94 | 1.06 | 1.08 | 1.08 | 1.04 | 1.12 | 1.09 | 1.07 | 1.06 | 1.07 | 1.10 | 1.10 |
| 1.11 | 1.06 | 1.11 | 1.09 | 1.02 | 1.08 | 1.09 | 1.10 | 1.16 | 1.16 | 1.14 | 1.05 | 1.02 | 1.06 | 0.98 | 1.05 | 1.03 | 1.14 | 1.09 | 1.14 | 1.12 | 1.09 | 1.15 | 1.08 | 1.10 | 1.14 |
| 0.76 | 1.01 | 0.85 | 0.90 | 0.93 | 0.82 | 0.85 | 0.87 | 0.93 | 0.93 | 0.84 | 0.89 | 0.89 | 0.88 | 0.83 | 0.77 | 0.84 | 0.78 | 0.76 | 0.78 | 0.76 | 0.77 | 0.82 | 0.75 | 0.86 | 0.87 |
| 1.05 | 1.00 | 1.04 | 1.03 | 1.03 | 1.07 | 1.04 | 1.05 | 1.05 | 0.93 | 1.03 | 1.04 | 1.03 | 1.01 | 0.94 | 1.05 | 1.08 | 1.09 | 1.06 | 1.11 | 1.10 | 1.09 | 1.07 | 1.09 | 1.08 | 1.08 |
| 1.33 | 0.89 | 1.14 | 1.10 | 1.16 | 1.25 | 1.17 | 1.11 | 0.90 | 0.95 | 1.10 | 1.12 | 1.16 | 1.16 | 1.24 | 1.35 | 1.30 | 1.27 | 1.40 | 1.25 | 1.33 | 1.36 | 1.20 | 1.45 | 1.14 | 1.09 |
| 1.57 | 0.99 | 1.35 | 1.20 | 1.18 | 1.40 | 1.29 | 1.22 | 0.99 | 1.10 | 1.28 | 1.21 | 1.20 | 1.29 | 1.77 | 1.64 | 1.38 | 1.66 | 1.71 | 1.61 | 1.65 | 1.62 | 1.54 | 1.71 | 1.38 | 1.33 |
| 1.06 | 0.90 | 0.99 | 0.98 | 1.04 | 1.02 | 0.99 | 0.99 | 0.92 | 0.91 | 1.02 | 1.00 | 1.05 | 1.03 | 1.05 | 1.13 | 1.08 | 1.00 | 1.07 | 1.00 | 1.04 | 1.07 | 0.97 | 1.09 | 0.97 | 0.95 |
| 1.09 | 1.08 | 1.11 | 1.10 | 1.01 | 1.06 | 1.09 | 1.09 | 1.14 | 1.20 | 1.14 | 1.05 | 1.00 | 1.06 | 0.97 | 1.03 | 1.01 | 1.14 | 1.07 | 1.15 | 1.10 | 1.06 | 1.18 | 1.05 | 1.12 | 1.16 |
| 1.30 | 1.00 | 1.18 | 1.08 | 1.09 | 1.21 | 1.12 | 1.12 | 1.10 | 1.01 | 1.15 | 1.10 | 1.09 | 1.13 | 1.21 | 1.29 | 1.18 | 1.28 | 1.31 | 1.23 | 1.30 | 1.31 | 1.22 | 1.31 | 1.13 | 1.13 |
| 1.37 | 1.06 | 1.22 | 1.15 | 1.06 | 1.22 | 1.19 | 1.16 | 1.17 | 1.27 | 1.20 | 1.12 | 1.07 | 1.18 | 1.28 | 1.20 | 1.12 | 1.33 | 1.33 | 1.27 | 1.29 | 1.28 | 1.32 | 1.30 | 1.20 | 1.20 |
| 0.98 | 1.04 | 0.97 | 0.97 | 0.99 | 0.95 | 0.97 | 0.97 | 1.03 | 1.03 | 1.01 | 0.95 | 0.97 | 1.03 | 1.07 | 0.96 | 0.93 | 0.92 | 0.94 | 0.94 | 0.93 | 0.93 | 0.92 | 0.93 | 0.95 | 0.94 |
| 0.73 | 1.04 | 0.85 | 0.92 | 0.91 | 0.81 | 0.85 | 0.88 | 0.94 | 0.93 | 0.84 | 0.90 | 0.89 | 0.89 | 0.83 | 0.74 | 0.83 | 0.74 | 0.72 | 0.73 | 0.73 | 0.74 | 0.76 | 0.71 | 0.84 | 0.85 |
| 1.01 | 0.88 | 0.94 | 0.96 | 1.02 | 0.99 | 0.98 | 0.96 | 0.88 | 0.90 | 0.99 | 0.99 | 1.04 | 0.98 | 1.03 | 1.11 | 1.08 | 0.97 | 1.04 | 0.98 | 1.00 | 1.04 | 0.95 | 1.06 | 0.97 | 0.94 |
| 1.01 | 1.02 | 1.01 | 0.99 | 1.02 | 1.00 | 0.98 | 0.99 | 1.04 | 1.01 | 1.03 | 0.98 | 0.99 | 1.04 | 1.06 | 1.03 | 0.98 | 0.99 | 1.01 | 0.99 | 1.00 | 1.01 | 1.00 | 1.00 | 0.98 | 0.98 |
| 0.75 | 0.93 | 0.79 | 0.84 | 0.93 | 0.79 | 0.81 | 0.84 | 0.85 | 0.89 | 0.80 | 0.88 | 0.94 | 0.87 | 0.96 | 0.79 | 0.84 | 0.72 | 0.75 | 0.71 | 0.73 | 0.74 | 0.74 | 0.74 | 0.78 | 0.77 |

fig. S3

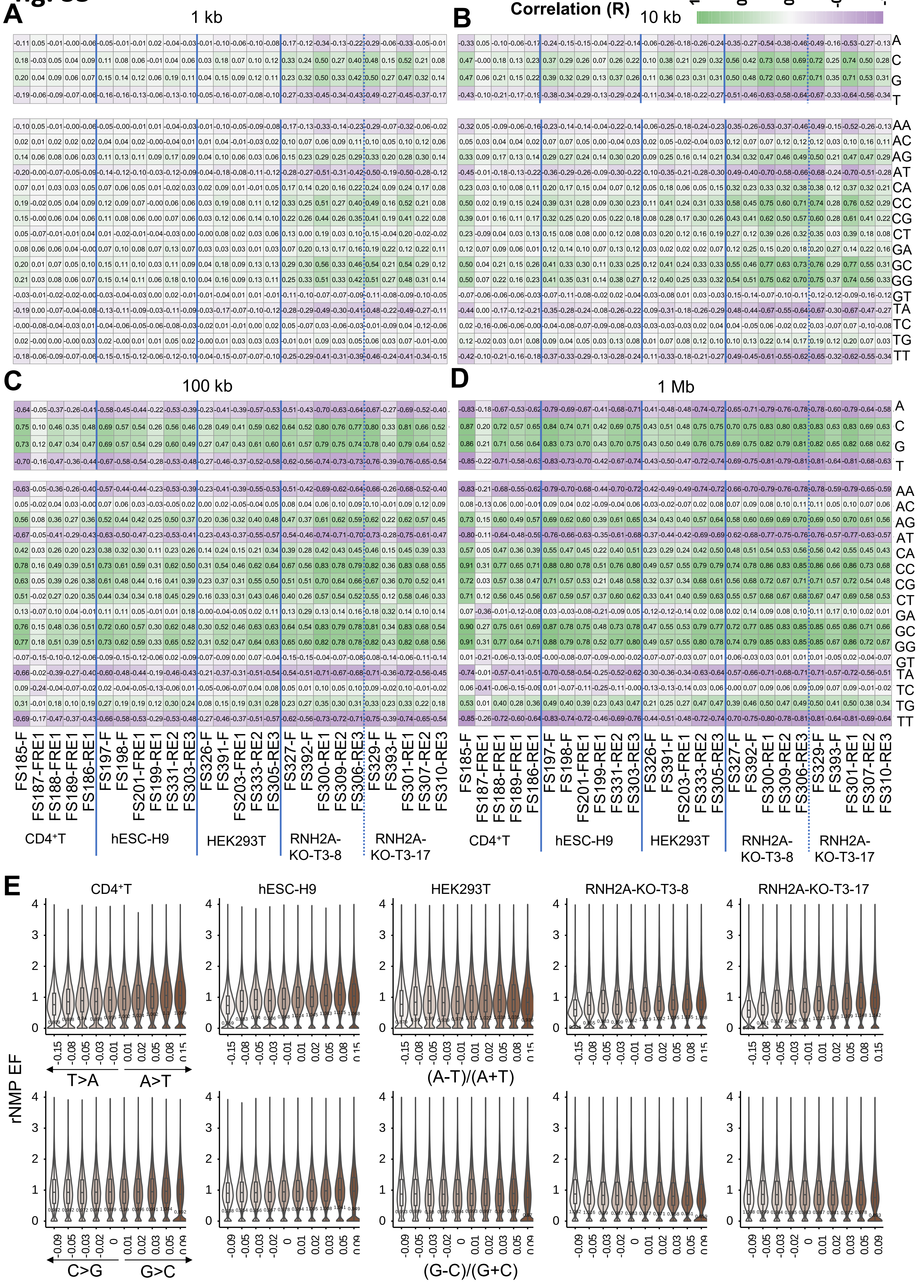

A
1 kb
B
10 kb
Pearson's Correlation (R)
1
0.5
0
-0.5
-1
C
100 kb
D
1 Mb
A
C
G
T
AA
AC
AG
AT
CA
CC
CG
CT
GA
GC
GG
GT
TA
TC
TG
TT
FS185-F
FS187-FRE1
FS188-FRE1
FS189-FRE1
FS186-RE1
FS197-F
FS198-F
FS201-FRE1
FS199-RE1
FS331-RE2
FS303-RE3
FS326-F
FS391-F
FS203-FRE1
FS333-RE2
FS305-RE3
FS327-F
FS392-F
FS300-RE1
FS309-RE2
FS306-RE3
FS329-F
FS393-F
FS301-RE1
FS307-RE2
FS310-RE3
CD4+T
hESC-H9
HEK293T
RNH2A-KO-T3-8
RNH2A-KO-T3-17
E
rNMP EF
T>A
A>T
(A-T)/(A+T)
C>G
G>C
(G-C)/(G+C)

### fig. S4

A

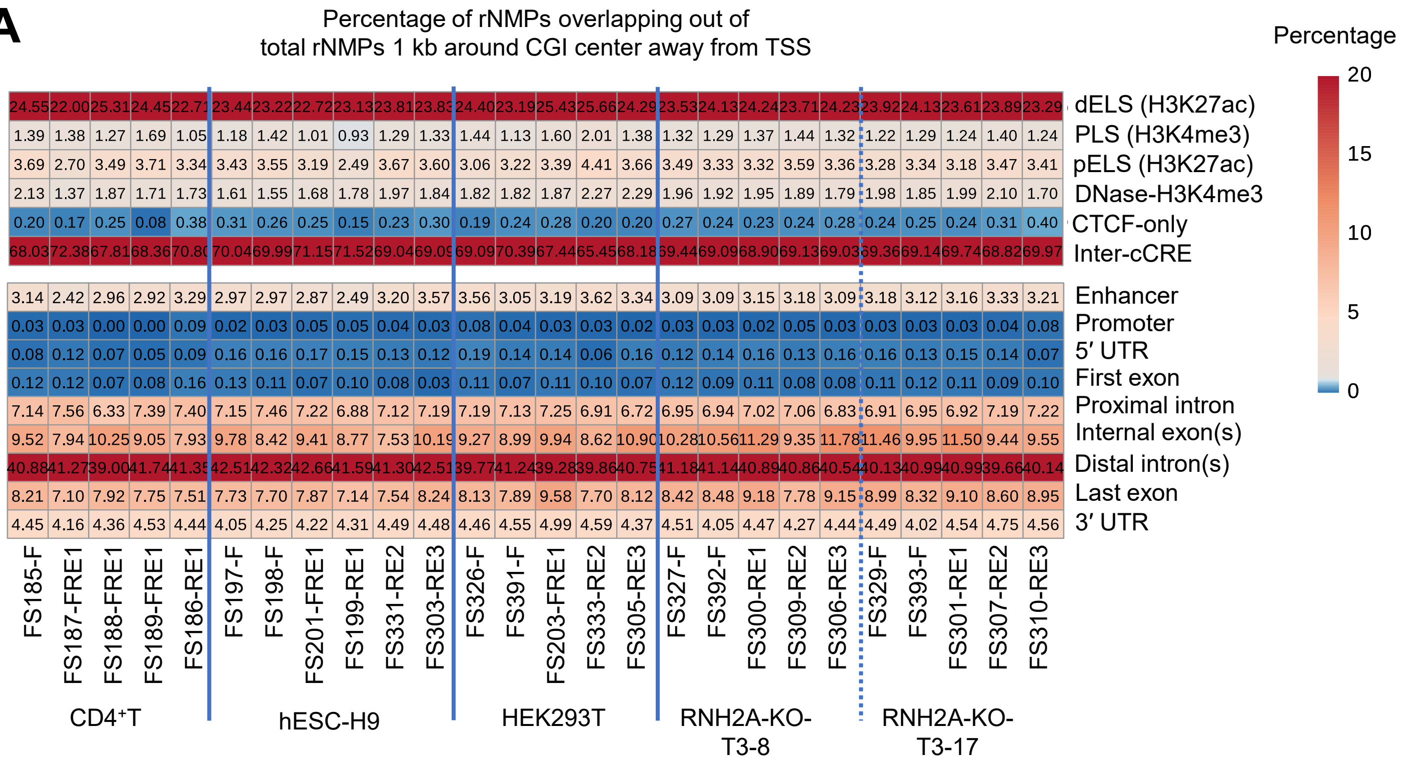

Percentage of rNMPs overlapping out of
total rNMPs 1 kb around CGI center away from TSS
Percentage
20
15
10
5
0
dELS (H3K27ac)
PLS (H3K4me3)
pELS (H3K27ac)
DNase-H3K4me3
CTCF-only
Inter-cCRE
Enhancer
Promoter
5′ UTR
First exon
Proximal intron
Internal exon(s)
Distal intron(s)
Last exon
3′ UTR
24.55 22.00 25.31 24.45 22.71 23.44 23.22 22.72 23.13 23.81 23.83 24.40 23.19 25.43 25.66 24.29 23.53 24.13 24.24 23.71 24.23 23.92 24.13 23.61 23.89 23.29
1.39 1.38 1.27 1.69 1.05 1.18 1.42 1.01 0.93 1.29 1.33 1.44 1.13 1.60 2.01 1.38 1.32 1.29 1.37 1.44 1.32 1.22 1.29 1.24 1.40 1.24
3.69 2.70 3.49 3.71 3.34 3.43 3.55 3.19 2.49 3.67 3.60 3.06 3.22 3.39 4.41 3.66 3.49 3.33 3.32 3.59 3.36 3.28 3.34 3.18 3.47 3.41
2.13 1.37 1.87 1.71 1.73 1.61 1.55 1.68 1.78 1.97 1.84 1.82 1.82 1.87 2.27 2.29 1.96 1.92 1.95 1.89 1.79 1.98 1.85 1.99 2.10 1.70
0.20 0.17 0.25 0.08 0.38 0.31 0.26 0.25 0.15 0.23 0.30 0.19 0.24 0.28 0.20 0.20 0.27 0.24 0.23 0.24 0.28 0.24 0.25 0.24 0.31 0.40
68.03 72.38 67.81 68.36 70.80 70.04 69.99 71.15 71.52 69.04 69.09 69.09 70.39 67.44 65.45 68.18 69.44 69.09 68.90 69.13 69.03 69.36 69.14 69.74 68.82 69.97
3.14 2.42 2.96 2.92 3.29 2.97 2.97 2.87 2.49 3.20 3.57 3.56 3.05 3.19 3.62 3.34 3.09 3.09 3.15 3.18 3.09 3.18 3.12 3.16 3.33 3.21
0.03 0.03 0.00 0.00 0.09 0.02 0.03 0.05 0.05 0.04 0.03 0.08 0.04 0.03 0.03 0.02 0.03 0.03 0.02 0.05 0.03 0.03 0.03 0.03 0.04 0.08
0.08 0.12 0.07 0.05 0.09 0.16 0.16 0.17 0.15 0.13 0.12 0.19 0.14 0.14 0.06 0.16 0.12 0.14 0.16 0.13 0.16 0.16 0.13 0.15 0.14 0.07
0.12 0.12 0.07 0.08 0.16 0.13 0.11 0.07 0.10 0.08 0.03 0.11 0.07 0.11 0.10 0.07 0.12 0.09 0.11 0.08 0.08 0.11 0.12 0.11 0.09 0.10
7.14 7.56 6.33 7.39 7.40 7.15 7.46 7.22 6.88 7.12 7.19 7.19 7.13 7.25 6.91 6.72 6.95 6.94 7.02 7.06 6.83 6.91 6.95 6.92 7.19 7.22
9.52 7.94 10.25 9.05 7.93 9.78 8.42 9.41 8.77 7.53 10.19 9.27 8.99 9.94 8.62 10.90 10.28 10.56 11.29 9.35 11.78 11.46 9.95 11.50 9.44 9.55
40.88 41.27 39.00 41.74 41.35 42.51 42.32 42.66 41.59 41.30 42.51 39.77 41.24 39.28 39.86 40.75 41.18 41.14 40.89 40.86 40.54 40.13 40.99 40.99 39.66 40.14
8.21 7.10 7.92 7.75 7.51 7.73 7.70 7.87 7.14 7.54 8.24 8.13 7.89 9.58 7.70 8.12 8.42 8.48 9.18 7.78 9.15 8.99 8.32 9.10 8.60 8.95
4.45 4.16 4.36 4.53 4.44 4.05 4.25 4.22 4.31 4.49 4.48 4.46 4.55 4.99 4.59 4.37 4.51 4.05 4.47 4.27 4.44 4.49 4.02 4.54 4.75 4.56
FS185-F
FS187-FRE1
FS188-FRE1
FS189-FRE1
FS186-RE1
FS197-F
FS198-F
FS201-FRE1
FS199-RE1
FS331-RE2
FS303-RE3
FS326-F
FS391-F
FS203-FRE1
FS333-RE2
FS305-RE3
FS327-F
FS392-F
FS300-RE1
FS309-RE2
FS306-RE3
FS329-F
FS393-F
FS301-RE1
FS307-RE2
FS310-RE3
CD4⁺T
hESC-H9
HEK293T
RNH2A-KO-T3-8
RNH2A-KO-T3-17

fig. S5

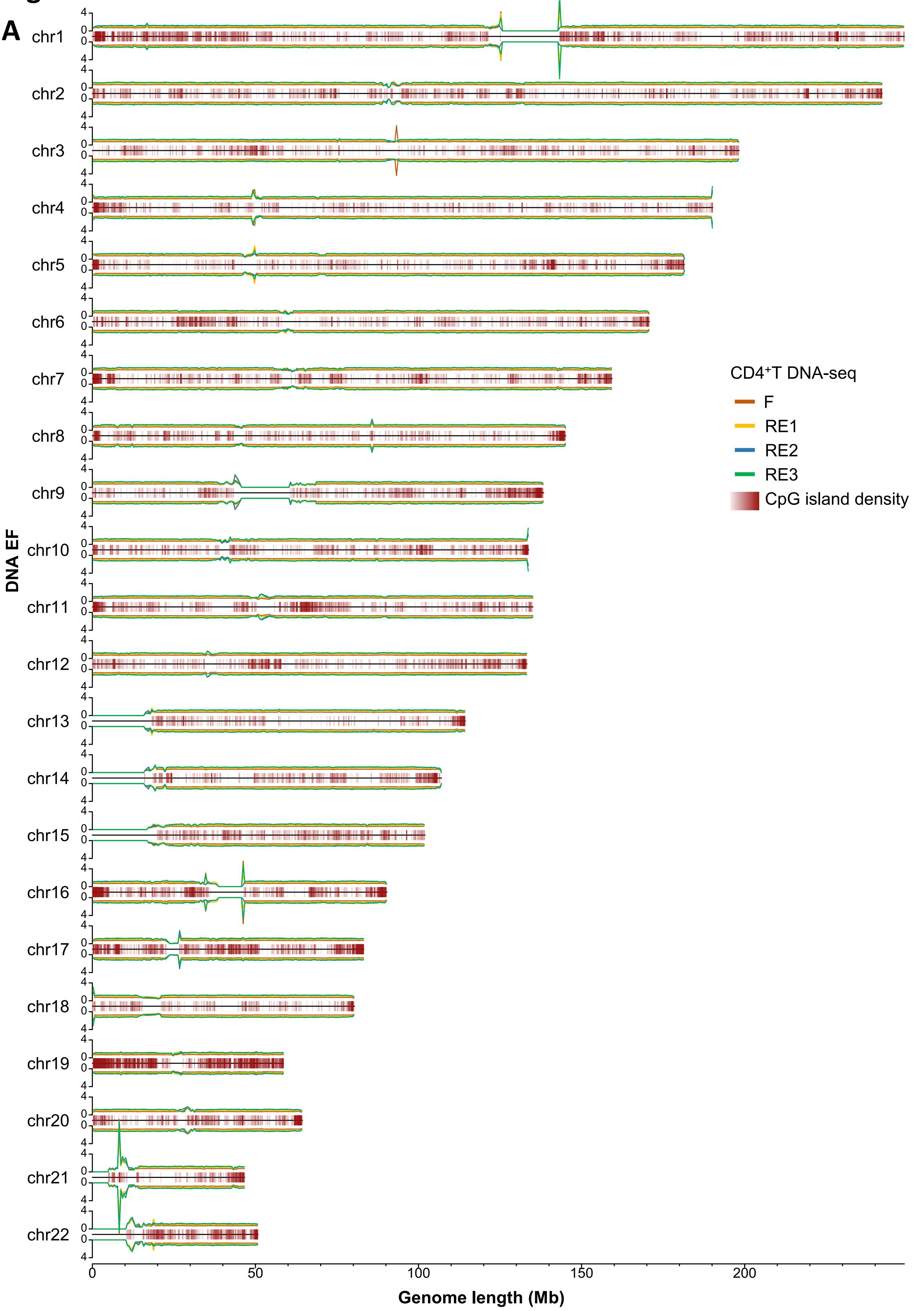
A
chr1
chr2
chr3
chr4
chr5
chr6
chr7
chr8
chr9
chr10
chr11
chr12
chr13
chr14
chr15
chr16
chr17
chr18
chr19
chr20
chr21
chr22
DNA EF
Genome length (Mb)
0
50
100
150
200
CD4⁺T DNA-seq
F
RE1
RE2
RE3
CpG island density

fig. S5

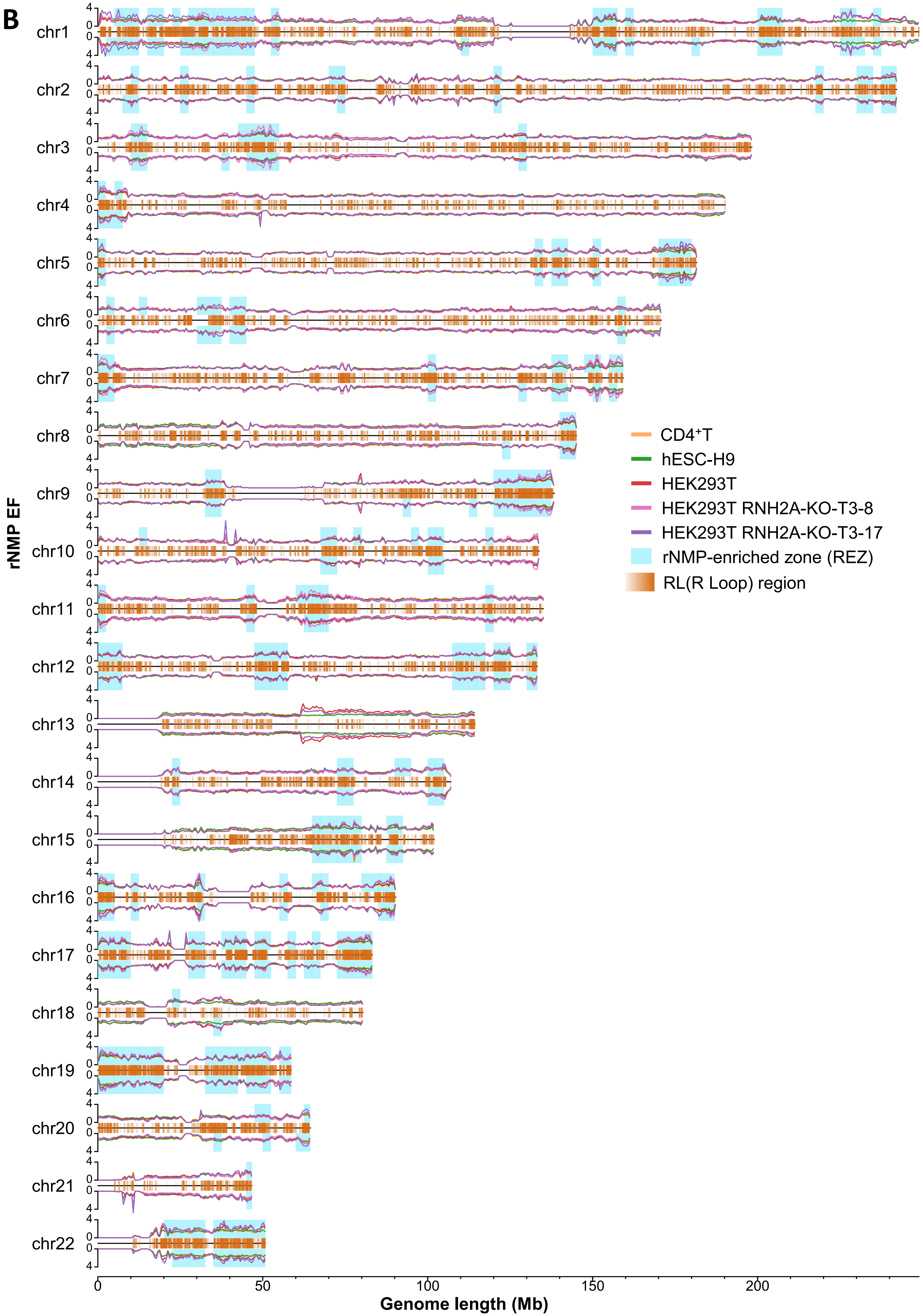
B
chr1
chr2
chr3
chr4
chr5
chr6
chr7
chr8
chr9
chr10
chr11
chr12
chr13
chr14
chr15
chr16
chr17
chr18
chr19
chr20
chr21
chr22
rNMP EF
CD4⁺T
hESC-H9
HEK293T
HEK293T RNH2A-KO-T3-8
HEK293T RNH2A-KO-T3-17
rNMP-enriched zone (REZ)
RL(R Loop) region
0
50
100
150
200
Genome length (Mb)

### fig. S5

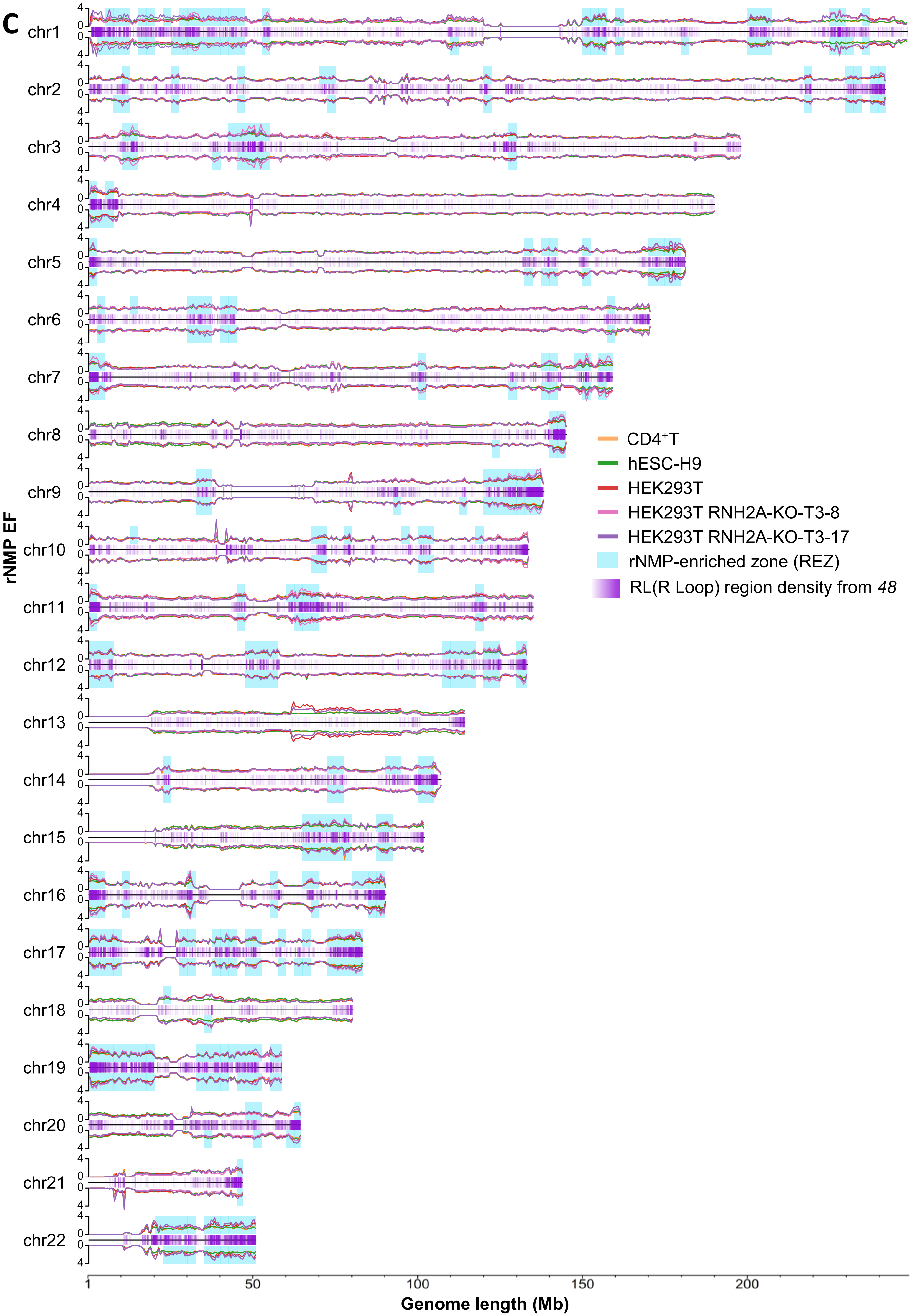
C
chr1
chr2
chr3
chr4
chr5
chr6
chr7
chr8
chr9
chr10
chr11
chr12
chr13
chr14
chr15
chr16
chr17
chr18
chr19
chr20
chr21
chr22
rNMP EF
CD4⁺T
hESC-H9
HEK293T
HEK293T RNH2A-KO-T3-8
HEK293T RNH2A-KO-T3-17
rNMP-enriched zone (REZ)
RL(R Loop) region density from 48
1
50
100
150
200
Genome length (Mb)

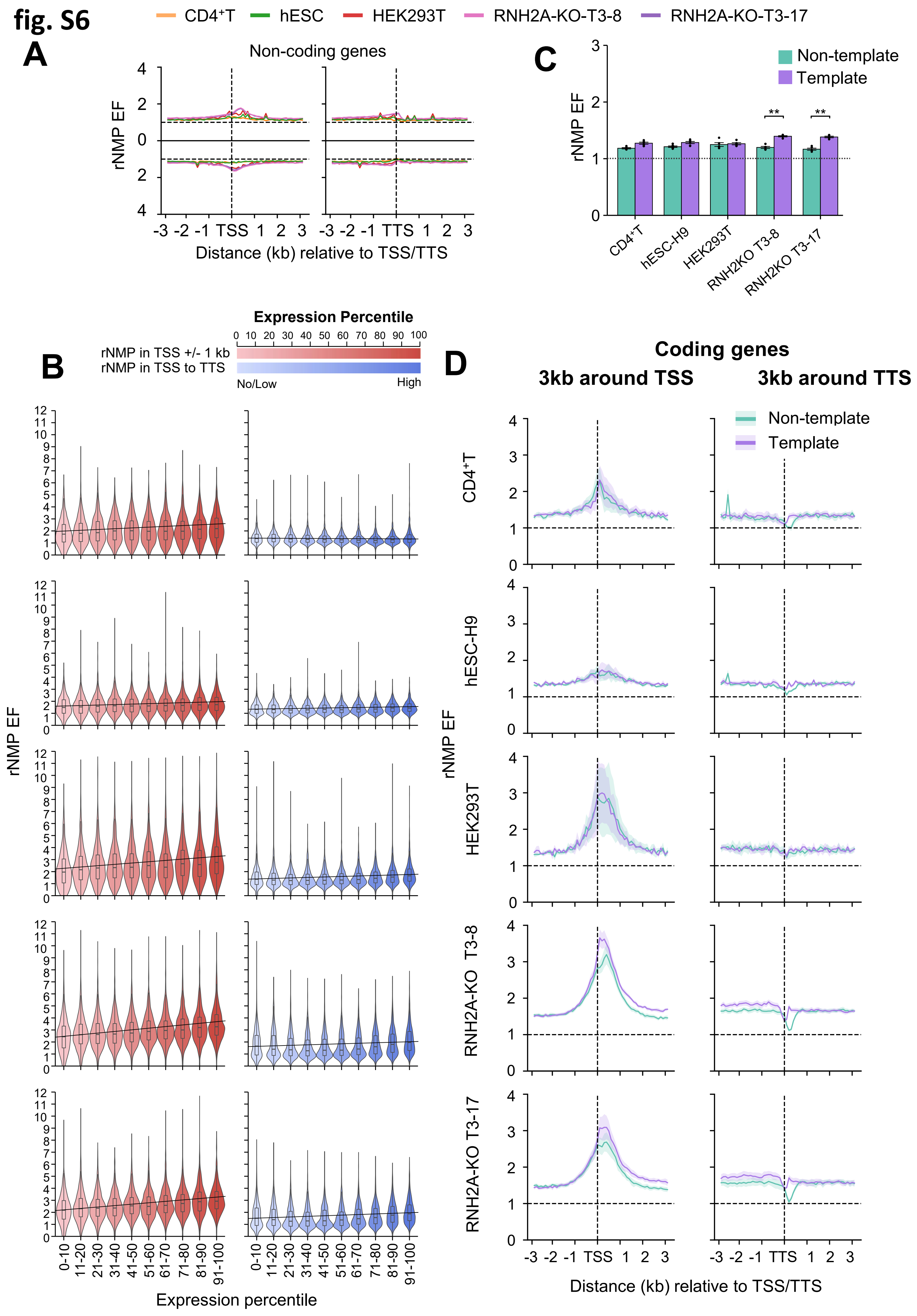
fig. S6
CD4⁺T
hESC
HEK293T
RNH2A-KO-T3-8
RNH2A-KO-T3-17
A
Non-coding genes
rNMP EF
Distance (kb) relative to TSS/TTS
C
Non-template
Template
rNMP EF
CD4⁺T
hESC-H9
HEK293T
RNH2KO T3-8
RNH2KO T3-17
Expression Percentile
rNMP in TSS +/- 1 kb
rNMP in TSS to TTS
No/Low
High
B
rNMP EF
Expression percentile
D
Coding genes
3kb around TSS
3kb around TTS
Non-template
Template
CD4⁺T
hESC-H9
HEK293T
RNH2A-KO T3-8
RNH2A-KO T3-17
rNMP EF
Distance (kb) relative to TSS/TTS

**fig. S7**

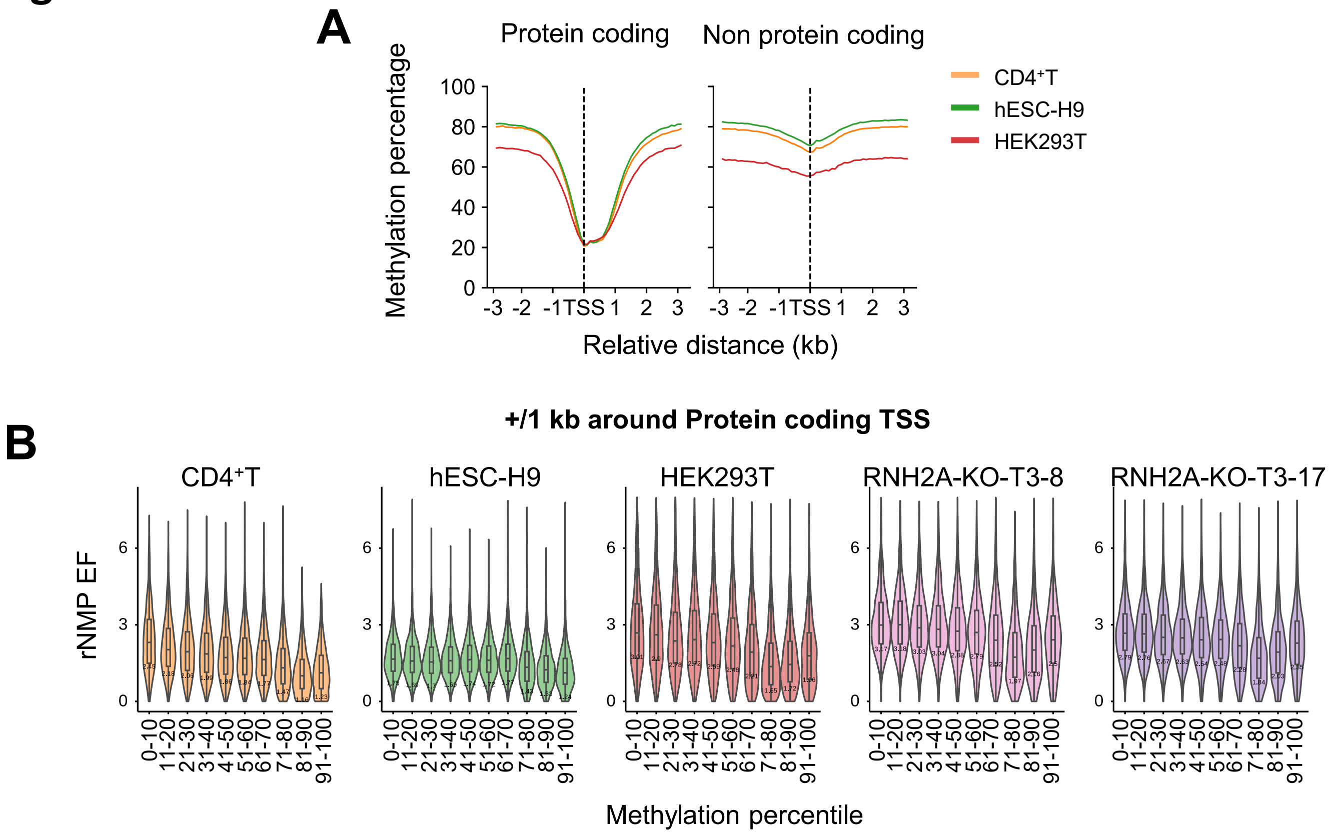
A
Protein coding
Non protein coding
Methylation percentage
100
80
60
40
20
0
-3 -2 -1 TSS 1 2 3
-3 -2 -1 TSS 1 2 3
Relative distance (kb)
CD4⁺T
hESC-H9
HEK293T
+/1 kb around Protein coding TSS
B
CD4⁺T
hESC-H9
HEK293T
RNH2A-KO-T3-8
RNH2A-KO-T3-17
rNMP EF
6
3
0
0-10
11-20
21-30
31-40
41-50
51-60
61-70
71-80
81-90
91-100
Methylation percentile

### fig. S8

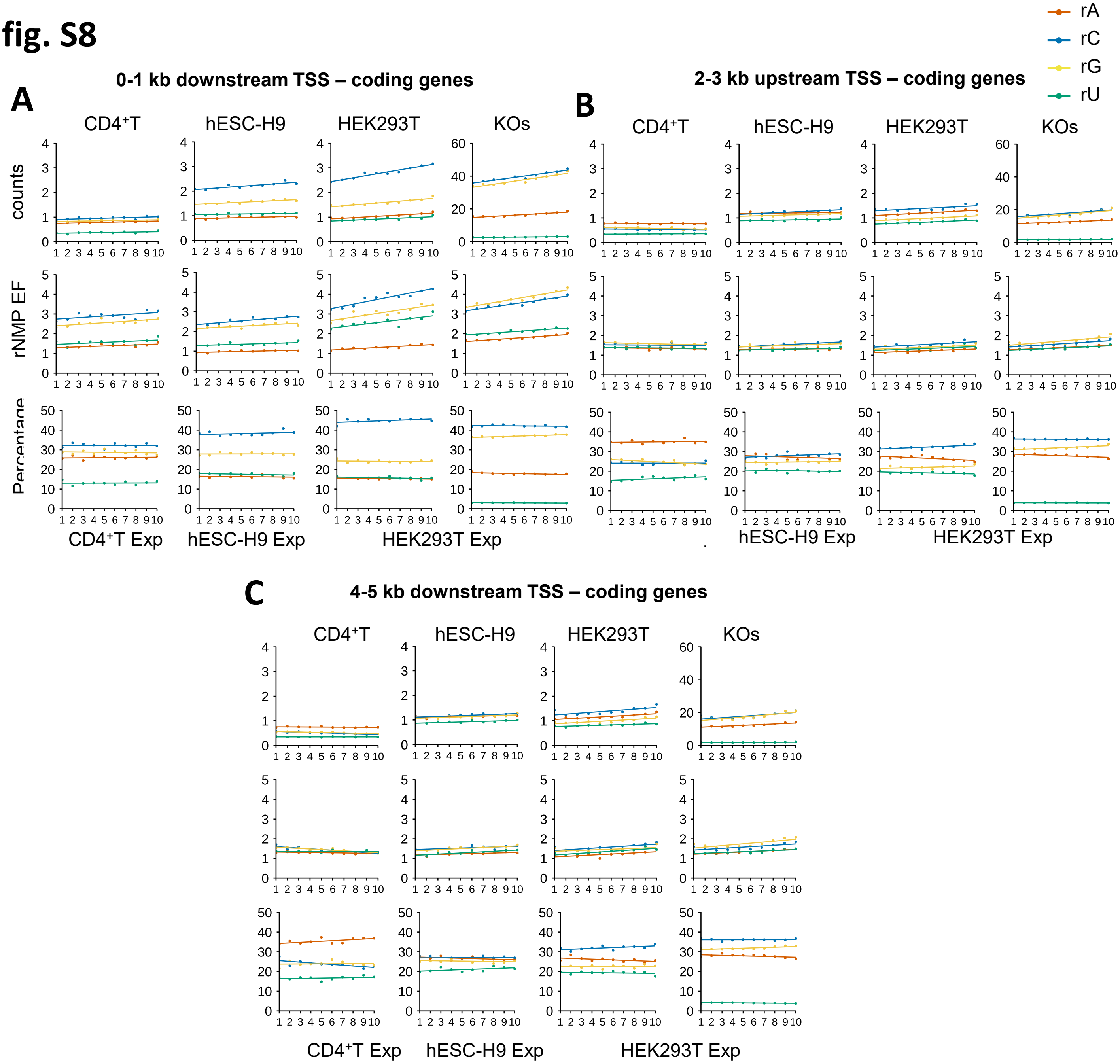
rA
rC
rG
rU
A
0-1 kb downstream TSS – coding genes
B
2-3 kb upstream TSS – coding genes
C
4-5 kb downstream TSS – coding genes
CD4⁺T
hESC-H9
HEK293T
KOs
counts
rNMP EF
Percentage
CD4⁺T Exp
hESC-H9 Exp
HEK293T Exp

fig. S9

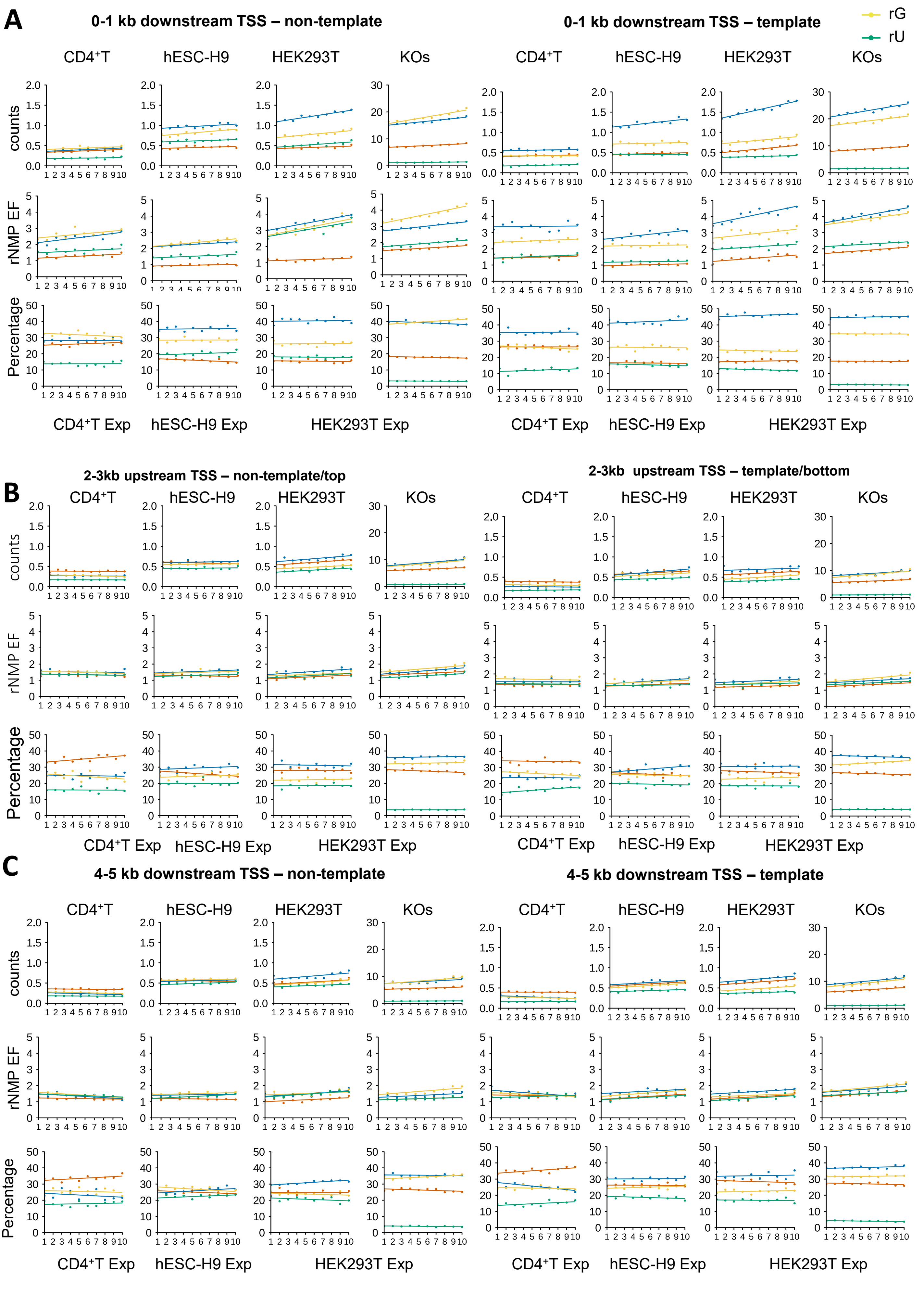

rA
rC
rG
rU
A
0-1 kb downstream TSS – non-template
0-1 kb downstream TSS – template
CD4+T
hESC-H9
HEK293T
KOs
counts
rNMP EF
Percentage
CD4+T Exp
hESC-H9 Exp
HEK293T Exp
B
2-3kb upstream TSS – non-template/top
2-3kb upstream TSS – template/bottom
C
4-5 kb downstream TSS – non-template
4-5 kb downstream TSS – template

fig. S10

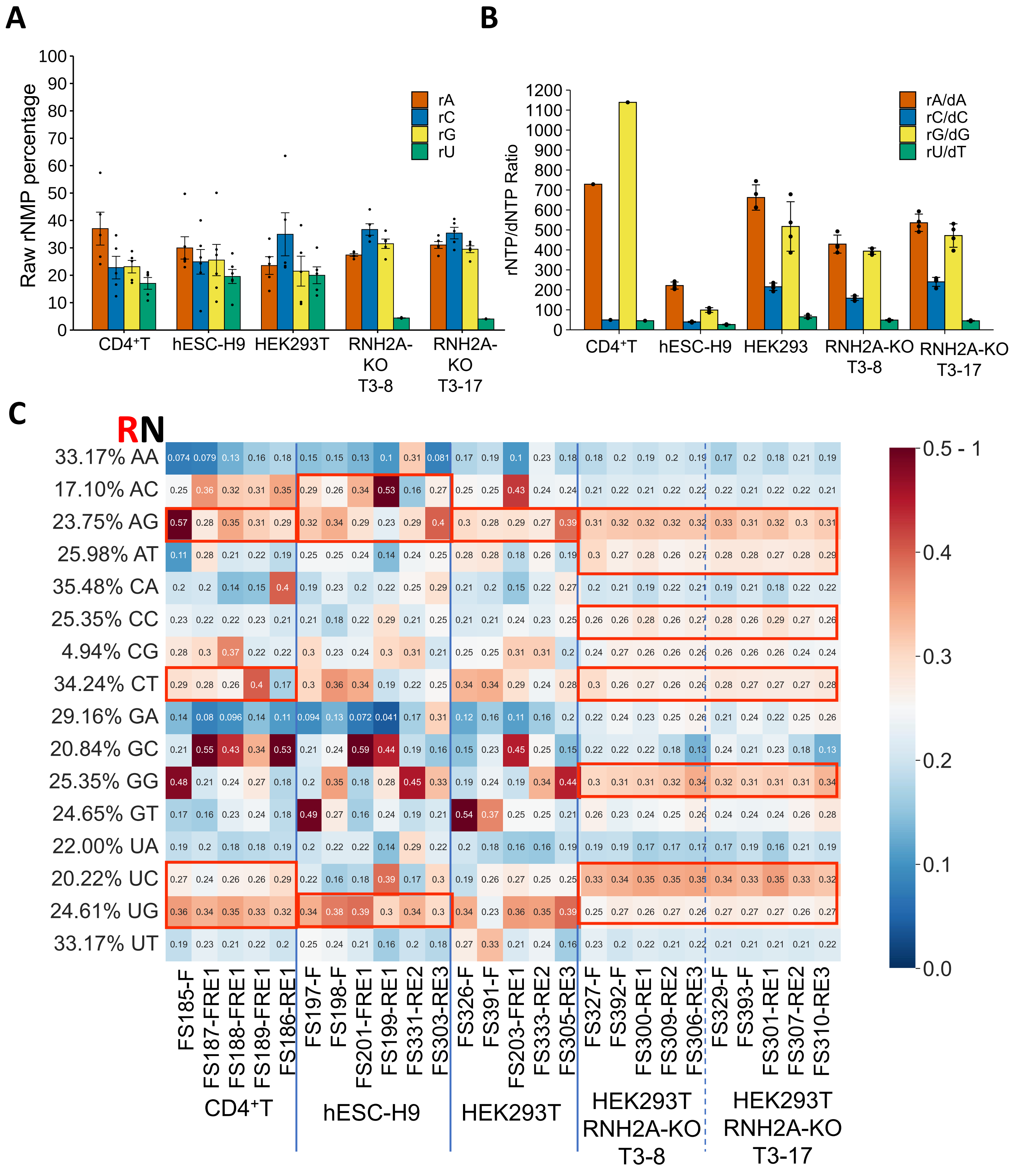

A
Raw rNMP percentage
rA
rC
rG
rU
CD4⁺T
hESC-H9
HEK293T
RNH2A-KO T3-8
RNH2A-KO T3-17
B
rNTP/dNTP Ratio
rA/dA
rC/dC
rG/dG
rU/dT
CD4⁺T
hESC-H9
HEK293
RNH2A-KO T3-8
RNH2A-KO T3-17
C
RN
33.17% AA
17.10% AC
23.75% AG
25.98% AT
35.48% CA
25.35% CC
4.94% CG
34.24% CT
29.16% GA
20.84% GC
25.35% GG
24.65% GT
22.00% UA
20.22% UC
24.61% UG
33.17% UT
FS185-F
FS187-FRE1
FS188-FRE1
FS189-FRE1
FS186-RE1
FS197-F
FS198-F
FS201-FRE1
FS199-RE1
FS331-RE2
FS303-RE3
FS326-F
FS391-F
FS203-FRE1
FS333-RE2
FS305-RE3
FS327-F
FS392-F
FS300-RE1
FS309-RE2
FS306-RE3
FS329-F
FS393-F
FS301-RE1
FS307-RE2
FS310-RE3
CD4⁺T
hESC-H9
HEK293T
HEK293T RNH2A-KO T3-8
HEK293T RNH2A-KO T3-17
0.5 - 1

**fig. S11**

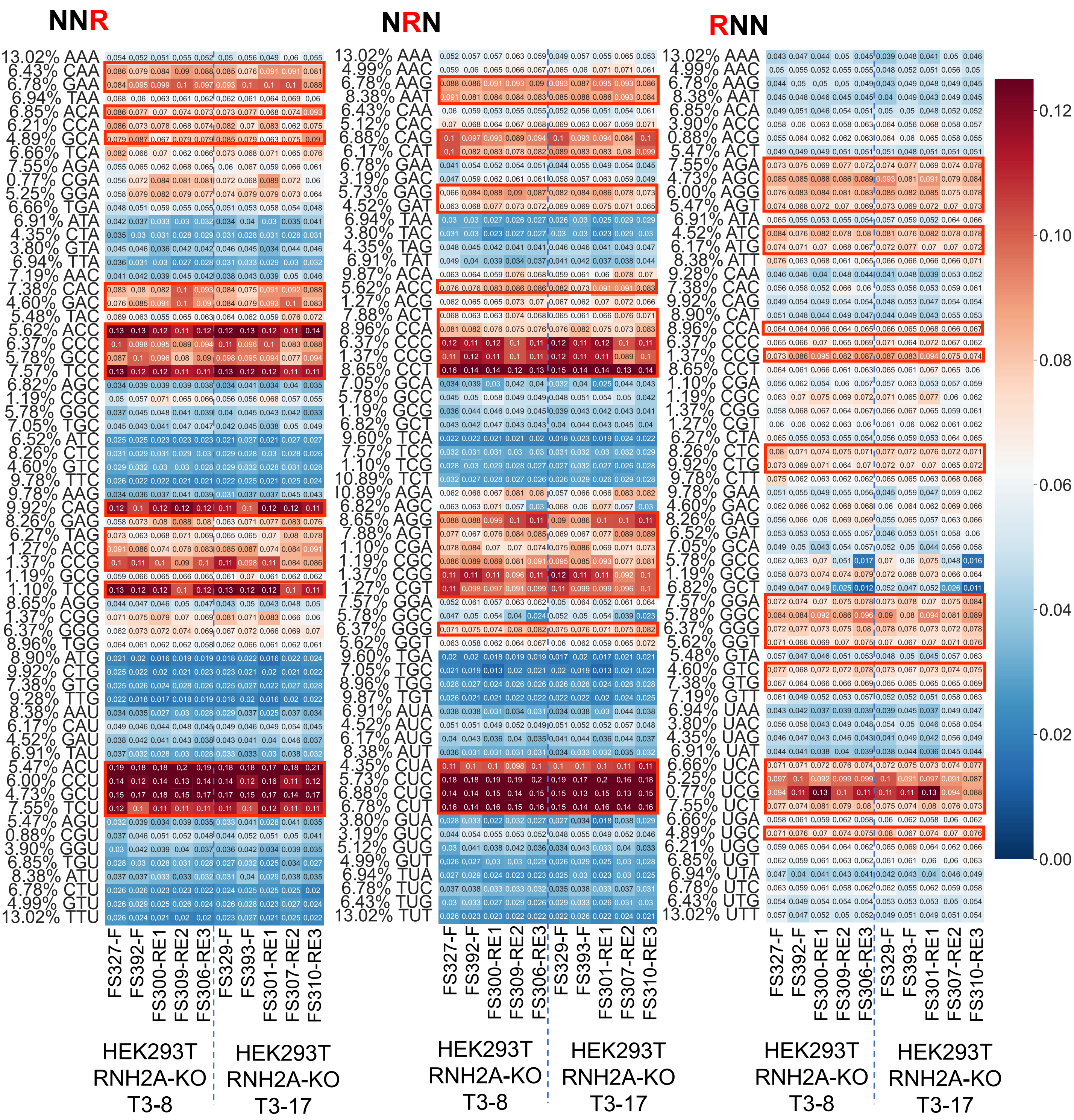

fig. S12

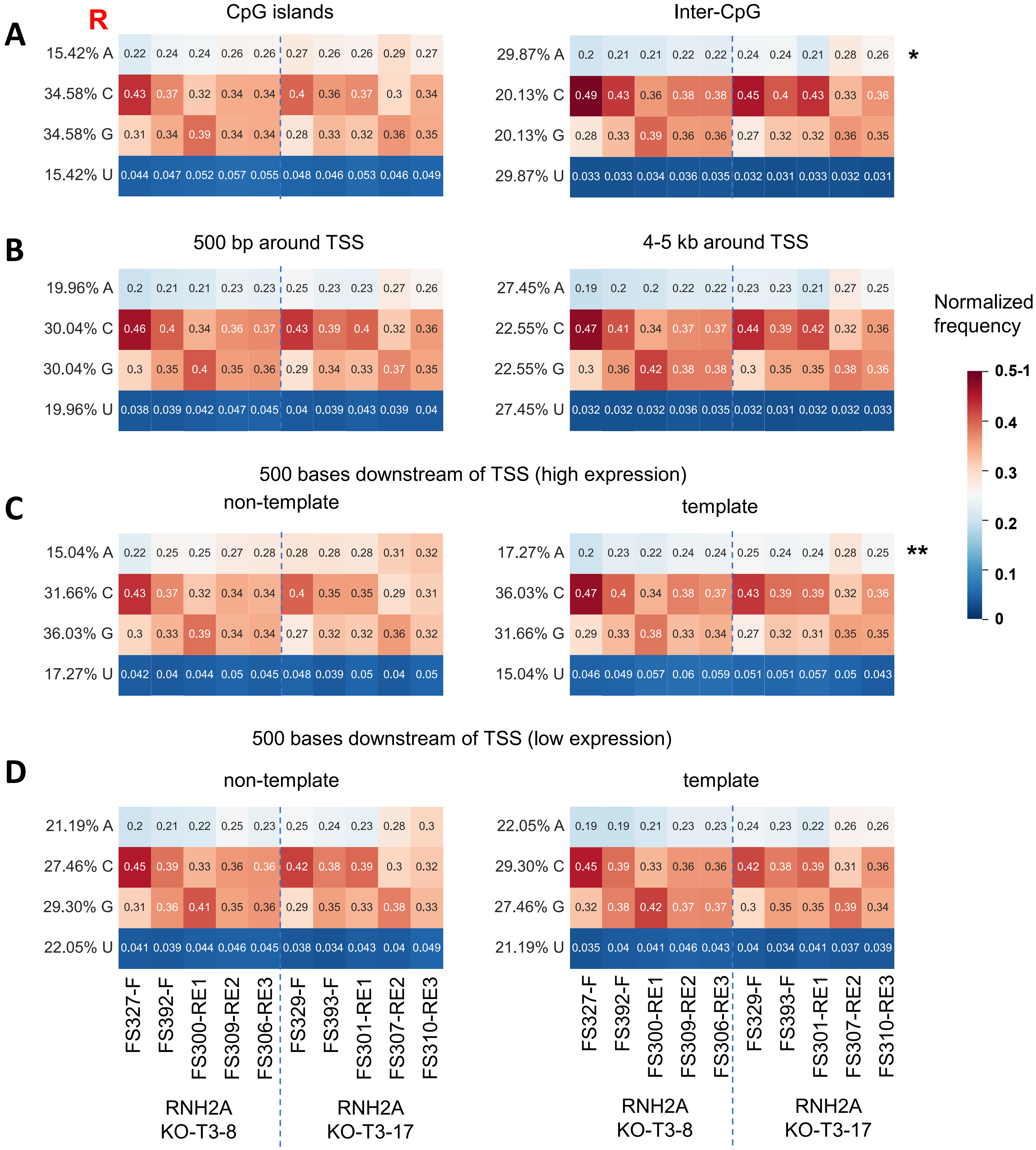

A
R
CpG islands
Inter-CpG
15.42% A
34.58% C
34.58% G
15.42% U
29.87% A
20.13% C
20.13% G
29.87% U
*
B
500 bp around TSS
4-5 kb around TSS
19.96% A
30.04% C
30.04% G
19.96% U
27.45% A
22.55% C
22.55% G
27.45% U
Normalized frequency
0.5-1
0.4
0.3
0.2
0.1
0
500 bases downstream of TSS (high expression)
C
non-template
template
15.04% A
31.66% C
36.03% G
17.27% U
17.27% A
36.03% C
31.66% G
15.04% U
**
500 bases downstream of TSS (low expression)
D
non-template
template
21.19% A
27.46% C
29.30% G
22.05% U
22.05% A
29.30% C
27.46% G
21.19% U
FS327-F
FS392-F
FS300-RE1
FS309-RE2
FS306-RE3
FS329-F
FS393-F
FS301-RE1
FS307-RE2
FS310-RE3
RNH2A KO-T3-8
RNH2A KO-T3-17

fig. S13

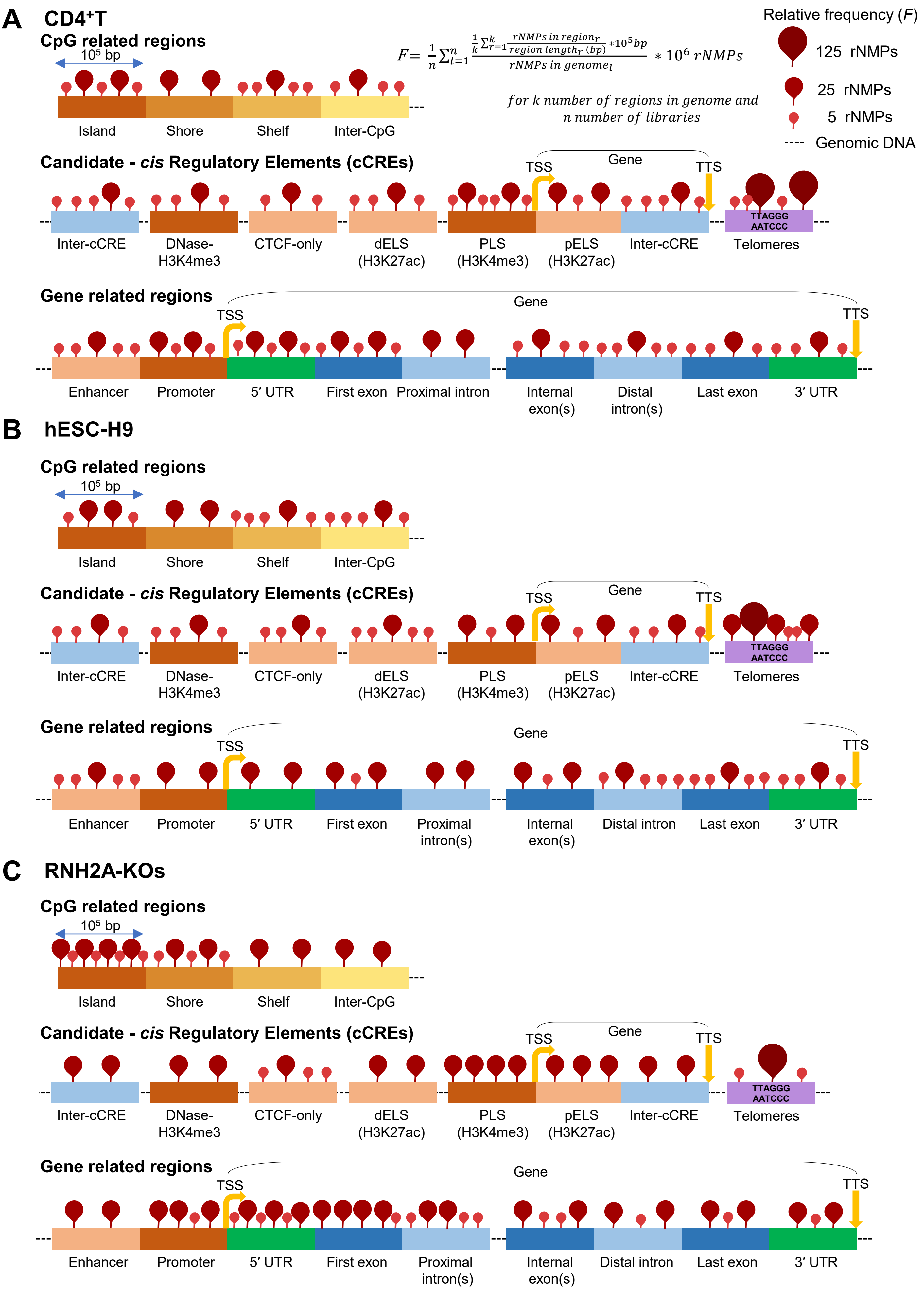

A CD4⁺T
CpG related regions
10⁵ bp
Island
Shore
Shelf
Inter-CpG
$F = \frac{1}{n}\sum_{l=1}^{n} \frac{\frac{1}{k}\sum_{r=1}^{k}\frac{rNMPs\ in\ region_r}{region\ length_r\ (bp)} * 10^5 bp}{rNMPs\ in\ genome_l} * 10^6\ rNMPs$
for k number of regions in genome and n number of libraries
Relative frequency (F)
125 rNMPs
25 rNMPs
5 rNMPs
---- Genomic DNA
Candidate - cis Regulatory Elements (cCREs)
TSS
Gene
TTS
Inter-cCRE
DNase-H3K4me3
CTCF-only
dELS (H3K27ac)
PLS (H3K4me3)
pELS (H3K27ac)
Inter-cCRE
TTAGGG AATCCC
Telomeres
Gene related regions
Gene
TSS
TTS
Enhancer
Promoter
5′ UTR
First exon
Proximal intron
Internal exon(s)
Distal intron(s)
Last exon
3′ UTR
B hESC-H9
CpG related regions
10⁵ bp
Island
Shore
Shelf
Inter-CpG
Candidate - cis Regulatory Elements (cCREs)
TSS
Gene
TTS
Inter-cCRE
DNase-H3K4me3
CTCF-only
dELS (H3K27ac)
PLS (H3K4me3)
pELS (H3K27ac)
Inter-cCRE
TTAGGG AATCCC
Telomeres
Gene related regions
Gene
TSS
TTS
Enhancer
Promoter
5′ UTR
First exon
Proximal intron(s)
Internal exon(s)
Distal intron
Last exon
3′ UTR
C RNH2A-KOs
CpG related regions
10⁵ bp
Island
Shore
Shelf
Inter-CpG
Candidate - cis Regulatory Elements (cCREs)
TSS
Gene
TTS
Inter-cCRE
DNase-H3K4me3
CTCF-only
dELS (H3K27ac)
PLS (H3K4me3)
pELS (H3K27ac)
Inter-cCRE
TTAGGG AATCCC
Telomeres
Gene related regions
Gene
TSS
TTS
Enhancer
Promoter
5′ UTR
First exon
Proximal intron(s)
Internal exon(s)
Distal intron
Last exon
3′ UTR

**fig. S14**

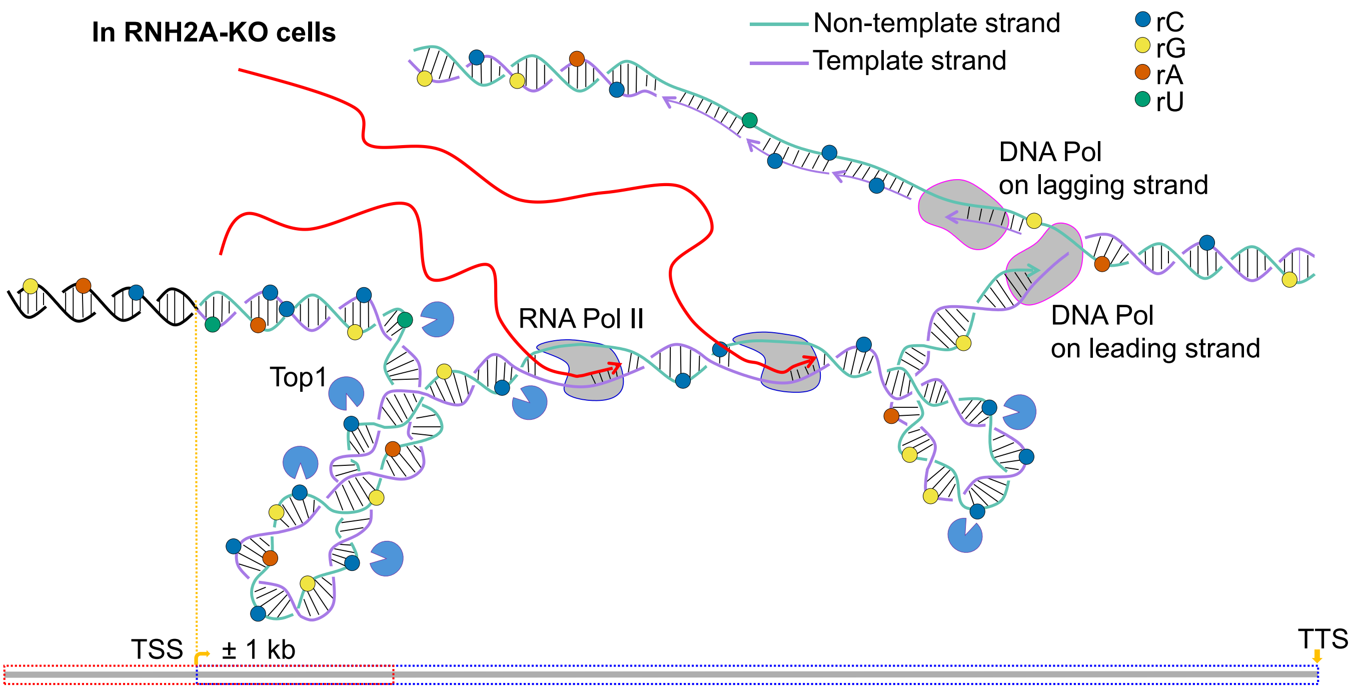
In RNH2A-KO cells
Non-template strand
Template strand
rC
rG
rA
rU
DNA Pol
on lagging strand
DNA Pol
on leading strand
RNA Pol II
Top1
TSS
± 1 kb
TTS
